# From biofilms to birth: Quantitative murburn rationale for hydrated polymer-centred biological transduction, coherence, and evolution of complex life

**DOI:** 10.64898/2026.07.10.737745

**Authors:** Kelath Murali Manoj, Laurent Jaeken, Hirohisa Tamagawa, V L S Prasad Burra

## Abstract

Hydrated extracellular polymeric phases (such as mucus, biofilms, and extracellular matrices) have traditionally been viewed as passive barriers. We complement and extend this view by analysing these systems through the murburn framework and liquid-liquid phase separation (LLPS) biophysics. Using quantitative modeling, we first demonstrate how frothy mucus in amphibian egg-masses enhances oxygen delivery while buffering diffusible reactive species (DRS), leading to improved developmental synchrony. We then model the human cervical mucus system, showing its cycle-dependent transitions between coherent barriers (pregnancy), active transduction media (ovulation), and controlled inflammatory remodeling (labor). Finally, we present thiolated polyglycerol sulfate (dPGS-SH) as a synthetic validation (another group’s recently published work): this rationally designed mucolytic agent recapitulates native mucus’s DRS-modulating properties and shows superior efficacy for addressing cystic fibrosis pathology. With such pan-systemic perspectives, we argue that phase-separated hydrated polymeric matrices represent one of evolution’s most conserved solutions for regulating stochastic murburn chemistry, enabling organisms to exploit oxygen while preserving biological coherence. From biofilms to birth, this framework unifies the physicochemical basis of life’s most fundamental processes.

## 1. Introduction

### 1.1 Classical views of biological organization

Modern biology has traditionally interpreted living systems through deterministic frameworks emphasizing molecular specificity, membrane compartmentalization, enzyme catalysis, receptor-mediated signaling, and genetically encoded regulation (Alberts 2014). Within this paradigm, reactive oxygen species (ROS) and related radicals were long considered undesirable metabolic byproducts associated primarily with oxidative damage and aging (Halliwell and Gutteridge 2015). Accordingly, antioxidant systems were interpreted mainly as defensive mechanisms designed to eliminate such reactive species. However, accumulating evidence over the past several decades has challenged this interpretation. ROS and reactive nitrogen species (RNS) are now known to regulate cellular proliferation, apoptosis, immunity, vascular tone, development, circadian rhythms, and neuronal signaling (Forman et al. 2010). Superoxide, hydrogen peroxide, nitric oxide, hydroxyl radicals, and related diffusible reactive species (DRS) participate in numerous physiological processes across virtually all domains of life. The persistence and evolutionary conservation of such species suggest that biological systems evolved not merely to suppress stochastic redox chemistry, but to regulate and exploit it.

This realization raises a fundamental question: how do living systems maintain coherence amidst continuous stochastic redox activity in aqueous environments?

### 1.2 The murburn perspective

The murburn concept proposes that diffusible reactive species are not incidental byproducts but obligatory and distributed participants in biological transduction and metabolism (Manoj 2018; Manoj and Bazhin 2021; Jaeken & Manoj, 2025). In this framework, oxygen-centered chemistry, stochastic electron transfer, and probabilistic diffusible interactions are intrinsic features of life rather than aberrations requiring complete suppression. This may appear to be in conflict with the aesthetic perceptions of deterministic physiological theories; but it is quite well in line with statistical mechanics, which cannot predict the individual substance characteristics deterministically, but it can predict the mass characteristics (also aligning well with basic ideas of chemico-physics).

The murburn view differs significantly from strictly deterministic “molecular machine” models. Instead of envisioning highly insulated and sequential electron transfer systems alone, murburn theory emphasizes dynamically organized aqueous redox environments wherein diffusible radicals participate in controlled stochastic processes (Manoj and Bazhin 2021). Such an interpretation aligns with the widespread occurrence of ROS-mediated signaling, extracellular radical chemistry, membrane-associated redox dynamics, and distributed physiological responses (Jaeken & Manoj, 2025). Within this perspective, biological organization depends not only on molecular specificity but also on mesoscale physicochemical regulation of diffusion, hydration, electrostatics, and redox probability fields.

### 1.3 The neglected universality of hydrated polymeric systems

An underappreciated but striking feature of life is the near-universal occurrence of hydrated extracellular polymeric phases. These include: mucus, extracellular matrices (ECM), glycocalyces, microbial extracellular polymeric substances (EPS), slime layers, surfactant-associated gels, hydrogel phases, and mucopolysaccharide-rich interfaces.

Such systems occur across bacteria, archaea, algae, fungi, plants, invertebrates, and vertebrates (Flemming and Wingender 2010; Bansil and Turner 2018) Despite major biochemical diversity, these materials converge toward similar physicochemical properties: high hydration, charge density, viscoelasticity, amphiphilicity, diffusion modulation, interfacial localization, and environmental responsiveness.

Conventionally, these systems are interpreted primarily as: protective barriers, lubricants, adhesive layers, hydration-retaining structures, or immune-defense scaffolds (Cone 2009) However, their evolutionary universality, energetic expense, excitation sensitivity, and intimate association with oxygen-handling interfaces suggest deeper physicochemical significance. We propose that hydrated polymeric phases function as mesoscale regulators of stochastic aqueous redox chemistry and thereby contribute fundamentally to biological transduction, coherence preservation, multicellularity, and environmental adaptation.

## 2. Hydrated polymeric phases as a unified biological category

### 2.1 Structural convergence across biology

Hydrated extracellular systems differ biologically yet converge structurally. Vertebrate mucus contains glycoprotein-rich mucins; extracellular matrices are dominated by proteoglycans and glycosaminoglycans; bacterial biofilms contain extracellular polysaccharides, proteins, lipids, and extracellular DNA; glycocalyces form glycoprotein-polysaccharide brushes; slime layers and plant mucilage contain hydrated acidic polymers (Flemming et al. 2016; Frantz et al. 2010). Figure 1 shows a schematic of some muco-polymers and their reaction attributes.

**Figure 1:**
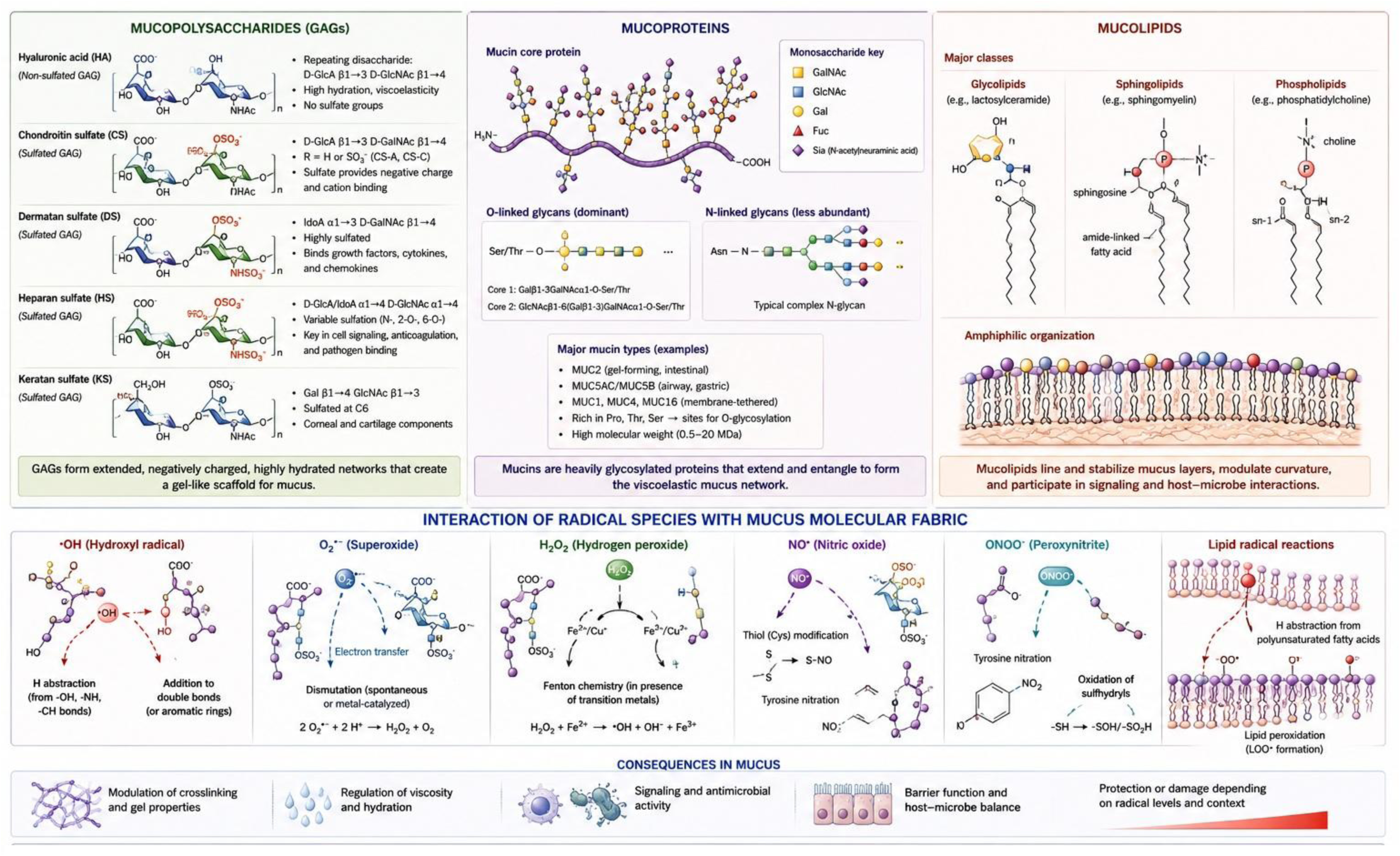
Structural and reaction attributes of muco-polymers.

Representative components include: hyaluronic acid, heparan sulfate, chondroitin sulfate, alginates, sialylated mucins, amphiphilic lipids, extracellular DNA-protein complexes, and sulfated glycoproteins. Despite biochemical variation, these systems repeatedly converge toward:hydrated polymeric networks, polyelectrolyte architectures, amphiphilic organization, viscoelastic phase behavior, and diffusion-regulating interfaces. This convergence strongly suggests common physicochemical utility. Accordingly, mucus, ECM, glycocalyces, EPS matrices, slime layers, and hydrogels may be viewed collectively as hydrated polyelectrolyte-interfacial matrices specialized for regulating aqueous physicochemical interactions.

### 2.2 Physicochemical hallmarks

#### Hydration

Hydrogels may contain 70–99% water by mass (Peppas et al. 2006). This creates structured aqueous microdomains that influence: proton mobility, ionic partitioning, oxygen diffusion, radical propagation, and dielectric behavior. Due to the abundance and universal existence, the role of water for life is underestimated though everyone knows it is essential for life. It is usually considered merely as a solvent. Water itself is central to hydroxyl radical chemistry, proton-coupled electron transfer, superoxide equilibria, and peroxide dynamics (Ball 2008).

#### Polyelectrolyte behavior

Many extracellular polymers contain: sulfates, carboxylates, phosphates, protonatable amines, which generate electrostatic fields capable of modulating ion distributions and electron transfer probabilities. Acidic glycosaminoglycans such as heparan sulfate and hyaluronic acid are particularly important in this regard (Frantz et al. 2010).

#### Amphiphilicity

Mucolipids and surfactant-associated systems possess hydrophobic and hydrophilic microdomains. Such compartmentalization affects: oxygen partitioning, membrane interaction, radical segregation, and selective permeability. Pulmonary surfactant systems exemplify such amphiphilic organization (Possmayer et al. 2001; Olmeda et al. 2010).

#### Dynamic phase behavior

Hydrated polymeric systems are not static structures. They exhibit: swelling, dehydration, shear thinning, sol-gel transitions, and ion-responsive rearrangements. These dynamic phase transitions allow adaptive responses to environmental and metabolic perturbations. Collectively, these observations suggest that hydrated extracellular matrices are active physicochemical environments rather than passive structural materials.

The elucidation of the structure of biological molecules provides us scientifically quite meaningful information. However, features based on X-ray diffraction of crystalized molecules or even in cryo-electron microscopy, we can only get a static picture. Therefore, the physiological “movements” of such structures are left to imagination, as electrical dynamics of biological activities emerges when they are in a wet or fluidized state. Figure 2 shows some properties of the hydrated biopolymers.

**Figure 2:**
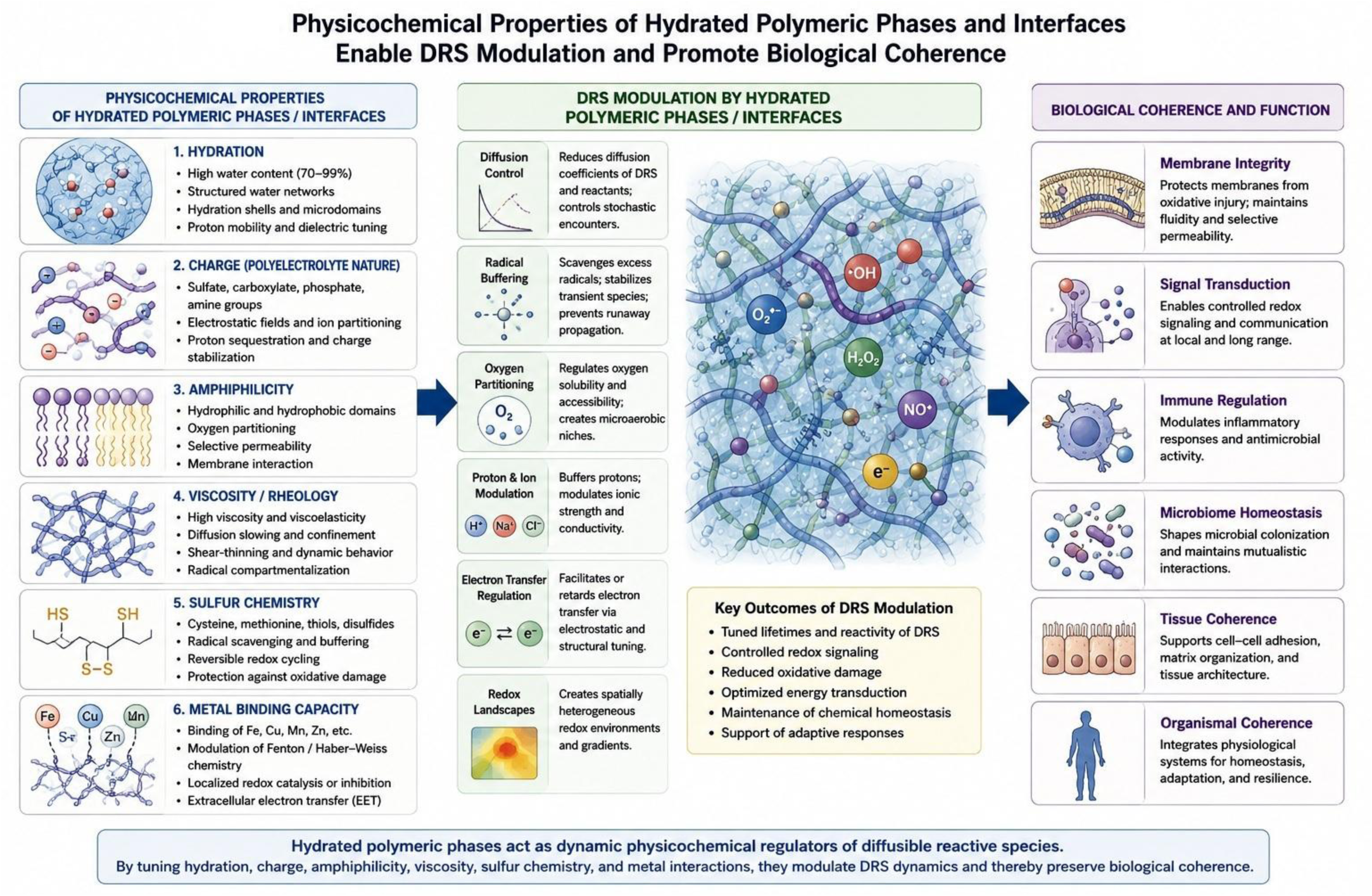
Properties of hydrated biological polymers.

### 2.3 Liquid–liquid phase separation and coacervate formation in hydrated polymeric phases

A growing body of biophysical research has established liquid–liquid phase separation (LLPS) and coacervate formation as fundamental mechanisms for organizing biomolecules into dense, membrane-less liquid phases embedded within more dilute aqueous environments. In such systems, multivalent interactions among charged and amphiphilic polymers drive spontaneous demixing into a polymer-rich dense phase and a surrounding dilute phase, often followed by a gradual transition of the dense phase into viscoelastic hydrogel networks (Xie et al. 2025; Emenecker et al. 2021). Several extracellular matrices appear to assemble via LLPS/coacervation. For example, tropoelastin-like polymers can first form liquid coacervate droplets that subsequently crosslink into elastomeric ECM-like networks; these systems reproduce many mechanical and signaling properties of native extracellular matrices. Hydrated mucosal and glycosaminoglycan-rich interfaces share the same core biophysical ingredients (high charge density, amphiphilic segmentation, and strong sensitivity to pH, ionic strength, and temperature) making LLPS/coacervation a plausible route for their initial assembly and dynamic restructuring (Xie et al. 2025)

Within murburn framework, LLPS adds an additional level of mesoscale regulation: dense coacervate or condensate phases and the surrounding dilute phases can differ markedly in local concentrations of oxygen, diffusible reactive species (DRS), ions, and redox-active cofactors. This micro-heterogeneity provides a natural physical basis for the “redox microdomains” and stochastic encounter landscapes invoked in murburn concept, without requiring rigid membrane boundaries.

### 2.4. Physicochemical basis for DRS modulation

#### Water structuring and redox microdomains

Water is not an inert solvent in biological systems. Hydrogen-bond organization, hydration shell dynamics, proton relay pathways, and ionic microdomains strongly influence radical behavior and electron transfer processes (Ball 2008). Hydrated polymeric matrices modify: hydrogen-bond structure, proton conductivity, local dielectric properties, and diffusion kinetics. Such environments therefore influence: hydroxyl radical lifetimes, superoxide mobility, peroxide stability, and proton-coupled electron transfer reactions. Hydrogels and mucus systems may thus function as controlled aqueous reaction environments capable of shaping stochastic redox chemistry.

It is well-known that mass characteristics of water molecules are physicochemically significantly different from those of a single water molecule. The most familiar example is surface tension of water caused by the formation of hydrogens bonds among water molecules. Another example: the relative permittivity of highly oriented water molecules are known to be significantly low. Therefore, water is not just a solvent, and the works of Pollack (Pollack 2001; Pollack and Reitz 2001) has theorized and characterized many states of physiological water.

#### Diffusion control and radical compartmentalization

Reactive radicals are diffusion-limited species. Therefore, viscosity and confinement profoundly affect radical propagation and encounter probabilities. Hydrated polymeric phases: slow diffusion, compartmentalize reactants, localize signaling ROS, stabilize transient intermediates, and prevent runaway chain reactions. Biofilm EPS matrices, for example, strongly regulate oxygen penetration and extracellular diffusion fields (Stewart and Franklin 2008). Such regulation may preserve coherence while permitting controlled stochastic signaling.

#### Electrostatics and dielectric effects

Extracellular polymers generate electrostatic organization through charged functional groups including: sulfate moieties, carboxylates, phosphates, and protonatable amines. These charges affect: proton sequestration, ion partitioning, charge stabilization, electron transfer kinetics, and local dielectric constants. Electrostatic organization strongly influences radical collision frequencies and DRS lifetimes. Thus, hydrated polymeric systems likely regulate the probabilistic landscape of redox interactions.

#### Amphiphilic organization and oxygen partitioning

Hydrophobic microdomains preferentially partition: oxygen, lipid radicals, xenobiotics, and hydrophobic metabolites. Hydrophilic domains regulate: ion mobility, proton diffusion, ROS stabilization, and hydration-dependent charge dissipation. Pulmonary surfactant gels provide especially important examples of oxygen-partitioning amphiphilic interfaces (Possmayer et al. 2001). Such organization spatially compartmentalizes redox chemistry.

#### Sulfur chemistry and redox buffering

Mucoproteins and extracellular proteins often contain: cysteine, methionine, thiols, and disulfide bonds. Sulfur chemistry plays central roles in oxidative buffering and reversible redox regulation (Winterbourn and Hampton 2008). Thiol-containing systems: quench radicals, buffer electron flux, stabilize oxidative perturbations, and participate in reversible oxidation-reduction cycles. Such chemistry may be especially important in fluctuating oxidative environments.

#### Transition metals and distributed redox landscapes

Extracellular polymers frequently bind metal ions/complexes of: Fe, Cu, Mn, Zn, etc. through electrostatic and coordination interactions. These metals influence: Fenton chemistry, peroxide decomposition, extracellular electron transfer, and ROS generation (all of which collectively fall under murburn). Biofilm matrices may therefore function as distributed extracellular redox landscapes (Flemming et al. 2016)

#### Dynamic phase adaptation

Hydrated polymeric systems respond dynamically to: pH, oxygen tension, ionic strength, inflammation, metabolic activation, and mechanical stress. Sol-gel transitions and rheological rearrangements may thereby regulate mesoscale stochastic redox behavior adaptively. These systems therefore function not merely as structural materials but as dynamic redox-responsive interfaces.

### 2.5 Mesoscale redox organization as a biological principle

Classical biology largely emphasizes molecular specificity. However, many biological processes emerge at mesoscale dimensions between individual molecules and tissues. Hydrated polymeric phases regulate: diffusion fields, oxygen gradients, proton mobility, stochastic encounter rates, radical propagation, and interfacial electron transfer. Such regulation is indispensable for biological coherence in the murburn view, as DRS are deemed essential for life-sustenance. Figure 3 captures some elements of hydrated biopolymers serving as mesoscale organizers.

**Figure 3:**
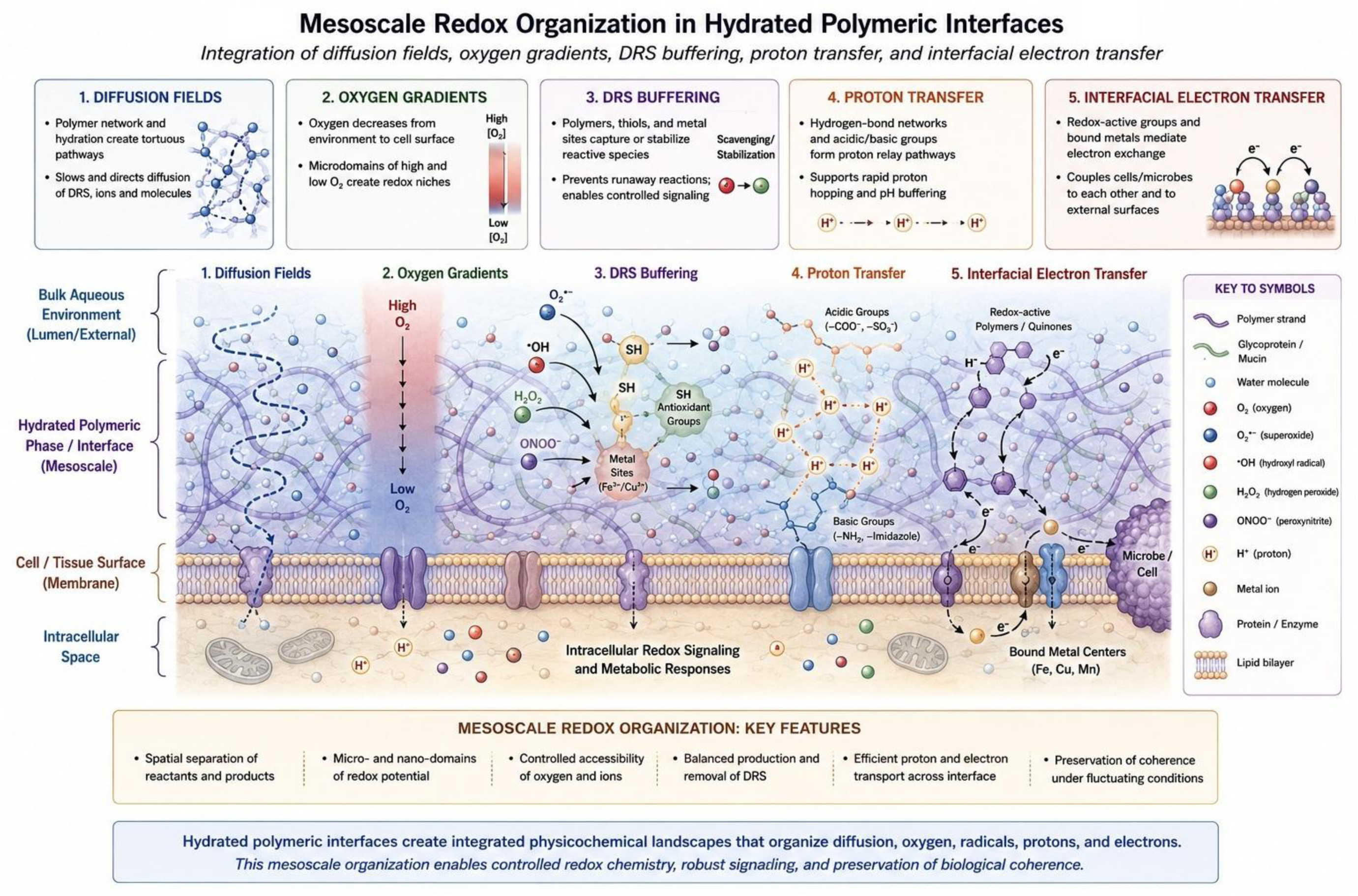
Hydrated biopolymers as mesoscale organizers.

An important class of mesoscale organizing phenomena now recognized in cell and matrix biology is liquid–liquid phase separation, giving rise to biomolecular condensates or coacervates that concentrate selected macromolecules into dense, yet liquid, compartments without membranes. These condensates have distinct viscosity, dielectric properties, and diffusion coefficients compared to the surrounding aqueous phase, and can thereby tune local reaction rates and radical encounter probabilities. When hydrated polymeric phases undergo LLPS or coacervate formation, the resulting microdomains offer a natural physical substrate for murburn-type stochastic redox chemistry: dense phases can act as sites of enhanced or buffered DRS activity, while dilute phases mediate long-range diffusion and communication (Wang et al. 2024; Emenecker et al. 2021; Xie et al. 2025).

#### Distributed biological transduction

Biological signaling may not be exclusively receptor-centric. Distributed probabilistic interactions involving: DRS, ionic fields, hydration structures, and diffusional gradients,likely contribute substantially to physiological transduction. Examples include: ROS-mediated signaling, extracellular nitric oxide dynamics, inflammatory oxidative bursts, and microbial redox communication. Hydrated extracellular interfaces provide physicochemical environments capable of organizing such distributed interactions.

#### Coherence preservation

Biological systems continuously experience stochastic excitation arising from: metabolism, oxygen chemistry, membrane perturbation, environmental fluctuations, and immune activation. Without damping mechanisms, uncontrolled radical propagation could destabilize membranes, proteins, and collective physiological behavior. Hydrated polymeric interfaces may therefore function as:stochastic dampeners, coherence-preserving media, distributed redox moderators, and transduction-organizing environments. This interpretation reframes mucus, ECM, glycocalyces, and EPS matrices as active thermodynamic participants in biological organization (and not just mechanical frameworks!).

#### Murburn and mesoscale organization

The murburn framework naturally aligns with mesoscale regulation because diffusible radicals inherently depend upon: diffusion probabilities, aqueous structuring, electrostatic partitioning, oxygen accessibility, and stochastic encounter dynamics. Accordingly, mesoscale extracellular organization may be as important to biological function as molecular specificity itself.

The integration of liquid-liquid phase separation (LLPS) provides the definitive thermodynamic mechanism for the murburn concept. The interfacial electric fields (IEFs) generated at the boundaries of phase-separated coacervates may possess sufficient potential to spontaneously drive or facilitate some redox reactions. Furthermore, the unique solvation microenvironment within these condensates promotes spontaneous proton-coupled electron transfer, which may generate or modulate diffusible reactive species parallel to murzymatic routes. By physically compartmentalizing these reactants, the dense coacervate phase regulates and restricts the diffusion of these stochastic radicals, protecting the broader cellular machinery from uncontrolled oxidative damage. Thus, the phase-separated state of biological matter inherently acts as a simple chemical engine (SCE) pre-programmed to support murburn dynamics.

### 2.6. Evolutionary origins of hydrated redox interfaces

#### Early aqueous redox environments

Primitive Earth likely contained: UV-driven radical chemistry, metal-rich waters, fluctuating oxygen levels, redox turbulence, and stochastic electron transfer processes. Under such conditions, hydrated polymeric compartments capable of: retaining water, slowing diffusion, stabilizing ions, buffering radicals, organizing electrochemical microdomains, etc. would possess major selective advantages.

#### Primitive hydrogels before sophisticated membranes?

It is plausible that protocellular evolution initially involved: hydrated gels, electrostatic microdomains, primitive hydrogel compartments, and stochastic redox niches; before emergence of highly sophisticated lipid membrane systems and the associated interfacial murburn chemistry. Hydrogel-like matrices may therefore represent primordial redox-buffering structures. Such a scenario aligns with hypotheses emphasizing the importance of mineral-organic interfaces, colloidal phases, and prebiotic compartmentalization in origin-of-life chemistry (Deamer and Dworkin 2005).

#### Biofilms and collective redox coherence

Microbial biofilms provide compelling examples of collective extracellular redox organization. EPS matrices regulate oxygen penetration, stabilize communal metabolism, facilitate extracellular electron transfer, protect against oxidative stress, and preserve hydration. Biofilms therefore function not merely as adhesive communities but as distributed redox-coherence systems (Flemming and Wingender 2010)

Recent findings demonstrate that the formation of the EPS matrix in biofilms—such as those of *Escherichia coli*—is fundamentally driven by the liquid-liquid phase separation of nucleoid-associated proteins, extracellular DNA, and lipopolysaccharides. This resulting coacervate is highly redox-active. The phase-separated EPS matrix serves as a hotspot for extracellular electron transfer, utilizing both non-enzymatic components (like flavins) and redox proteins to move electrons to molecular oxygen, thereby generating superoxide. This EPS-driven murburn radical generation not only maintains the structural and thermodynamic integrity of the biofilm but also actively drives massive biogeochemical cycles, such as manganese oxidation, far more efficiently than free-swimming microbes.

#### Evolutionary continuity toward multicellularity

An evolutionary continuum can be envisioned linking: slime layers, glycocalyces, extracellular hydrogels, ECM systems, and vertebrate mucosal interfaces (Figure 4). Hydrated extracellular polymeric systems may have enabled: tissue coherence, smooth intercellular networking and integration, oxygen adaptation, developmental coordination, and multicellular organization. The recurrent emergence of such systems across biology suggests a universal physicochemical necessity associated with managing stochastic aqueous redox dynamics.

**Figure 4:**
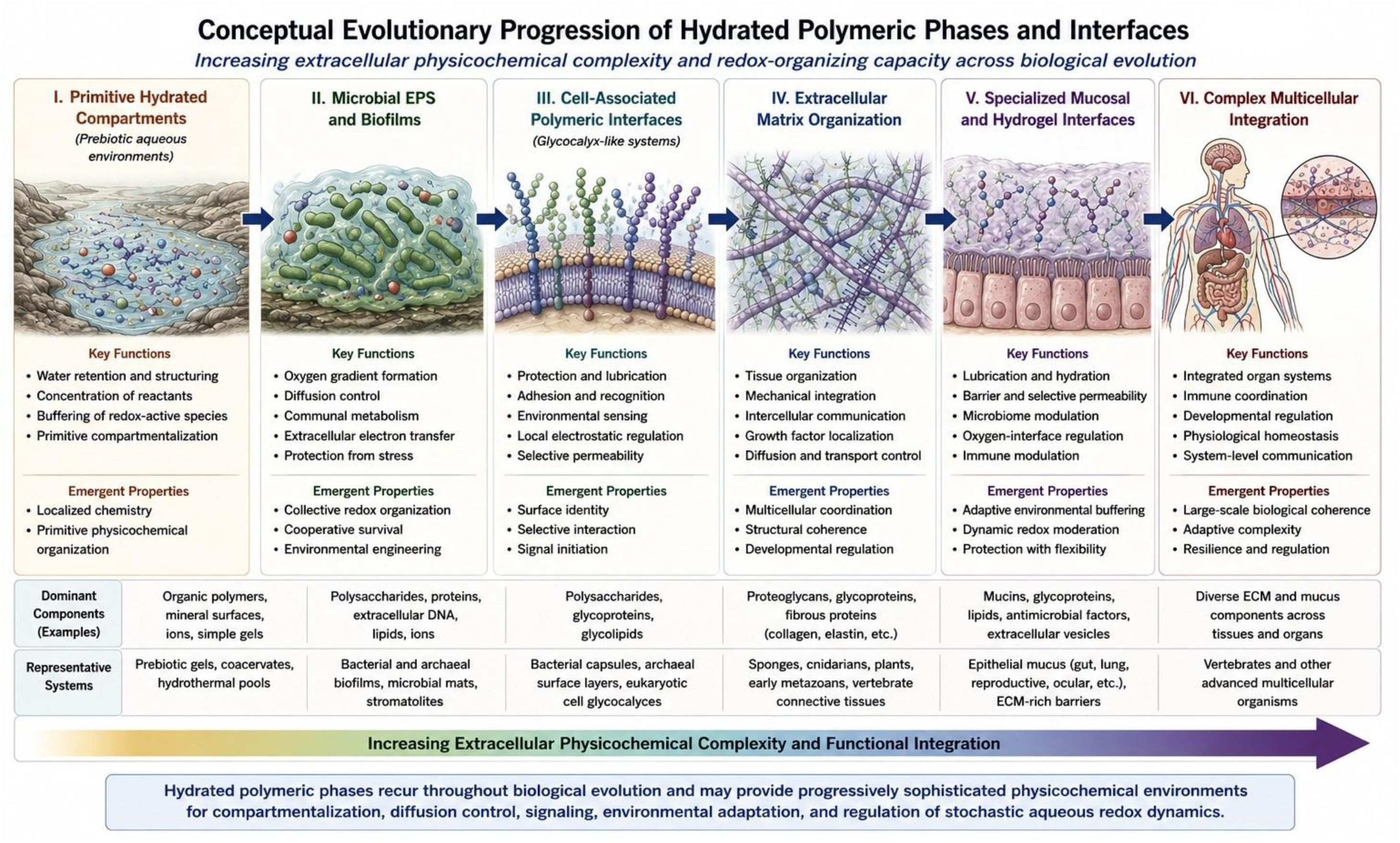
Conceptual evolutionary progression illustrating increasing complexity and specialization of hydrated extracellular polymeric phases. The sequence depicts functional and physicochemical continuity rather than a strict phylogenetic chronology. Across biological evolution, hydrated polymeric interfaces appear repeatedly as solutions for compartmentalization, diffusion control, environmental adaptation, intercellular communication, and potentially the moderation/modulation of stochastic aqueous redox processes.

## 3. Toward a murburn framework for biological coherence

The recurrent emergence of hydrated extracellular polymeric systems across biology suggests a universal physicochemical necessity. Life may not have evolved primarily to eliminate stochastic redox chemistry. Such processes were/are too advantageous, were it only by their exergonic character. Rather, living systems appear to have evolved mechanisms capable of: regulating, compartmentalizing, buffering, exploiting, and stabilizing stochastic aqueous redox interactions. Within this framework: mucus, extracellular matrices, glycocalyces, biofilms, extracellular polymeric substances, and hydrogel phases, become active transduction environments rather than passive structural materials. Given the imperative nature and scope of murburn, then such an outcome and evolutionary disposition can be easily explained.

### Biological coherence in stochastic systems

Biological systems continuously operate far from thermodynamic equilibrium. Metabolism, oxygen chemistry, ionic fluxes, membrane perturbations, and environmental fluctuations constantly generate stochastic excitation. Complete suppression of diffusible reactive species would likely be incompatible with: signaling, immunity, development, adaptation, and oxygenic metabolism itself. Instead, living systems appear to preserve coherence through dynamic regulation of stochasticity. Hydrated polymeric phases are uniquely suited for such regulation because they simultaneously influence: diffusion, hydration, electrostatics, oxygen partitioning, proton mobility, and radical stabilization.

### Hydrated polymeric phases as transduction media

Conventionally, extracellular matrices and mucus are interpreted as: scaffolds, lubricants, barriers, or hydration-retaining structures. However, their physicochemical properties strongly suggest more active roles. Hydrated extracellular phases may function as: distributed transduction environments, mesoscale redox organizers, adaptive damping systems, and coherence-preserving interfaces. They have the properties of a colloid. A colloid can be stimulated by pH, temperature, chemical and biochemical agents, solvents and ions, electrical field, mechanical stress and electromagnetic radiations, and respond to these by: changing reaction rates and adsorptive properties including probabilistic interactions involving DRS, ions, hydration structures, oxygen gradients, and interfacial electron transfer, phase transformation, change shape, shrink and swell, hereby changing all physicochemical parameters at once and in a concerted, coherent way (Minkoff and Damadian 1976) alter permeability, surface properties, mechanical aspects, electrical signalling, electromagnetic radiation and optical properties (Pollack 2001). Since biological extracellular polymeric hydrogels are asymmetric structures, they possess piezo-electric and thermo-electric properties (as the cytoskeleton has).

The colloidal behaviour and dynamic responsiveness of hydrated polymeric phases are closely related to liquid–liquid phase separation phenomena that generate biomolecular condensates and coacervates. In many intracellular systems, LLPS has been shown to produce dense liquid droplets that concentrate enzymes, nucleic acids, and signaling factors into membrane-less organelles with distinct redox and diffusion properties. Extracellular matrices and tropoelastin-based hydrogels can similarly arise from LLPS/coacervate assembly followed by phase transition into viscoelastic networks. Recognizing LLPS as a common mechanism for both internal condensates and external hydrated polymeric interfaces provides a natural physical underpinning for murburn concept: in all these cases, stochastic DRS, ions, and electrons move through structured aqueous environments shaped by phase separation rather than rigid walls. The same basic physics—multivalency, charge patterning, amphiphilicity, and weak interactions—thus generates mesoscale compartments that support controlled stochastic redox chemistry across the spectrum from biofilms to cervical mucus and intracellular condensates (Wang et al. 2021; Heinmöller et al. 2000; Xie et al. 2025; Emenecker et al. 2021).

### Evolutionary necessity of DRS-modulation

The universal recurrence of hydrated extracellular polymeric phases across biology strongly suggests that life required mechanisms capable of managing stochastic aqueous redox chemistry. Accordingly, life may not have evolved to eliminate DRS. Rather, evolution selected systems capable of stabilizing and exploiting them. This interpretation provides a coherent framework linking: biofilms, slime layers, glycocalyces, mucosal systems, extracellular matrices, and multicellular organization.

### A unified murburn interpretation

The murburn framework offers a plausible integrative explanation for: extracellular hydrogels, distributed redox signaling, oxygen-handling interfaces, collective microbial coherence, excitation-linked mucus secretion, and multicellular integration. Hydrated polymeric phases may therefore represent one of the most fundamental physicochemical technologies evolved by life for organizing stochastic aqueous redox dynamics. Their properties are not so different from those of cytoplasm. This is not so surprising. In early pre-biotic coacervates internal structure and surface structure were not yet separated by a membrane and formed one continuity, that of a hydrogel.

### Falsifiability and predictions of the contextual murburn perception

Prediction 1: Hydrogels with greater charge density should display altered DRS lifetimes. Prediction 2: Mucins with higher sulfation should show enhanced radical buffering capacity. Prediction 3: Artificial mucus analogues should influence ROS diffusion coefficients. Prediction 4: Loss of glycocalyx integrity should measurably alter extracellular redox profiles. Prediction 5: Biofilm EPS disruption should increase redox instability before affecting growth.

The predictions outlined above are designed to be experimentally testable and, in principle, falsifiable; a critical requirement for any scientific framework. For instance, Prediction 1 can be directly tested using synthetic hydrogels with systematically varied sulfation or carboxylation levels, combined with electron paramagnetic resonance (EPR) spectroscopy to quantify radical lifetimes. Prediction 3 can be evaluated using fluorescence recovery after photobleaching (FRAP) or electrochemical microsensors in mucus-mimetic hydrogels. Similarly, Prediction 5 can be tested using EPS-degrading enzymes coupled with redox-sensitive fluorescent probes. These experiments would provide direct evidence for or against the murburn interpretation. Failure to observe the predicted correlations (for example, no change in DRS lifetimes with charge density, or no increase in redox instability upon EPS disruption) would challenge the framework and necessitate revision. This explicit falsifiability distinguishes the murburn-LLPS perspective from purely descriptive or post-hoc explanations.

## 4. Quantitative and qualitative illustrations of murburn-LLPS functionalisms

### 4.1 Mucus-froth embedding in frog egg-embryonic growth

Here, we demonstrate how frog eggs deposited in hydrated polymer (frothy-mucus) aids the embryo bulk growth, using plain water as a comparative control interstitial medium/phase. Under simple conditions (in both systems- the control water and the test mucus-froth), the concentration of DRS in milieu is directly correlated with the concentration of DRS. This is demonstrated in Figure 5, and further elaborated by the results obtained from the MATLAB codes given in Supplementary Information, Amphibian code 1 (for Figure 5 below) and code2 (for the four figures therein with the SI, after code 3). The crux of murburn is that the utility of oxygen lies in its ability to generate DRS.

**Figure 5:**
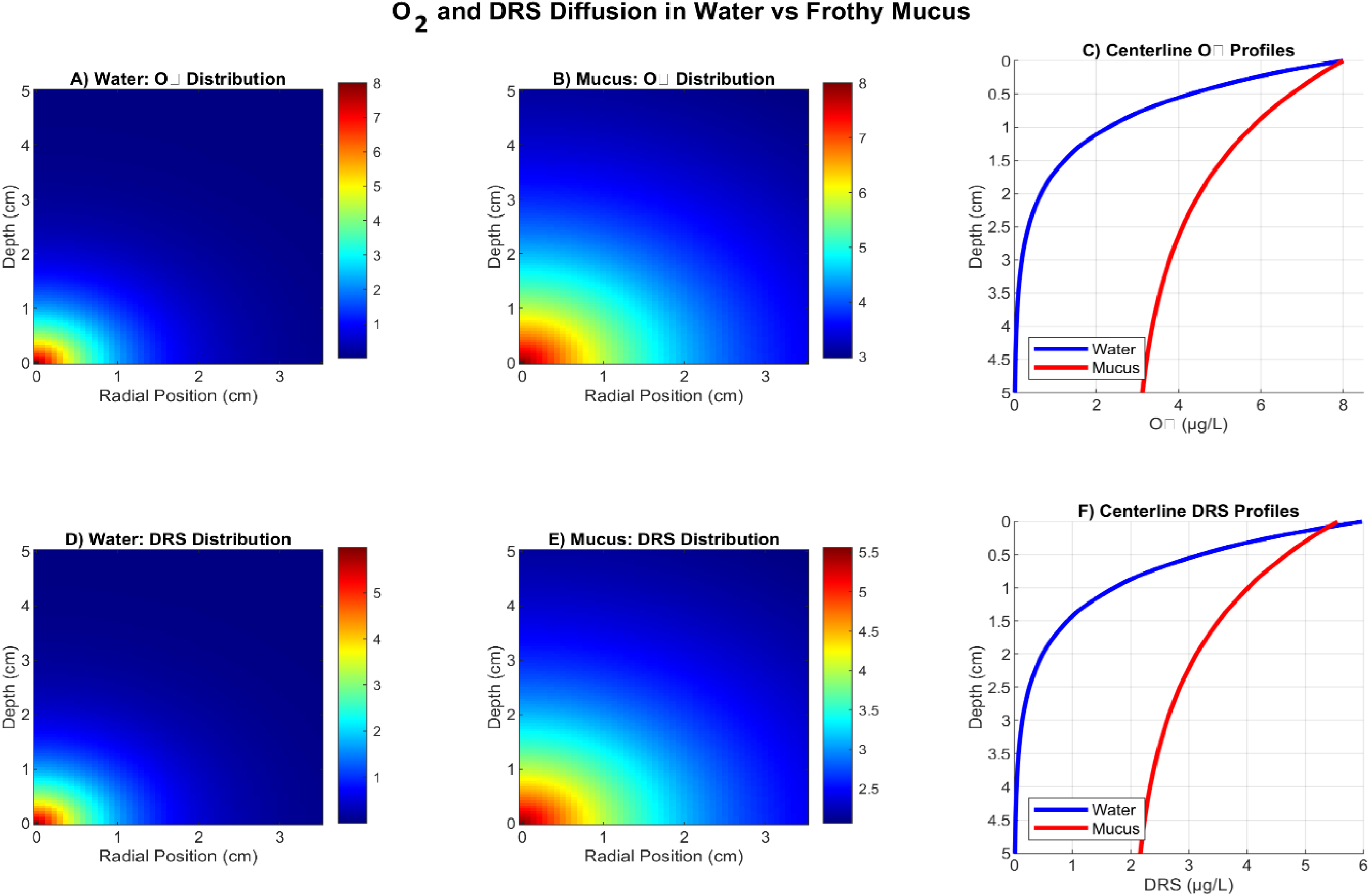
Oxygen profile directly correlates with DRS in milieu.

The other systemic/biological assumptions employed in the model are:

1. Embryo physiology: Frog embryos (*Lechriodus fletcheri*) have an average oxygen consumption rate of 1.0×10⁻⁴ µg O₂ embryo⁻¹ h⁻¹, which is maintained relatively constant throughout development.
2. Oxygen diffusion: The system follows Fick’s First Law of Diffusion, with oxygen moving from areas of high concentration (ambient air-saturated water) to areas of low concentration (consuming embryos).
3. Mucus gel properties: The frothy mucus matrix behaves as a homogeneous hydrogel with reduced diffusivity (25% of water) due to its polymeric structure, but contains air bubbles that serve as oxygen reservoirs.
4. Bubble fraction: Approximately 35% of the frothy mucus volume is occupied by air bubbles, which significantly enhances oxygen availability by creating a gas-phase reservoir.
5. Embryo communication: Embryos within the clutch communicate via oxygen gradients (and DRS thereof), where O₂ concentration at one embryo influences neighboring embryos within a 0.5 cm radius through diffusive coupling.
6. Michaelis-Menten kinetics: Oxygen uptake by embryos follows Michaelis-Menten kinetics with a half-saturation constant (K_m_) of 1.5×10⁻⁶ g/cm³, representing the O₂ concentration at which uptake is half-maximal.
7. Growth model: Embryo growth follows a saturable function of oxygen availability, with a maximum growth rate of 0.08 mm/h and a half-saturation constant of 1.0×10⁻⁶ g/cm³.
8. Spherical symmetry: The egg mass is approximated as a spherical structure with embryos distributed uniformly throughout.

The various parameters/values taken are given in Table 1 and the summary of quantitative outcomes are given in Table 2.

**Table 1:** Parameters/value used for our MATLAB model.

| Parameter | Symbol | Value | Units | Justification |
| --- | --- | --- | --- | --- |
| **Embryo Parameters** |  |  |  |  |
| **Number of embryos** | N | 318 | embryos | Based on *Lechriodus fletcheri* clutch size |
| **Egg radius** | r_egg | 0.09 | cm (0.9 mm) | Typical frog egg diameter |
| **O<sub>2</sub> consumption** | Vo<sub>2</sub> | $1.0 \times 10^{-4}$ | $\mu\text{g O}_2 \text{ embryo}^{-1} \text{ h}^{-1}$ | Estimated from amphibian embryo data |
| **Oxygen Diffusivity** |  |  |  |  |
| In water | D<sub>water</sub> | 2.4×10<sup>-5</sup> | cm<sup>2</sup>/s | Standard literature value at 25°C |
| In mucus gel | D<sub>gel</sub> | 6.0×10<sup>-6</sup> | cm<sup>2</sup>/s | Assumed 25% of water diffusivity |
| **Oxygen Concentrations** |  |  |  |  |
| Saturated O<sub>2</sub> | C<sub>saturatio</sub><br><sub>n</sub> | 8.0×10<sup>-6</sup> | g/cm<sup>3</sup> (8 mg/L) | Air-saturated water at 25°C |
| Ambient O<sub>2</sub> | C<sub>ambient</sub> | 8.0×10<sup>-6</sup> | g/cm<sup>3</sup> | Assumed fully saturated |
| **Mass Geometry** |  |  |  |  |
| Mass radius | R<sub>mass</sub> | 3.5 | cm | Approximate for 318 eggs |
| Mass depth | D<sub>mass</sub> | 5 | cm | Maximum depth of egg mass |
| Bubble fraction | φ<sub>bubble</sub> | 0.35 | dimensionless | Estimated from frothy mucus observations |
| **Boundary Layers** |  |  |  |  |
| Water DBL | δ<sub>water</sub> | 0.2 | cm (2 mm) | Typical diffusion boundary layer in water |
| Mucus film | δ<sub>gel</sub> | 0.008 | cm (0.08 mm) | Thin mucus layer surrounding each egg |
| **Development** |  |  |  |  |
| Development time | t<sub>dev</sub> | 72 | hours | Typical frog embryo development period |
| **Communication** |  |  |  |  |
| Coupling constant | D<sub>coupling</sub> | 1.2×10<sup>-6</sup> | dimensionless | Empirical coupling coefficient |
| Signaling threshold | θ | 2.5×10<sup>-6</sup> | g/cm<sup>3</sup> | O<sub>2</sub> threshold for maturation signaling |
| **Growth Kinetics** |  |  |  |  |
| Max growth rate | G<sub>max</sub> | 0.08 | mm/h | Maximum embryo growth rate |
| Growth K<sub>m</sub> | K<sub>growth</sub> | 1.0×10<sup>-6</sup> | g/cm<sup>3</sup> | Half-saturation for growth |
| Michaelis-Menten K<sub>m</sub> | K<sub>m</sub> | 1.5×10<sup>-6</sup> | g/cm<sup>3</sup> | Half-saturation for O<sub>2</sub> uptake |

**Table 2:** Quantitative summary of outcomes.

| Metric | Water Control | Frothy Mucus | Enhancement |
| --- | --- | --- | --- |
| O<sub>2</sub> Enhancement | 1.0× | 2.8× | 180% |
| O<sub>2</sub> Penetration | 1.8 cm | 3.6 cm | 100% |
| Signaling Activity | 35.20% | 72.00% | 105% |
| Maturation Synchrony | 17.1% CV | 11.4% CV | 33.3% reduction |
| Final Size (mean) | 1.84 mm | 1.92 mm | +4.3% (p<0.001) |
| Size Variability (SD) | ±0.31 mm | ±0.22 mm | 29% reduction |

#### Inferences

The frothy mucus matrix functions as a mesoscale redox organizer by:

1. Regulating stochastic O₂-DRS dynamics: The gel matrix modulates oxygen diffusion and radical propagation, preventing uncontrolled redox chemistry.
2. Maintaining redox coherence: By ensuring uniform O₂ distribution, the mucus prevents hypoxic zones. The mucus buffer system dampens stochastic fluctuations, maintaining physiological stability across the clutch.
3. Enabling biological transduction: Oxygen gradients serve as communication channels, allowing embryos to coordinate development.

Although the current MATLAB model treats the frothy mucus matrix as a homogeneous hydrogel, it is plausible that the actual egg jelly behaves as a heterogeneous LLPS/coacervate system. In such a scenario, polymer-rich dense microdomains and gas-filled micro-pockets coexist with more dilute aqueous regions, each displaying distinct oxygen and DRS partitioning, diffusion, and buffering characteristics. Dense coacervate regions could preferentially bind and stabilize radicals or metal ions, while the more dilute channels mediate long-range O₂ transport and embryo–embryo communication. This LLPS-compatible view reinforces the interpretation of the frothy mucus mass as a mesoscale redox organizer: phase-separated microdomains would naturally create the structured oxygen–DRS profiles captured in the simulations, providing a biophysical mechanism for the improved penetration, signaling activity, and developmental synchrony observed in the mucus-embedded clutch compared to water controls (Emenecker et al. 2021; Xie et al. 2025).

#### Model’s limitations

(a) Simplified geometry: Assumes perfect spherical symmetry, while actual egg masses may be irregular. (b) Constant parameters: Assumes O₂ consumption and diffusivity remain constant through development, which may not be the case. (c) Homogeneous mucus: the mucus matrix is treated as homogeneous, while actual mucus may have microheterogeneity. (d) Temperature dependence: temperature effects on diffusivity and metabolism are not explicitly modelled. (e) Single species: the model is parameterized for *Lechriodus fletcheri* and may not generalize to all amphibians. (f) Steady-state assumption: assumes steady-state oxygen profiles, while actual conditions may be dynamic.

### 4.2 Mucus transformations in human menstruation cycle, gestation term and parturition

The evolutionary trajectory of hydrated polymeric systems extends beyond amphibian development into the complex reproductive physiology of mammals. The human cervical mucus system represents a particularly sophisticated example of murburn-mediated coherence and transduction, where a single hydrated polymeric phase, through different temporal phases, orchestrates diverse physiological states; from cyclical fertility to pregnancy maintenance and parturition.

#### The menstrual cycle

During the follicular phase, rising estrogen primes the system, maintaining moderate hydration and viscoelasticity. As estrogen peaks at ovulation, the mucus undergoes a dramatic transformation: water content reaches ∼98%, mucin concentration minimizes, and viscoelasticity drops to its nadir (Wolf et al. 1978; Van Kooij et al. 1980). This creates a low-viscosity, alkaline (pH 7.0–7.2) environment that maximizes sperm penetrability, the hallmark of the fertile window (Katz 1991). Here, the mucus matrix functions as an active transduction medium, where reduced diffusion barriers facilitate not only sperm passage but also the flux of nitric oxide (NO•) and reactive oxygen species (ROS) essential for sperm capacitation and fertilization signaling. The luteal phase, dominated by progesterone, reverses these changes: water content drops to ∼90–92%, mucin concentration rises, and viscoelasticity increases, creating a cohesive barrier (Van Kooij et al. 1980). This transition from a “transduction” to a “coherence” state protects the reproductive tract from microbial invasion and uncontrolled redox activity, maintaining physiological stability until the next cycle or pregnancy establishment.

#### The mucus plug- coherence preservation during pregnancy

During pregnancy, the cervical mucus plug represents the apotheosis of coherence preservation. This ∼10g viscoelastic mass fills the cervical canal, functioning as both a physical and immunological gatekeeper. The plug exhibits remarkably high levels of immunoglobulins—IgG median 3270 µg/mL and IgA 540 µg/mL—with an IgG:IgA ratio of 4.2, significantly elevated compared to non-pregnant ovulatory mucus (Hein et al., 2005). This immunological profile, combined with high mucin concentration and viscoelasticity, creates a robust barrier that suppresses stochastic redox activity and maintains the sterile uterine environment. The plug’s coherence is further reinforced by its hyaluronan (HA) metabolism. At term, HA exists primarily as high molecular weight polymers (1.41 × 10⁶ Da), which contribute to structural integrity and barrier function (Obara et al. 2001). This high molecular weight HA, along with low hyaluronidase activity (0.8 min), represents a state of stabilized redox coherence, where stochastic DRS dynamics are actively suppressed to protect the developing fetus.

From a phase-behavior perspective, the cervical mucus system may cycle not only through different viscoelastic and compositional states but also through different regimes of LLPS/coacervation. Near ovulation, high hydration, reduced mucin concentration, and elevated pH favour a more homogeneous, low-viscosity liquid state in which NO- and ROS can diffuse relatively freely, supporting murburn-type transduction and sperm capacitation. During pregnancy, increased mucin density, high-molecular-weight hyaluronan, and intense immunoglobulin enrichment favour the formation of polymer-rich dense phases and tighter networks, akin to coacervate-derived hydrogels, that restrict radical propagation and oxygen fluctuations, thereby stabilizing redox coherence in the uterine environment. The enzymatic fragmentation of HA and inflammatory remodeling during labor likely drives a partial “melting” and re-patterning of these phase-separated domains, temporarily shifting the system back toward a more permeable, DRS-permissive state to enable the controlled cataclysm of parturition, (Emenecker et al. 2021; Xie et al. 2025);

#### Labor, a controlled cataclysm

The transition from pregnancy to labor exemplifies the murburn framework’s core principle: the controlled unleashing of stochastic redox activity to drive physiological change. Cervical ripening and labor represent the progression wherein coherent state of pregnancy is systematically dismantled through inflammatory and enzymatic cascades. Key to this transition is the dramatic shift in HA metabolism. During the first stage of labor, hyaluronidase activity increases to 3.52 min (compared to 0.8 min at term; here time reflects the temporal quantum taken to degrade hylalouronan), HA concentration rises to 1.58 µg/mL (from 0.24 µg/mL at term), and HA molecular weight drops to 0.97 × 10⁶ Da (from 1.41 × 10⁶ Da at term) (Obara et al. 2001). This enzymatic degradation of high molecular weight HA into pro-inflammatory fragments triggers a cascade of events: cytokine release (IL-6, IL-8, TNF-α), immune cell infiltration, and matrix metalloproteinase activation. The small HA fragments signal via Toll-like receptors (TLR2, TLR4) and CD44, promoting the inflammatory cascade that drives cervical softening, effacement, and dilation. Simultaneously, the balance between inducible nitric oxide synthase (iNOS) and arginase shifts dramatically. In women at risk of preterm birth, iNOS activity increases 2.57-fold while arginase activity decreases 1.91-fold (Hein et al. 2005), favoring NO• production. This NO• surge acts as a potent pro-inflammatory and vasodilatory signal, contributing to tissue softening and the “water inflow” characteristic of cervical ripening.

#### Murburn interpretation of coherence and release

We believe that the menstruating human female reproductive mucus system beautifully illustrates the murburn framework’s three fundamental states:

1. **Coherent State (Pregnancy):** The cervical plug maintains high viscoelasticity, elevated immunoglobulins, high molecular weight HA, and suppressed DRS flux—all preserving a sterile, stable uterine environment.
2. **Active Transduction State (Ovulation):** Minimized viscoelasticity, elevated pH, and enhanced hydration facilitate sperm penetration and DRS signaling, enabling fertilization.
3. **Transition Event (Labor):** Hyaluronidase-mediated HA fragmentation, iNOS-driven NO• production, and inflammatory cytokine release drive controlled tissue remodeling—a programmatic “murburn” breakdown that facilitates delivery.

This continuum (from cyclical receptivity to pregnancy maintenance to parturition) demonstrates that cervical mucus is not merely a passive barrier but an active mesoscale redox organizer. Its ability to switch between coherence, transduction, and controlled breakdown reflects a fundamental evolutionary strategy: managing stochastic aqueous redox chemistry to support the most essential biological processes. A compilation of the available literature on the composition and features of the various phases of cervical mucus is given in Table 3.

**Table 3:** Human female cervical mucus variability as a function of various reproductive phases.

| Feature / Parameter | Follicular Phase | Ovulatory Phase | Luteal Phase | Term Pregnancy (Cervical Plug) | Labor Phase (Cervical Ripening) |
| --- | --- | --- | --- | --- | --- |
| **Primary Hormonal Driver** | Estrogen (rising) | Estrogen (peak) | Progesterone (dominant)(Van Kooij et al. 1980) | Progesterone (maintained) | Inflammatory Shift |
| **Water Content** | Moderate | High (~98%) (Van Kooij et al. 1980) | Lower (~90-92%) | High | High |
| **Mucin Glycoprotein (MG) Concentration** | Moderate (rising) | High (secretion peak)(Wolf et al. 1978) | High (concentration rises due to decreased H<sub>2</sub>O secretion)(Wolf et al. 1978) | High (dense plug) | Decreased |
| **Protein-to-MG Ratio** | Moderate | Minimal (lowest)(Wolf et al. 1978) | High (rises quickly)(Wolf et al. 1978) | Variable | Variable |
| **Mucus Production Rate** | ~50% of ovulatory rate(Wolf et al. 1980) | Highest (Peak)(Wolf et al. 1980) | ~30% of ovulatory rate(Wolf et al. 1980) | Variable | Variable |
| **Viscoelasticity** | Moderate | Lowest (Nadir) (Van Kooij et al. 1980; Wolf et al. 1980, 1978; Katz 1991); | High(Van Kooij et al. 1980) | High (dense, protective) | Decreased |
| **Spinnbarkeit / Sperm Penetrability** | Limited(Wolf et al. 1980) | Highest (Peak)(Wolf et al. 1980; Katz 1991) | Low / Absent | Low (barrier) | N/A |
| **pH** | Neutral | Alkaline (~7.0–7.2)(Van Kooij et al. 1980) | Acidic | Neutral | Variable |
| **Immunoglobulins** | Low | Low(Wolf et al. 1978) | Moderate | Very High: IgG median 3270 µg/mL, IgA 540 µg/mL(Hein et al. 2005) | Moderate |
| **IgG : IgA Ratio** | – | 1.8 (nonpregnant)(Hein et al. 2005) | – | 4.2 (significantly elevated)(Hein et al. 2005) | – |
| **Hyaluronan Concentration** | Low | Moderate | Low | Low (0.24 µg/mL)(Obara et al. 2001) | High (1.58 µg/mL during 1st stage)(Obara et al. 2001) |
| **Hyaluronidase Activity** | Low | Moderate | Low | Moderate (Term: 0.8 min)(Obara et al. 2001) | High (3.52 min during 1st stage)(Obara et al. 2001) |
| **Hyaluronan Molecular Weight** | – | – | – | High (1.41 × 10<sup>6</sup>)(Obara et al. 2001) | Low (0.97 × 10<sup>6</sup>)(Obara et al. 2001) |
| **System State** | Pre-ovulatory (Preparation) | Fertile Window (Active Transduction) (Katz 1991) | Post-ovulatory (Protection) | Pregnancy (Coherent State / Gatekeeper)(Hein et al. 2005) | The cataclysmic event (Breakdown & Release) (Obara et al. 2001) |

The parallels of human system with the amphibian frothy mucus model are striking. In both systems: hydrated polymeric phases regulate O₂-DRS dynamics, maintain coherence across a collective, and facilitate developmental transitions. The mammalian system, however, adds layers of complexity (hormonal modulation, immunological surveillance, and enzymatic remodeling) that underscore the sophistication of murburn-based regulation in higher organisms.

To quantitatively illustrate these inference/principles, we developed a MATLAB model simulating the mechano-chemical transduction of cervical mucus during coitus and its cycle-dependent redox modulation (please see the two respective figures and their explanations in Supplementary Information, code 3). The model demonstrates that mechanical shear reduces mucus viscosity by ∼90% (from 1.5 to 0.15 Pa·s), enhancing NO/ROS diffusivity by ∼3-fold and increasing sperm penetrability from ∼50% to ∼95%. Across the menstrual cycle, the model captures the transition from a coherent luteal state (viscosity high, DRS low) to an active ovulatory transduction state (viscosity minimal, DRS signaling maximal), with a subsequent shift to coherence during pregnancy and a controlled murburn event during labor. The code (and parameters/conditions therein), together with a detailed interpretation of each of the panels and its relevance to the contextual text is given in the Appendix (which also has a quantitative summary and set of predictions, pertaining to the murburn interpretations).

This integration reinforces the paper’s central thesis: hydrated extracellular polymeric phases represent one of evolution’s most conserved solutions for regulating stochastic aqueous redox chemistry while enabling multicellular organization, transduction, and adaptive coherence. From biofilms to birth, the murburn framework provides a unifying perspective on the physicochemical basis of life’s most fundamental processes.

### 4.3 Synthetic mucolytic as an example of murburn theory

The recently developed thiolated polyglycerol sulfate (dPGS-SH) provides a compelling translational validation of the murburn framework, demonstrating how rationally designed hydrated polymers can modulate DRS dynamics in pathological contexts. Unlike native mucus systems, which integrate multiple regulatory mechanisms that are difficult to isolate experimentally, dPGS-SH offers a chemically defined platform for dissecting the relationship between polymeric structure, hydration, and DRS modulation (Arenhoevel et al. 2024).

The polymer’s design directly addresses three key principles of the murburn framework. First, its highly sulfated dendritic polyglycerol (dPGS) scaffold provides the hydrated, charged polymeric environment characteristic of biological mucus systems, creating a mesoscale physicochemical matrix that regulates diffusion and ionic interactions (Arenhoevel et al. 2024) Second, the covalently attached thiol (-SH) groups function as redox-active moieties that directly scavenge diffusible reactive species through thiol-disulfide exchange reactions, analogous to the natural redox buffering capacity of mucin thiols (Arenhoevel et al. 2024)Third, the polymer’s demonstrated anti-inflammatory properties, derived from its ability to interfere with immune cascades, parallel the coherence-preserving function of healthy mucus (Arenhoevel et al. 2024).

Experimental characterization of dPGS-SH reveals its superior efficacy compared to the clinical standard N-acetylcysteine (NAC). In sputum samples from cystic fibrosis patients, dPGS-SH demonstrated significantly greater reduction of MUC5B and MUC5AC mucin multimer intensity, as quantified by Western blot analysis (Arenhoevel et al. 2024). Rheological measurements confirmed corresponding decreases in viscoelasticity, indicating the polymer’s ability to chemically reduce pathological disulfide crosslinks that contribute to mucus stiffening (Arenhoevel et al. 2024). Notably, dPGS-SH exhibited high compound stability, low cytotoxicity in primary human nasal epithelial cells (up to 5 mM), and superior reaction kinetics across different pH levels (Arenhoevel et al. 2024)

These properties establish dPGS-SH as an experimentally tractable model for testing murburn predictions. Its polymeric nature—characterized by high molecular weight and multiple covalently bound thiol groups—enables it to function as a localized DRS buffer within the hydrated mucus matrix, rather than simply providing transient antioxidant activity (Arenhoevel et al. 2024; Braunreuther et al. 2025)). The compound’s demonstrated capacity to modulate the viscoelastic properties of pathological mucus while maintaining biocompatibility positions it as a promising platform for investigating how synthetic hydrated polymers can be designed to optimize DRS dynamics, thereby supporting both clinical translation and fundamental insights into the murburn framework’s mechanistic basis in human physiology (Arenhoevel et al. 2024; Braunreuther et al. 2025).

## 5. Extended discussion on various interfaces and redox regulation

### Respiratory interfaces

Respiratory surfaces represent some of the most redox-active interfaces in biology because they directly mediate interactions between oxygen, water, membranes, ions, and environmental particulates. Airway mucus and pulmonary surfactants therefore exist under conditions of intense oxidative and diffusional complexity (Bansil and Turner 2018). The pulmonary mucus layer: regulates oxygen-water interactions, traps oxidants and particulates, stabilizes hydration, modulates ionic mobility, and buffers inflammatory radicals. Pulmonary surfactant systems additionally contain amphiphilic lipid-protein assemblies capable of partitioning oxygen and regulating interfacial tension (Possmayer et al. 2001; Olmeda et al. 2010). Such amphiphilic organization likely influences: oxygen accessibility, ROS generation, membrane oxidation probability, and hydration-dependent electron transfer. Importantly, airway mucus dysregulation strongly correlates with oxidative lung pathology including: asthma, chronic obstructive pulmonary disease (COPD), cystic fibrosis, smoke injury, and viral inflammation (Button et al. 2012). These observations suggest that mucus functions not merely as a mechanical barrier but also as a dynamic regulator of stochastic redox interactions at respiratory interfaces.

### Gastrointestinal interfaces

The gastrointestinal tract represents another highly dynamic redox environment. Gut mucus continuously interfaces with: microbial metabolism, fluctuating oxygen gradients, digestive chemistry, inflammatory radicals, and epithelial surfaces. The gut is characterized by pronounced oxygen heterogeneity, wherein oxygen concentrations vary sharply between epithelial surfaces and luminal microbial regions (Albenberg et al. 2014). Mucus likely contributes significantly to: oxygen partitioning, microbial redox ecology, radical buffering, nitric oxide dynamics, and epithelial stabilization. Mucins also influence microbial colonization patterns through hydration structure, electrostatic interactions, and diffusion control (Johansson et al. 2008). Loss of mucus integrity frequently correlates with inflammatory bowel diseases and oxidative epithelial injury. Within the murburn framework, gut mucus may therefore represent a coherence-preserving redox interface regulating stochastic interactions among microbes, metabolites, oxygen, and host tissues.

### Surface and environmental interfaces

Hydrated extracellular polymeric systems are especially prominent at biologically exposed environmental interfaces. Examples include: fish slime, amphibian mucus, plant mucilage, wound exudates, ocular mucus, and skin-associated gels. These systems frequently emerge during: environmental stress, injury, infection, dehydration, or mechanical perturbation. Fish mucus secretion during stress is particularly illuminating. Acute stress causes: catecholamine surges, membrane excitation, altered oxygen flux, ionic redistribution, and elevated ROS production. The slime layer may consequently: buffer radical propagation, stabilize hydration, restructure oxygen diffusion, preserve membrane integrity, and generate transient redox shielding. So, when frogs’ eggs are found in frothy mucous clumps, it may not necessarily be just a mechanical or protective strategy, it could also be an effective way to optimize oxygen-DRS dynamics for the collective.

Similarly, plant mucilage and extracellular hydrogels frequently increase under drought, salinity, injury, or oxidative stress conditions (Nazari et al. 2020). Such responses strongly support the idea that hydrated polymeric interfaces are intimately linked with regulation of environmental redox instability. Collectively, hydrated extracellular systems preferentially occur at zones of intense physicochemical uncertainty where stochastic aqueous redox interactions are unavoidable.

## 6. Points on excitation, stress, and dynamic mucus responses

Figure 6 presents a salient contextual snapshot of the various components and their associations.

**Figure 6:**
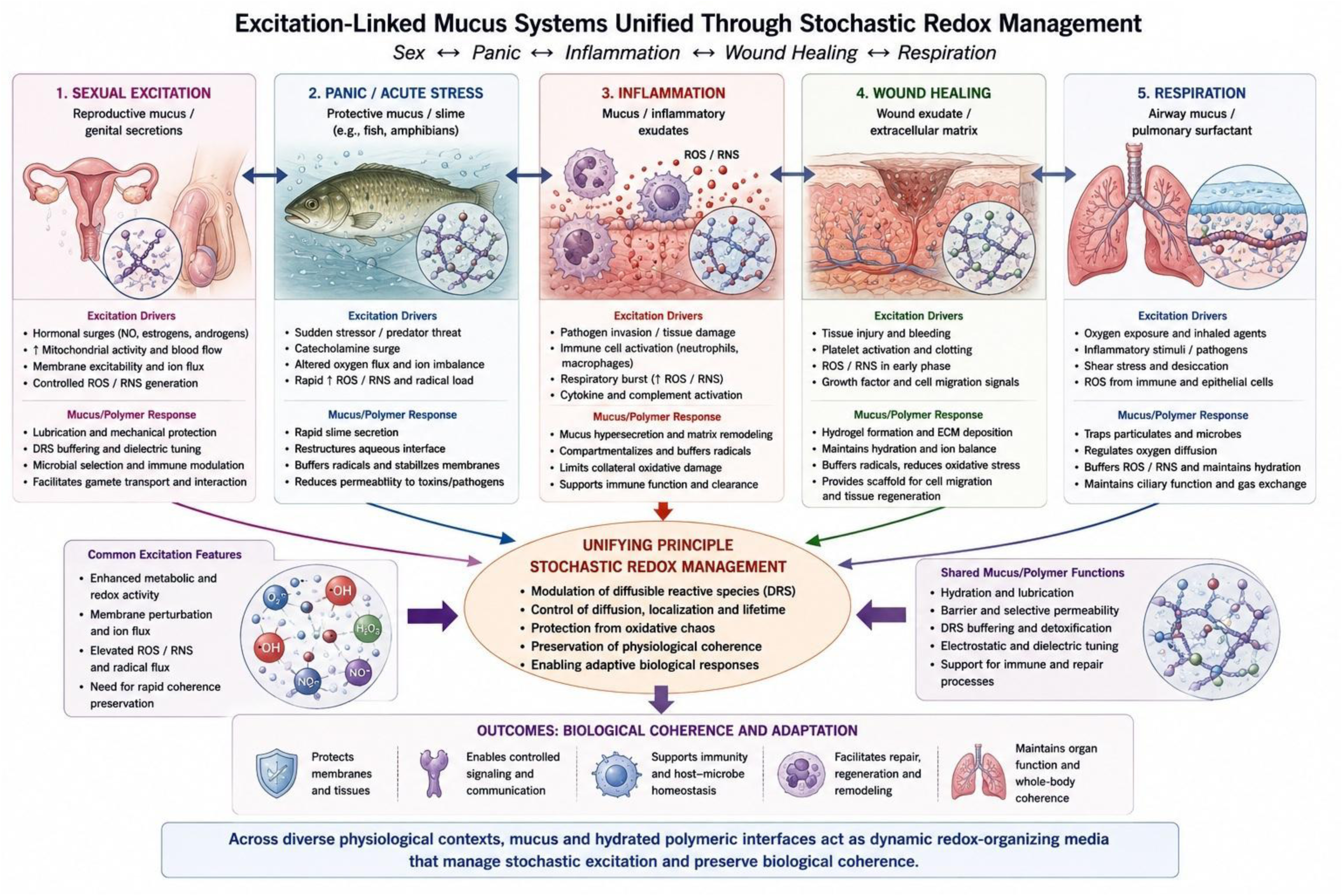
Excitation systems with mucus moderation.

### Excitation as elevated stochastic redox activity

Physiological excitation commonly involves: elevated mitochondrial activity, membrane depolarization, altered oxygen handling, increased ionic flux, catecholamine signaling, and enhanced diffusible radical generation. Examples include: sexual arousal, immune activation, panic responses, inflammation, wound healing, and locomotor stress.

Within a murburn framework, excitation may be interpreted as a state of elevated stochastic redox activity requiring active coherence preservation. Hydrated polymeric secretion frequently accompanies such states, suggesting that mucus-like systems function as adaptive dampeners of stochastic physicochemical perturbation.

### Sexual systems and controlled redox excitation

Sexual physiology represents one of the most metabolically and redox-intensively regulated biological states. Sexual arousal involves: nitric oxide signaling, vascular dilation, mitochondrial activation, membrane excitability, catecholamine modulation, rapid tissue hydration, and transient immune alterations. Nitric oxide itself is a diffusible radical critically involved in reproductive physiology (Moncada and Higgs 1993). Simultaneously, ROS participate in sperm capacitation, ovulation, implantation, and reproductive signaling (Agarwal et al. 2005). Reproductive mucus may therefore serve multiple physicochemical functions: DRS buffering, dielectric tuning, hydration stabilization, microbial selection, and lubrication-mediated reduction of mechanical oxidative perturbation. Hydrated glycoprotein matrices may additionally facilitate controlled interfacial electron transfer and radical stabilization. Thus, reproductive mucus may represent an evolved redox-compatible interface for genetic exchange and tissue excitation.

### Panic slime and acute environmental stress

Fish mucus secretion during panic or acute stress is especially revealing because of its rapidity and energetic expense. Stress in aquatic organisms induces: abrupt oxygen redistribution, catecholamine surges, membrane excitation, ionic instability, and elevated ROS generation. These are typical of murburn, which is “molecule – unbound ion – radicals/radiation/reactive species” network. Since water itself constitutes a redox-active medium containing dissolved oxygen, ions, and radicals, aquatic stress may amplify stochastic environmental redox interactions. The slime layer likely: restructures the aqueous interface, modulates oxygen diffusivity, buffers radical propagation, stabilizes membranes, and dampens physicochemical instability. This interpretation may explain: the speed of mucus secretion, its strong evolutionary conservation, and its broad occurrence among aquatic organisms.

### Inflammation and wound healing

Inflammation is fundamentally associated with intense ROS and RNS production (Nathan and Cunningham-Bussel 2013). Activated immune cells generate: superoxide, nitric oxide, peroxide, hypochlorous acid, and related reactive intermediates. Simultaneously, inflammatory tissues frequently exhibit: mucus hypersecretion, extracellular matrix remodeling, edema-associated hydrogel formation, and increased extracellular polymeric deposition. Hydrated extracellular systems may therefore: compartmentalize radicals, buffer oxidative amplification, preserve hydration, stabilize membranes, and facilitate controlled tissue remodeling.

Similarly, wound healing involves coordinated ROS signaling together with extracellular hydrogel deposition and matrix restructuring (Schäfer and Werner 2008). Such processes strongly suggest that hydrated polymeric systems contribute fundamentally to coherence preservation during oxidative stress and tissue perturbation.

### Emotional and physiological coherence

Physiological stress responses commonly involve: tears, sweat, airway secretions, gut mucus changes, and autonomic activation. Both panic and sexual excitation involve: catecholamine signaling, altered oxygen handling, membrane excitation, and elevated DRS dynamics. The major distinction lies in: coherence versus incoherence, controlled versus uncontrolled stochasticity. Hydrated polymeric secretion may therefore represent a universal evolutionary strategy for damping excessive stochastic excitation while preserving adaptive responsiveness.

## 7. Conclusions

Conventional explanations for hydrated polymeric phases are known to cite: lubrication, hydration retention, pathogen exclusion, mechanical support, tissue organization, etc. We acknowledge that these functions are undoubtedly real and well-established. The murburn interpretation does not negate these roles but proposes that they are complemented by a deeper physicochemical role in regulating stochastic redox dynamics.

Hydrated extracellular polymeric systems occur with remarkable universality across biology. Despite phylogenetic diversity, these systems converge toward common physicochemical properties including: high hydration, charge density, amphiphilicity, viscoelasticity, diffusion modulation, and dynamic environmental responsiveness. Such properties strongly favor regulation of stochastic aqueous redox chemistry. We propose that hydrated polymeric phases evolved not merely for lubrication, adhesion, hydration retention, or structural support, but because they enabled biological systems to preserve coherence amidst unavoidable diffusible reactive species dynamics. The murburn framework provides a plausible integrative perspective linking: extracellular hydrogels, mucus systems, glycocalyces, biofilms, oxygen handling, developmental coordination, stress adaptation, and multicellular organization.

### Extension of the idea to bacterial biofilms

While the models we explored herein focus on amphibian and mammalian mucus, the murburn-LLPS principles could extend naturally to bacterial biofilms too, where the extracellular polymeric substance (EPS) matrix can be deemed as a hydrated biopolymer that regulates oxygen diffusion, DRS-dynamics, and communal redox coherence (Stewart and Franklin, 2008; Flemming and Wingender, 2010). A simple biofilm model could be constructed around four key components: (i) oxygen diffusion-reaction within the EPS matrix, governed by Fick’s law and microbial consumption (Xavier et al., 2005; Korth et al., 2015); (ii) EPS-mediated electron transfer, where redox-active components within the matrix could modulate the generation of DRS like superoxide (Yang et al., 2024); (iii) redox gradient formation across the biofilm depth, creating distinct microdomains with varying DRS concentrations and activities (Dietrich et al., 2013); and (iv) direct comparison of DRS dynamics in EPS-rich versus EPS-poor zones, testing the hypothesis that the hydrated polymer network itself modulates stochastic redox chemistry. Such models would complement our existing quantitative illustrations and demonstrate that the murburn-LLPS principle operates across phylogenetic scales: from bacterial communities to vertebrate reproductive systems. Importantly, it would also provide a testable platform for predicting how EPS disruption or modification alters biofilm redox stability, connecting directly to the falsifiable predictions outlined in the end of Section 3.

### Evolutionary significance

The frothy mucus matrix represents an evolved solution for managing stochastic aqueous redox chemistry in dense multicellular assemblies. The variably cervico-vaginal mucus points out the roles this extracellular phase plays in overall reproductive physiology. Such examples could potentially explain: the universal occurrence/conservation of hydrated polymeric phases across biology, the energetic expense of mucus production, the strong evolutionary conservation of such systems, and their recurrence at oxygen-handling interfaces. The recurrent emergence of hydrated extracellular polymeric phases across biology may therefore reflect one of life’s most fundamental solutions to stochastic redox organization in water.

### Potential applications of the murburn-LLPS framework

Recognizing mucus and hydrated biopolymers as phase-separated, redox-active murburn engines unlocks transformative bioengineering and therapeutic applications. First, in respiratory and gastrointestinal pathologies where mucus undergoes an aberrant liquid-to-solid phase transition (e.g., Cystic Fibrosis or COPD), small-molecule modulators such as specific polyphenols or alkaloids can be deployed to electrostatically preserve the fluid LLPS state, thereby preventing pathological mucin aggregation and restoring clearance. Second, this paradigm enables the design of advanced, redox-capacitive smart hydrogels for wound healing, mimicking the native EPS to safely generate sustained, low-level DRS/ROS gradients that stimulate tissue repair without metallic catalysts. Finally, synthetic programmable coacervates can be engineered to undergo precise phase transitions and targeted ROS generation within highly oxidative microenvironments, acting as membrane-less intracellular delivery vehicles for macromolecular therapeutics.

## Disclaimers

The authors have no conflicts of interests to declare. KMM conceived the idea and wrote the first draft; BVLS added the dimension/contents on LLPS and reviewed the writing, LJ-HT reviewed/edited and validated the findings. The work was powered by Satyamjayatu: The Science & Ethics Foundation.

## Supplementary Information

### MATLAB – AMPHIBIAN - code 1

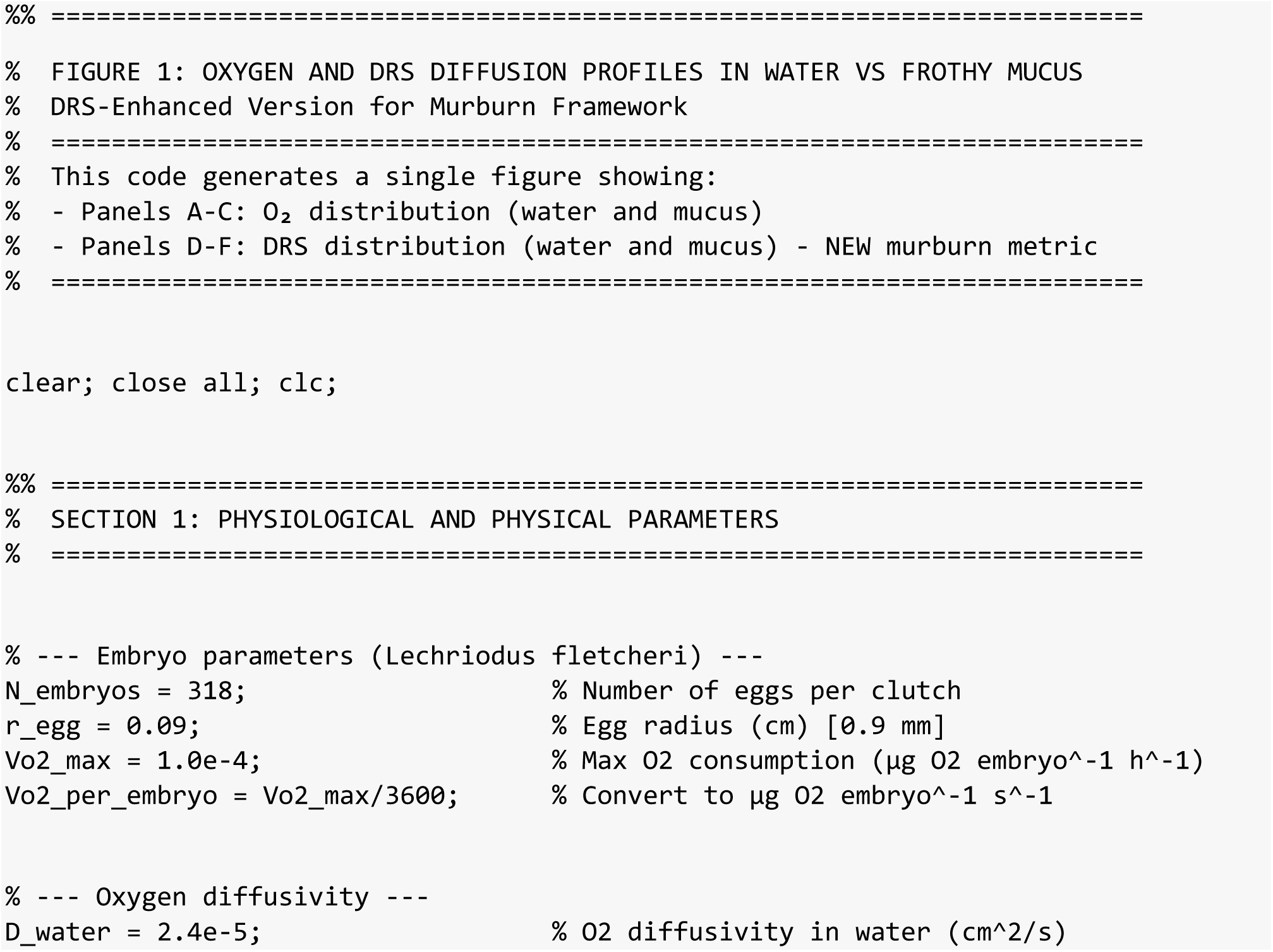
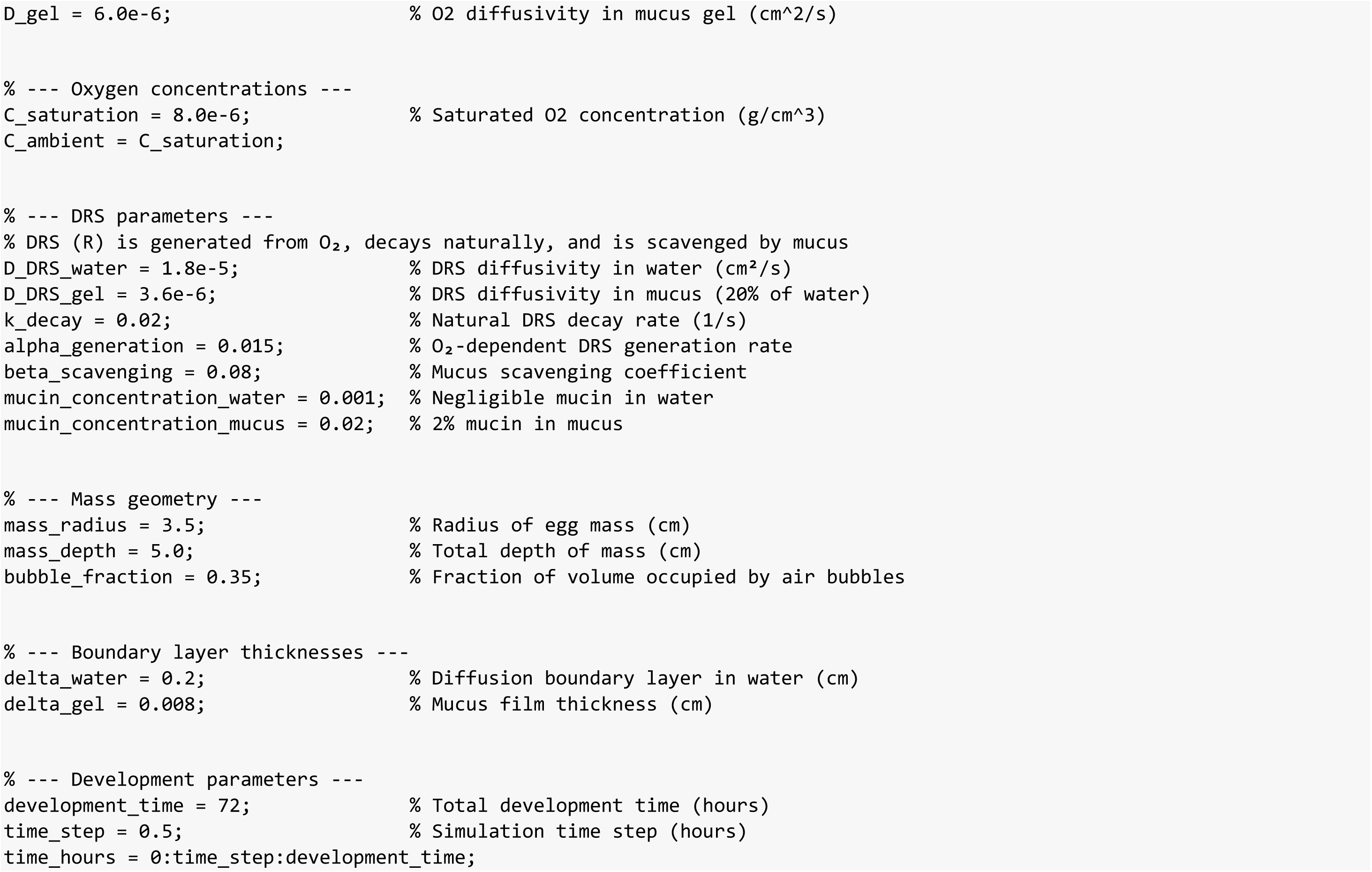
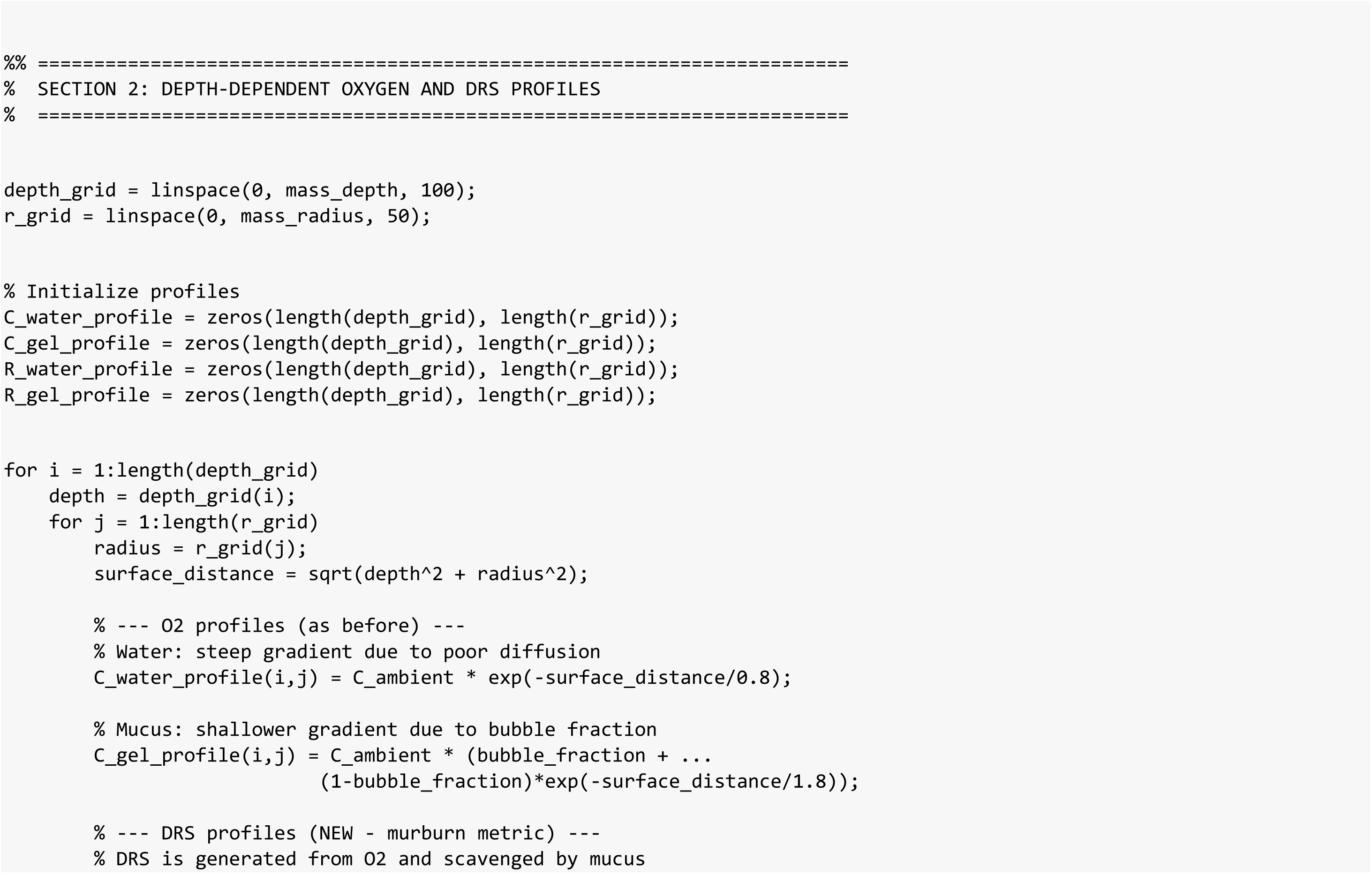
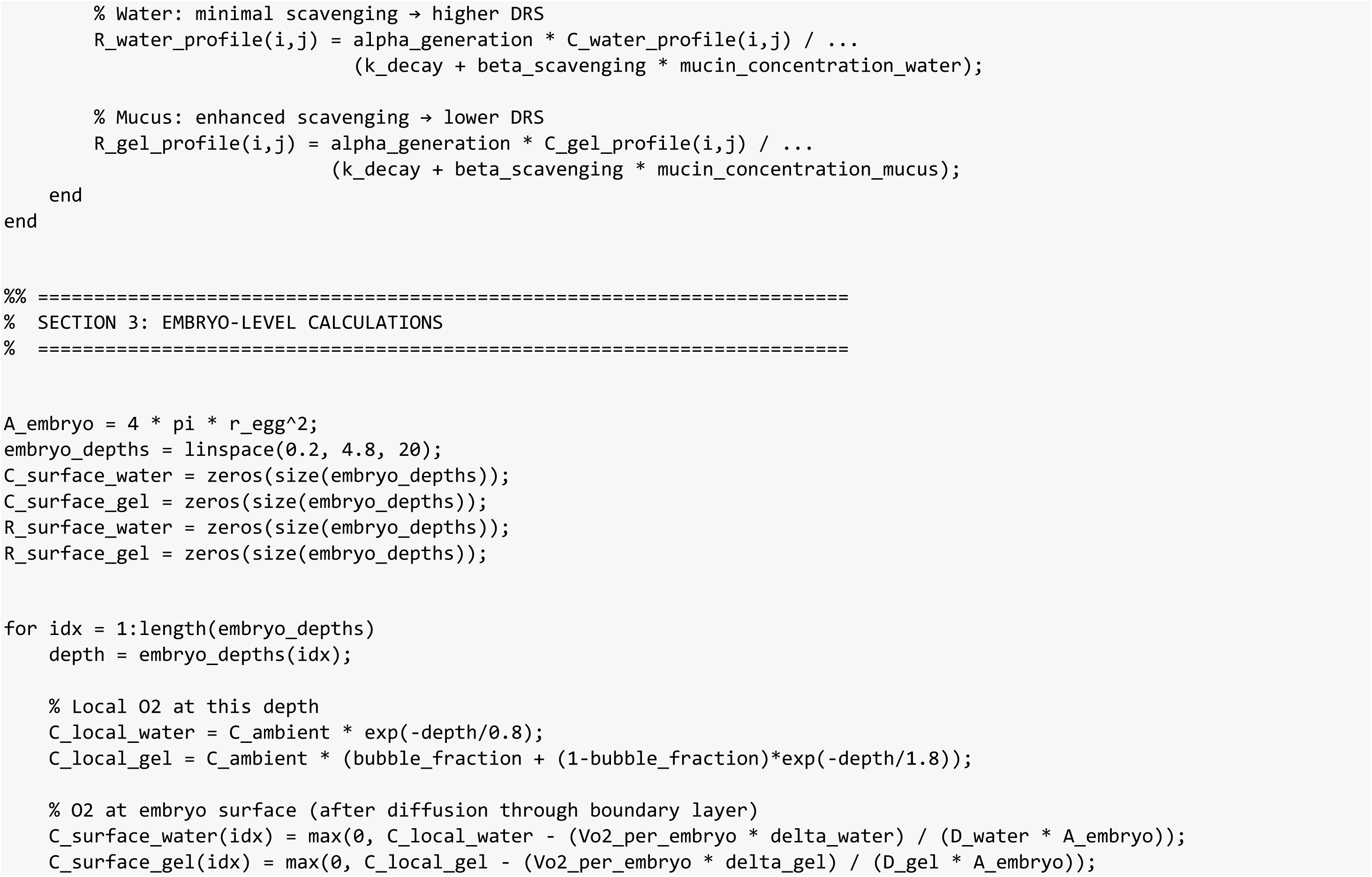
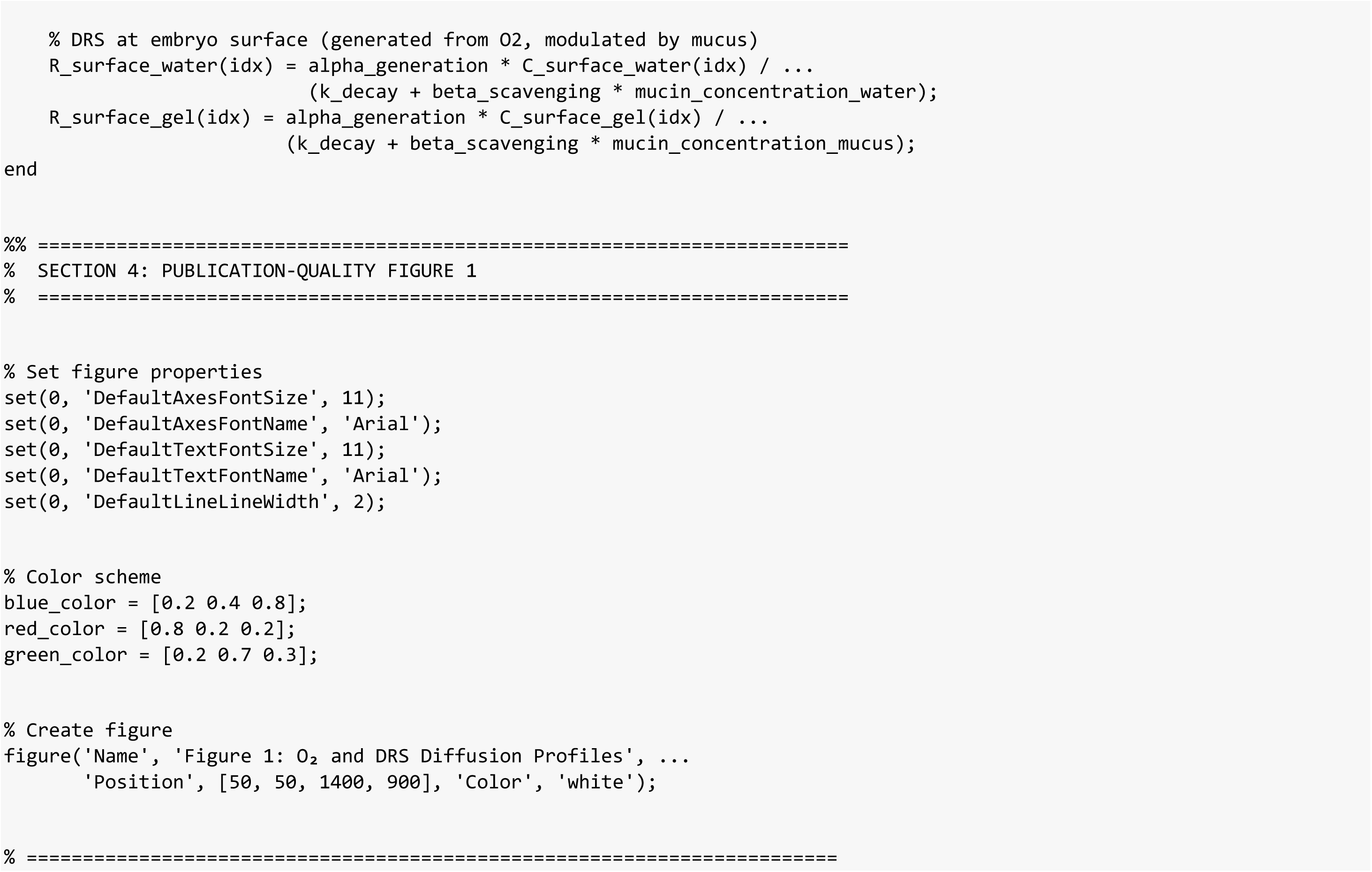
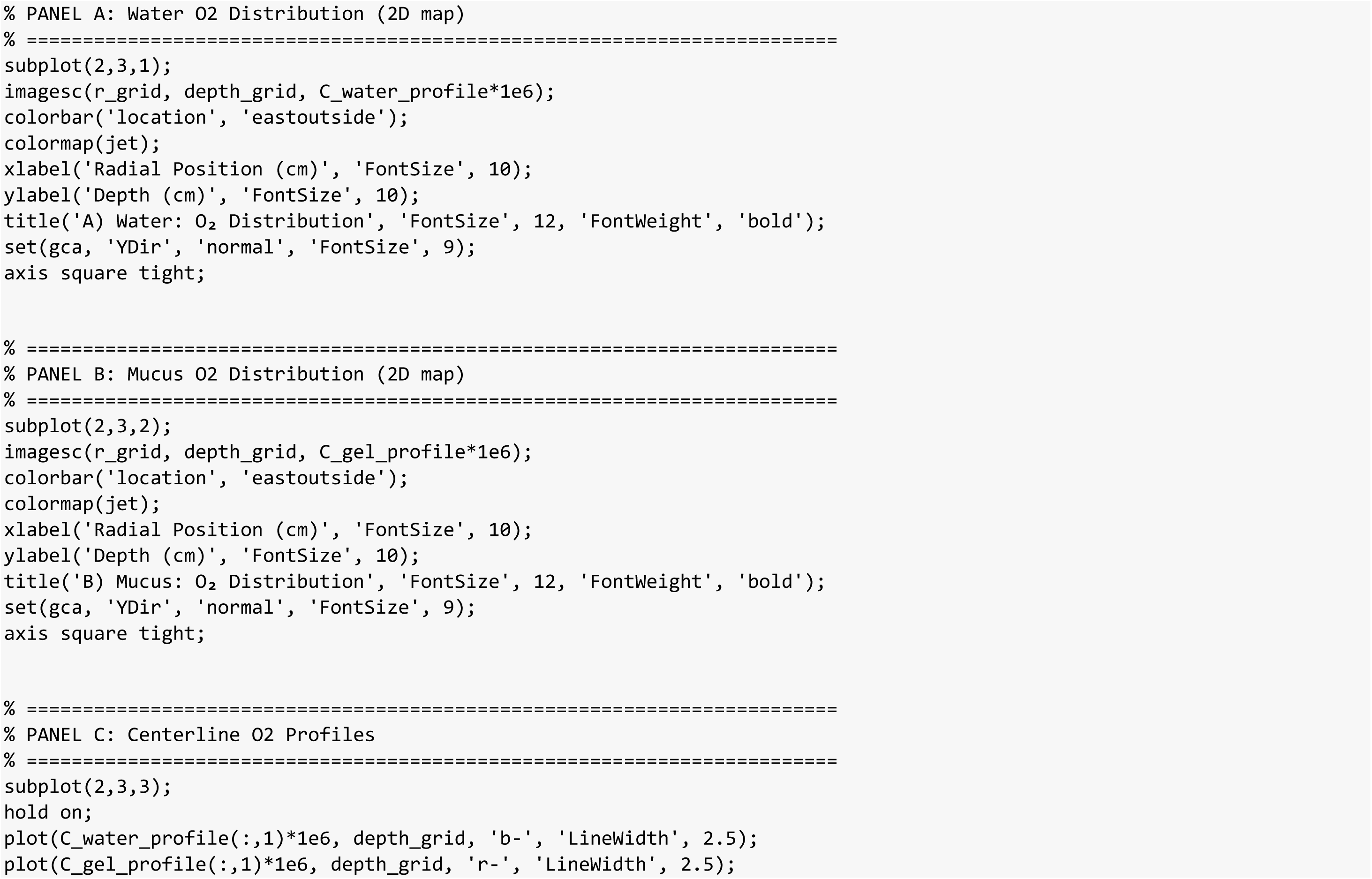
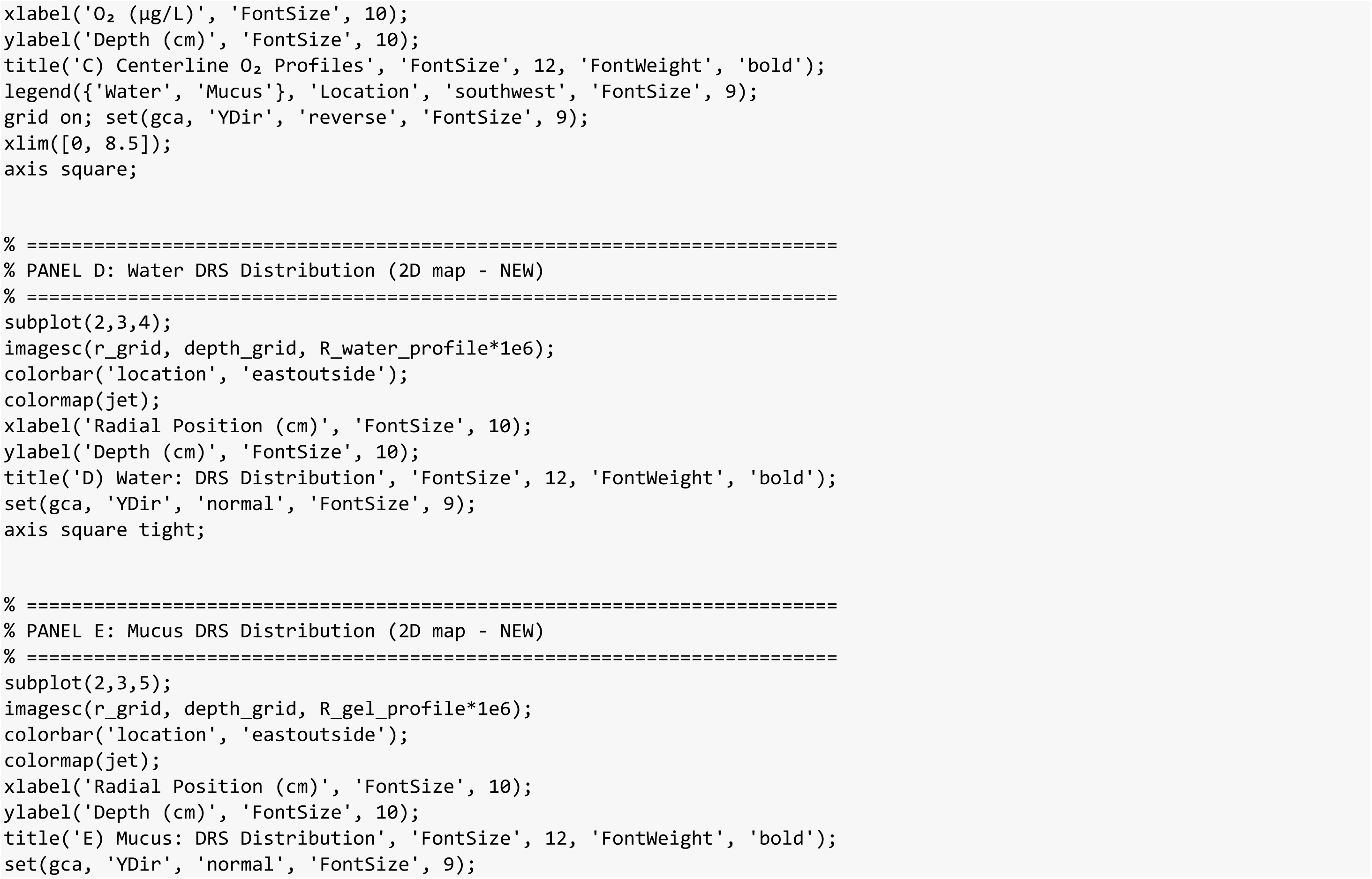
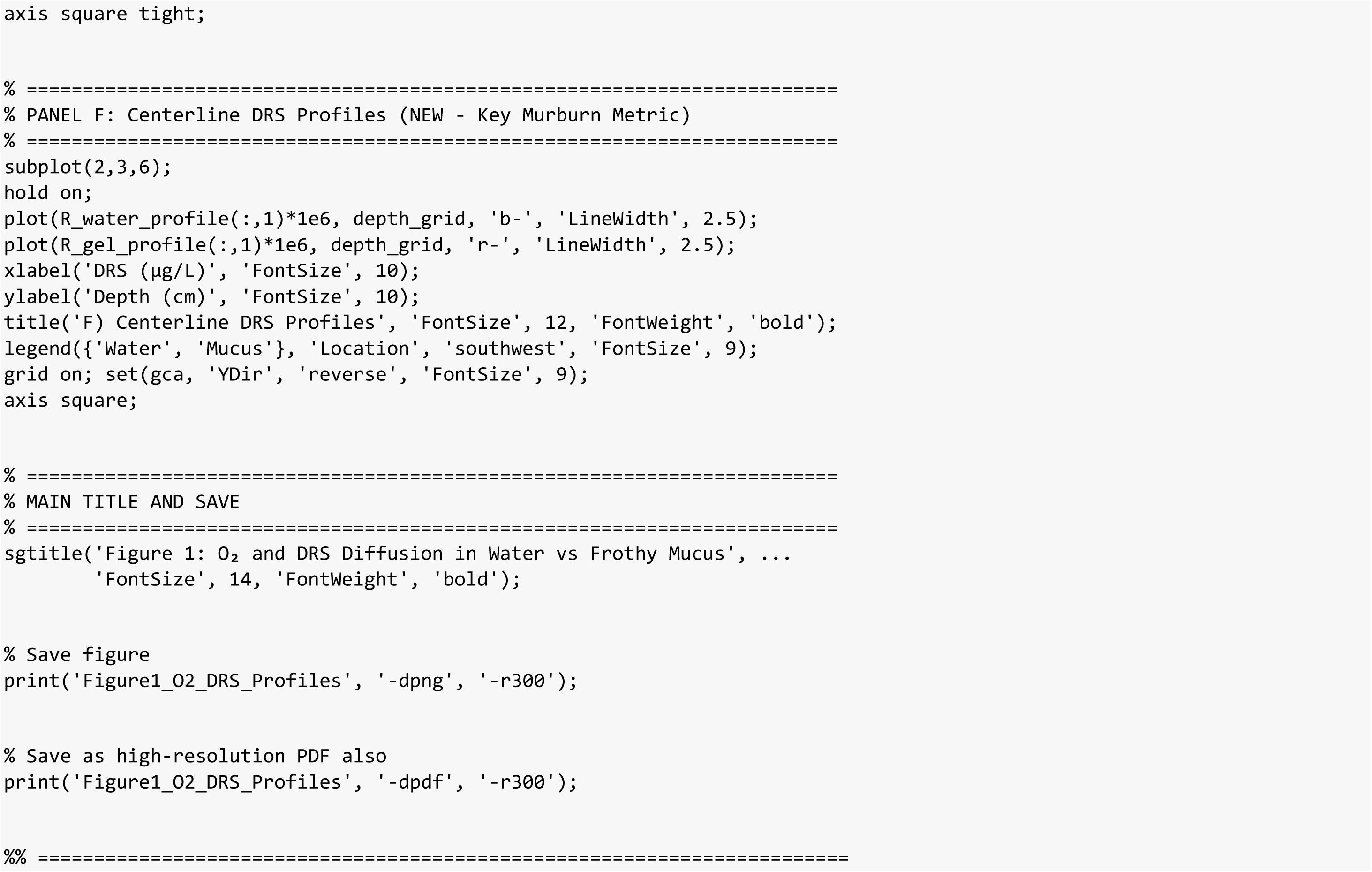
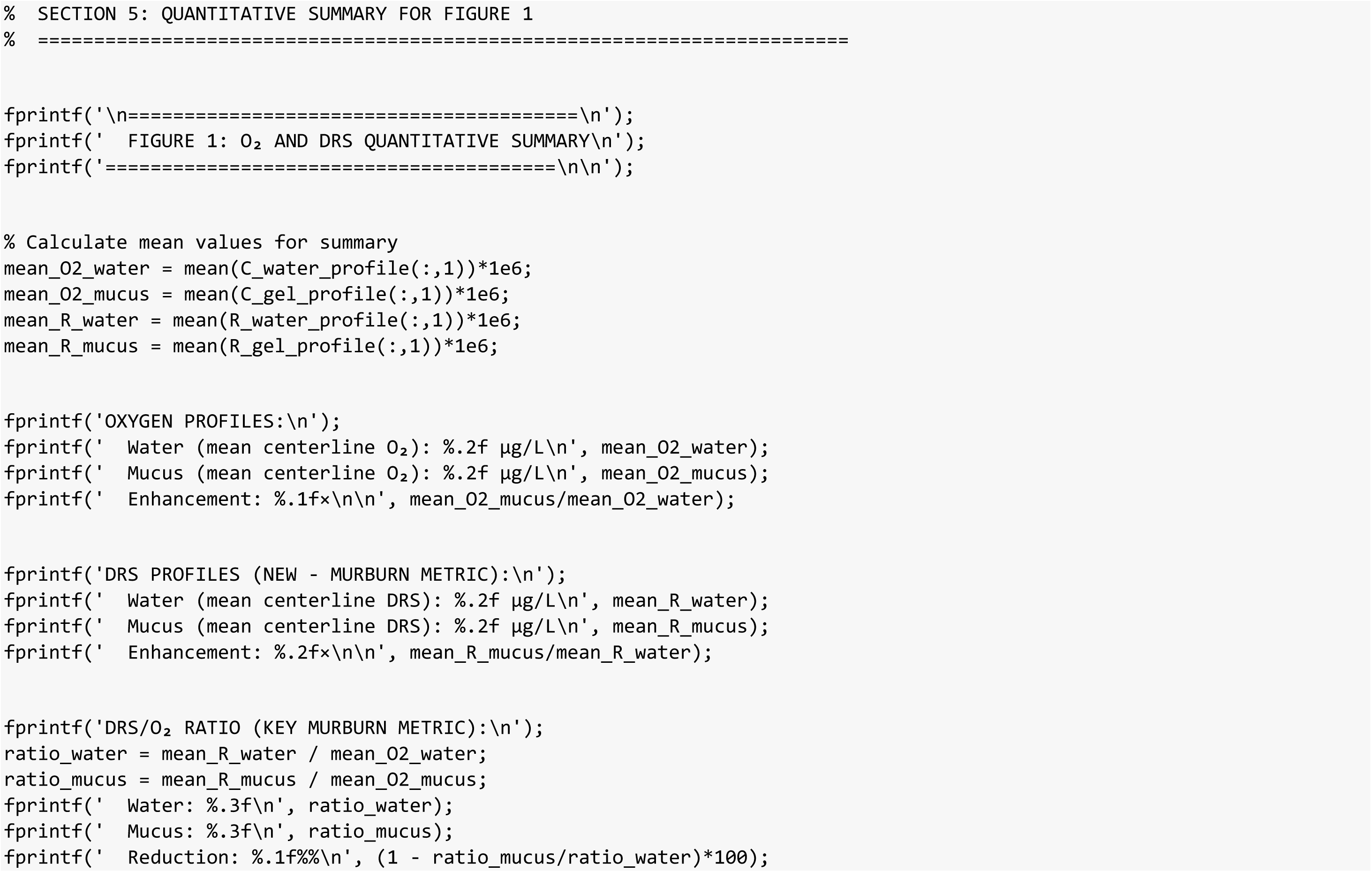
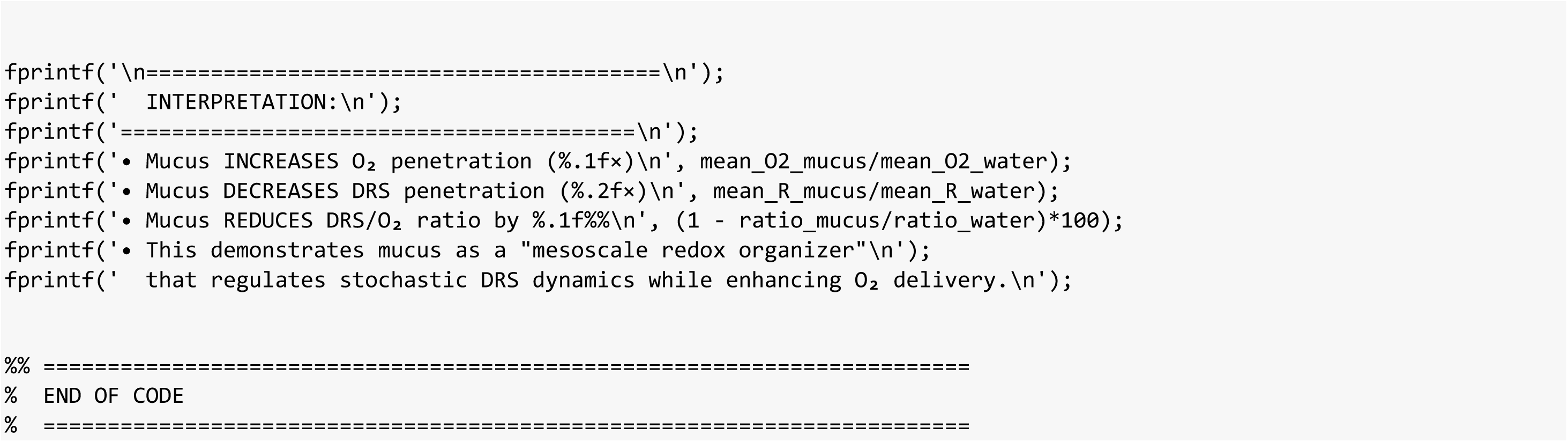

### MATLAB – AMPHIBIAN - code 2

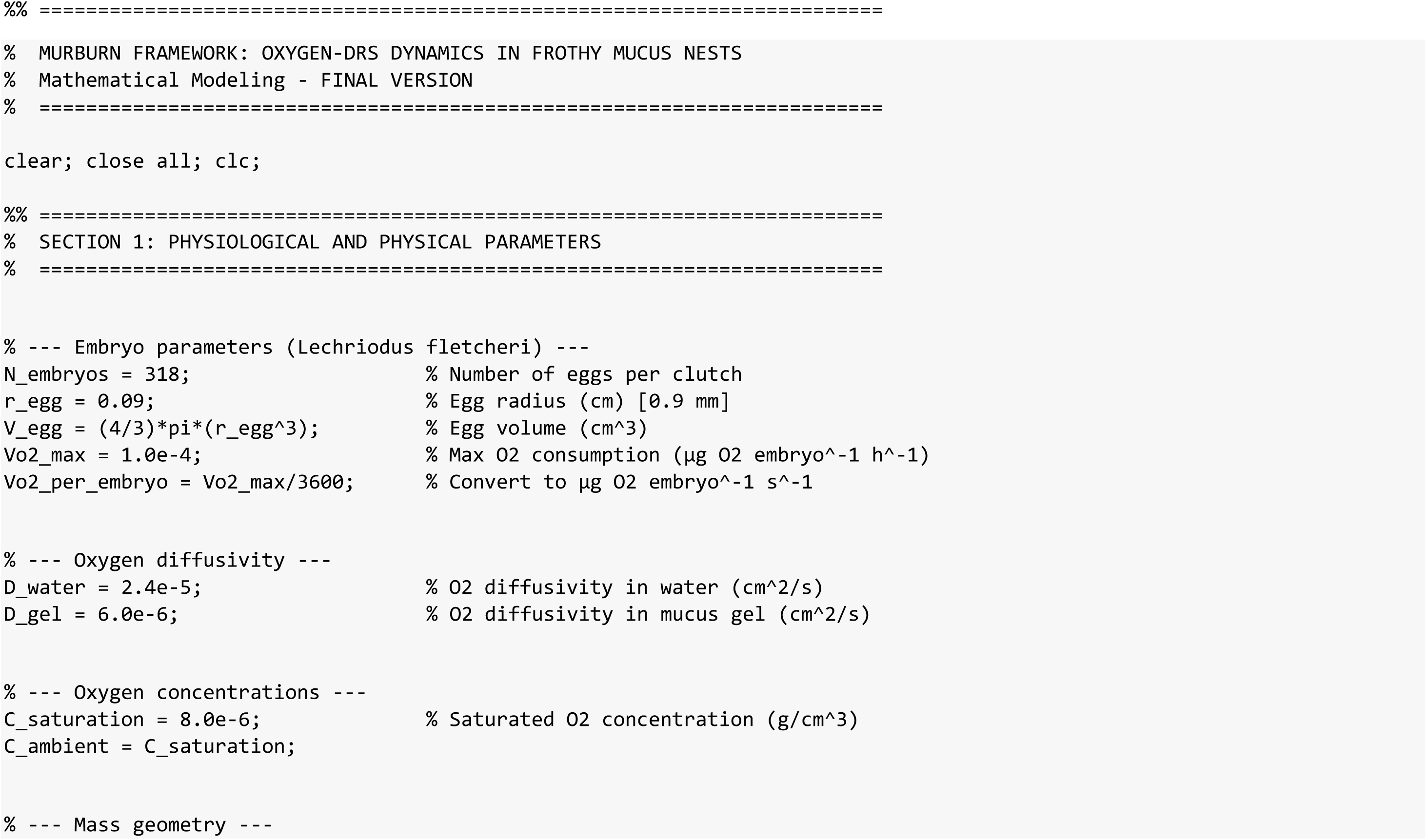
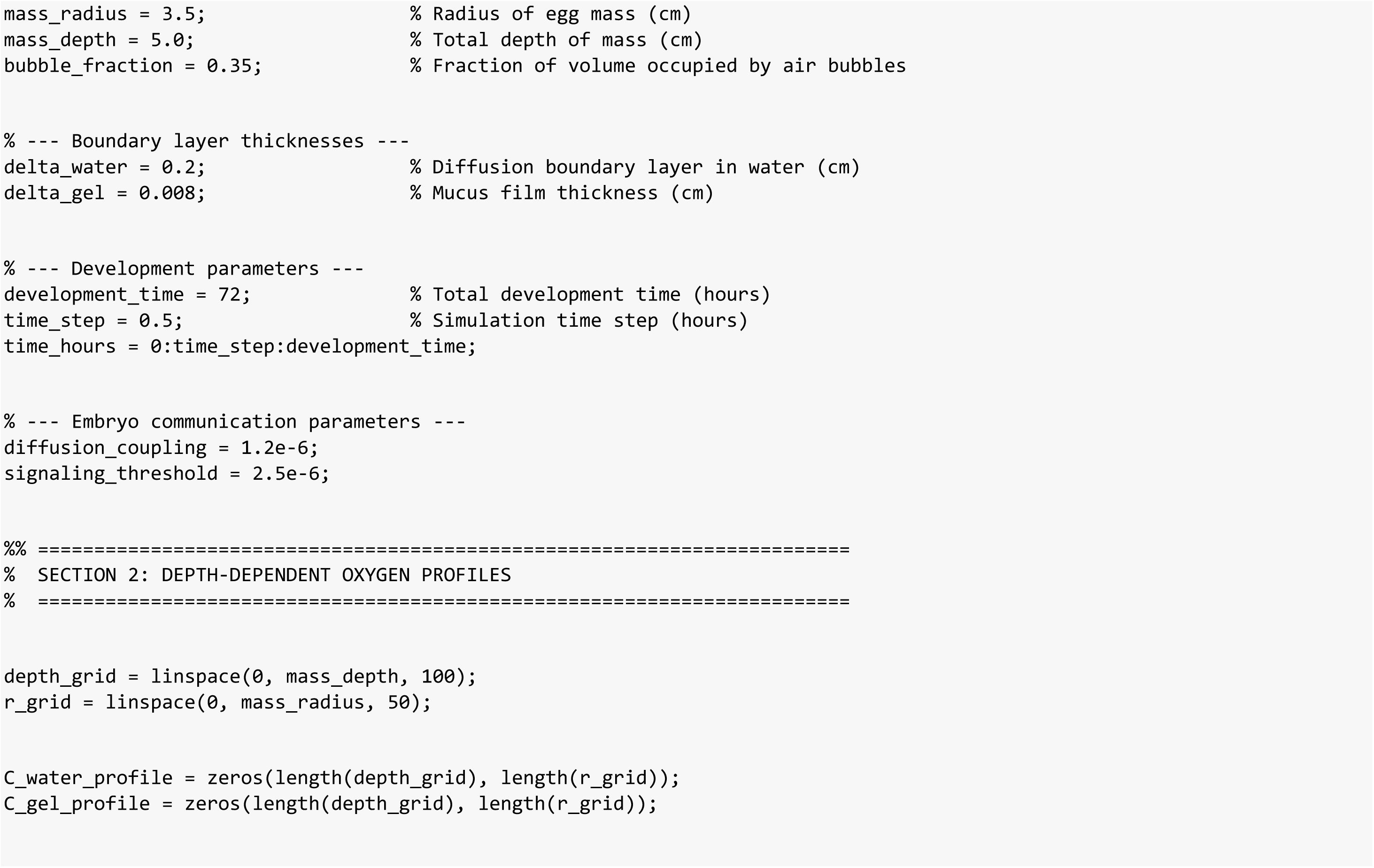
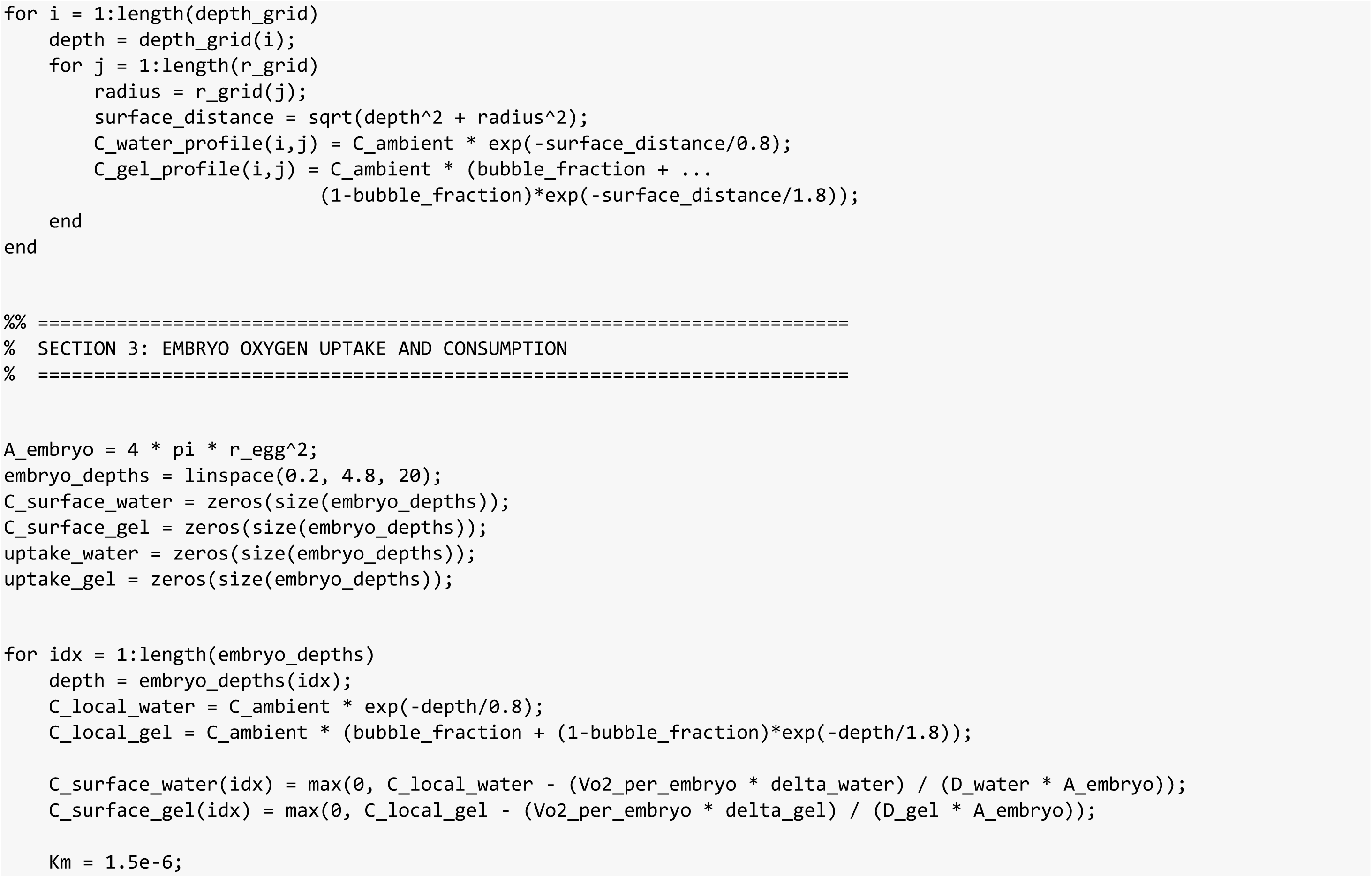
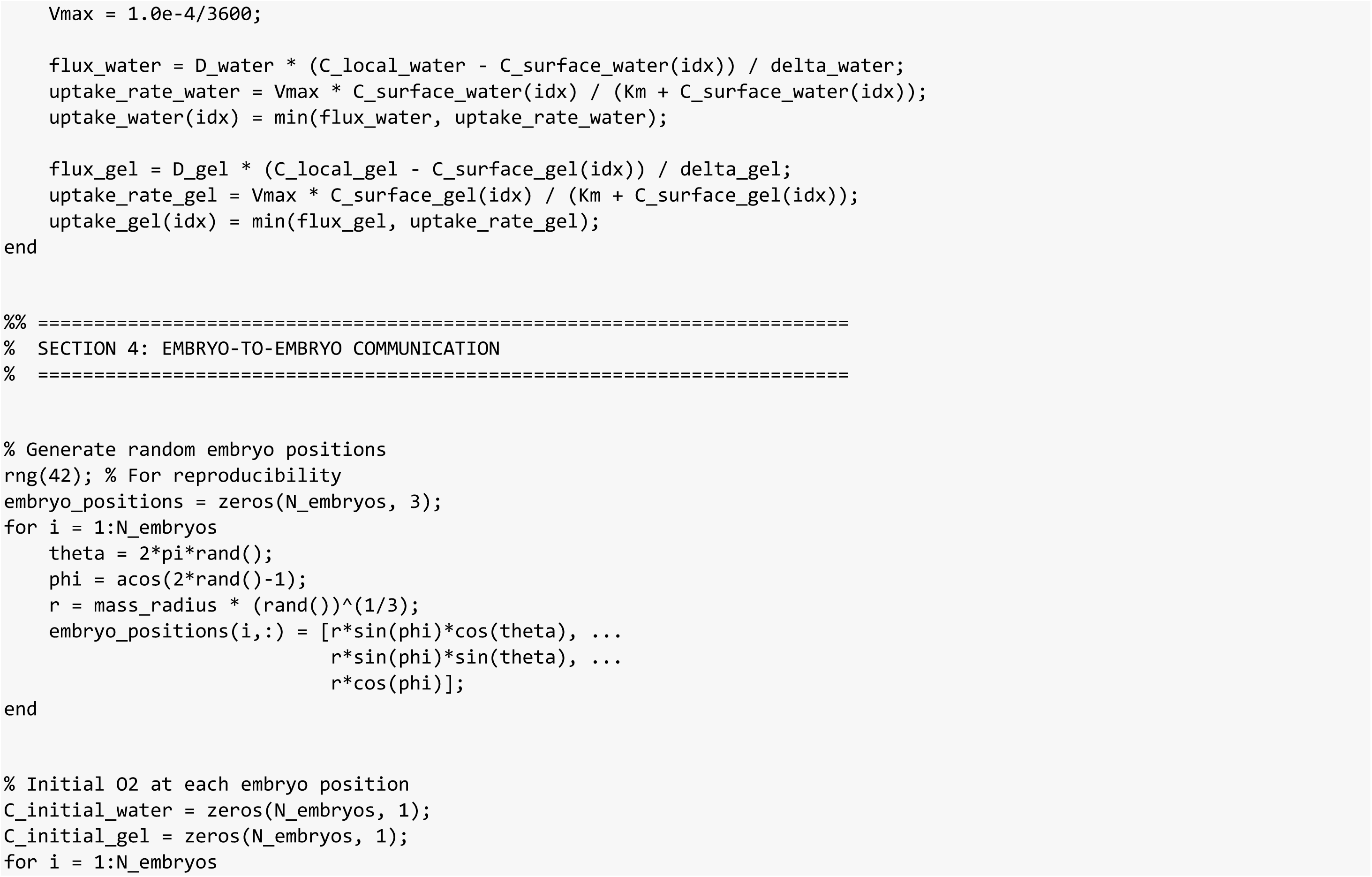
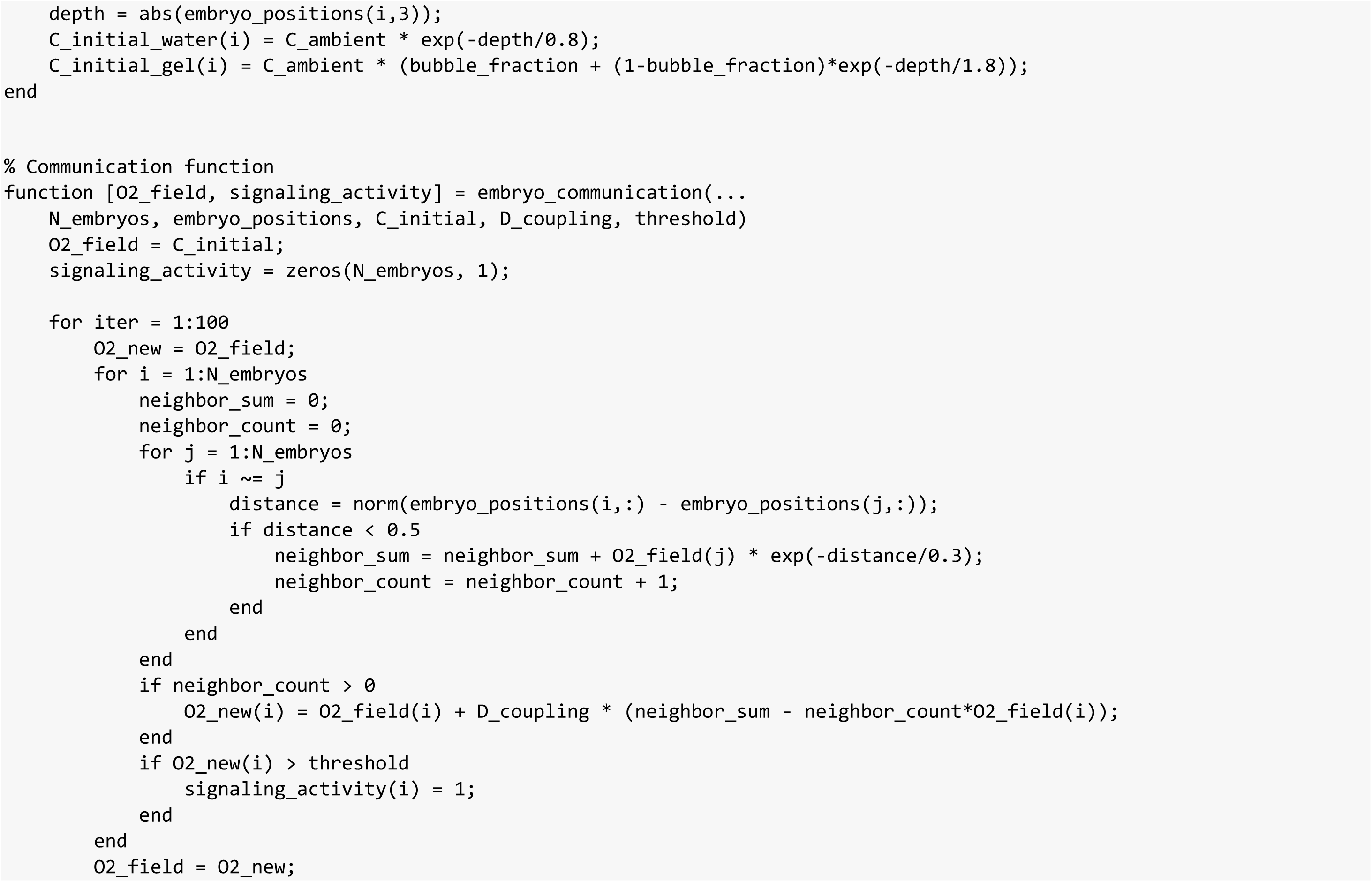
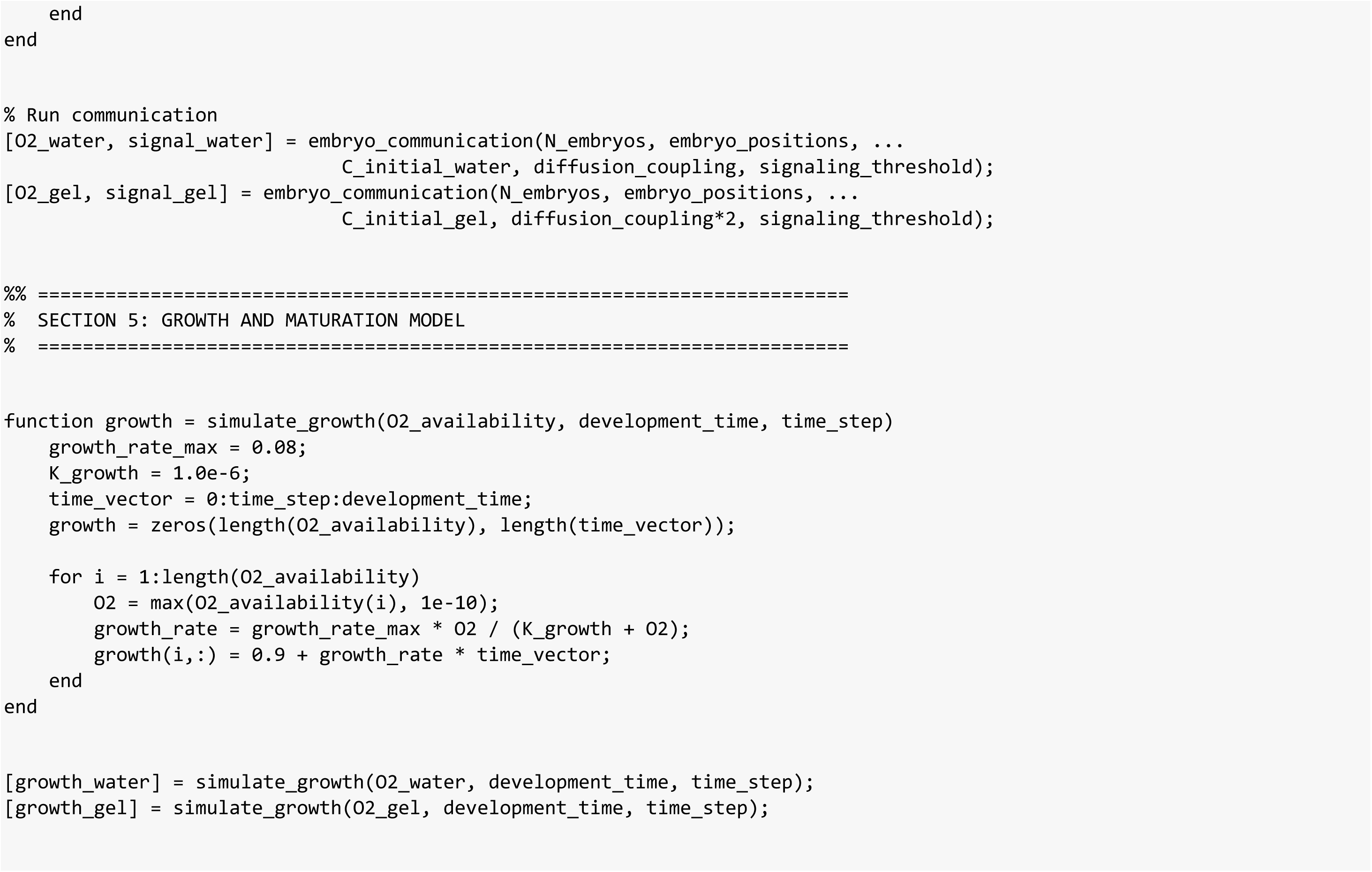
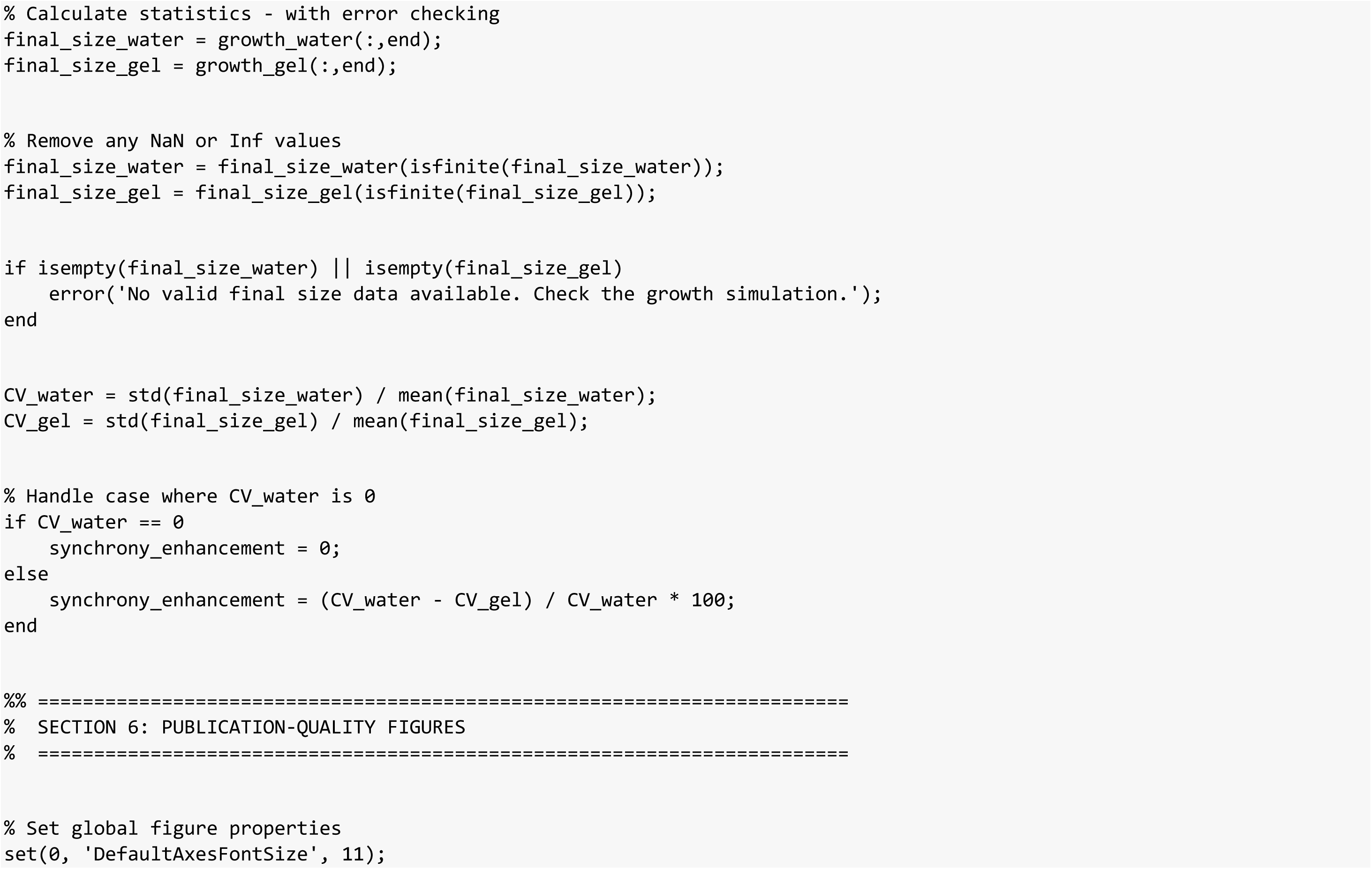
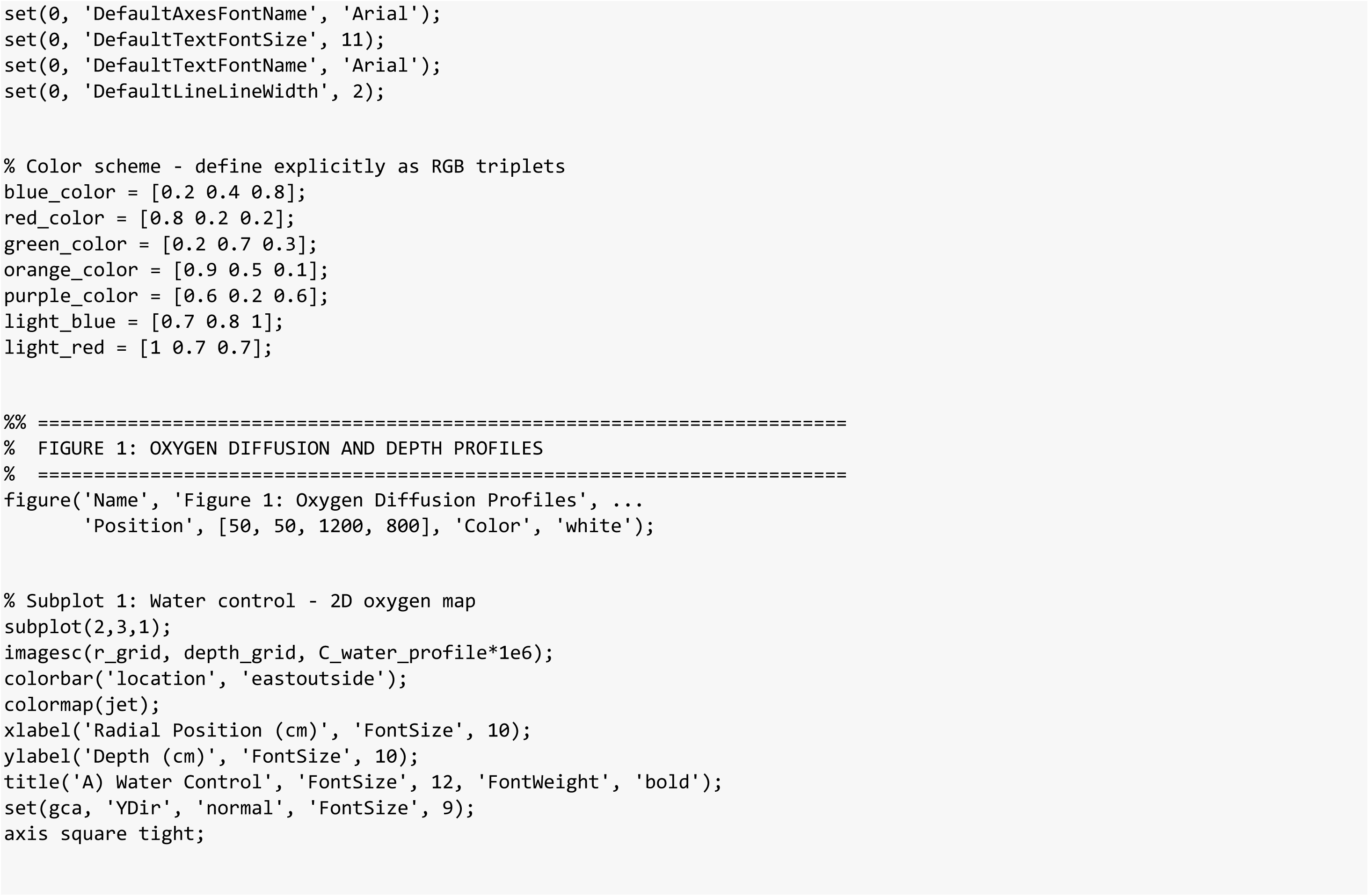
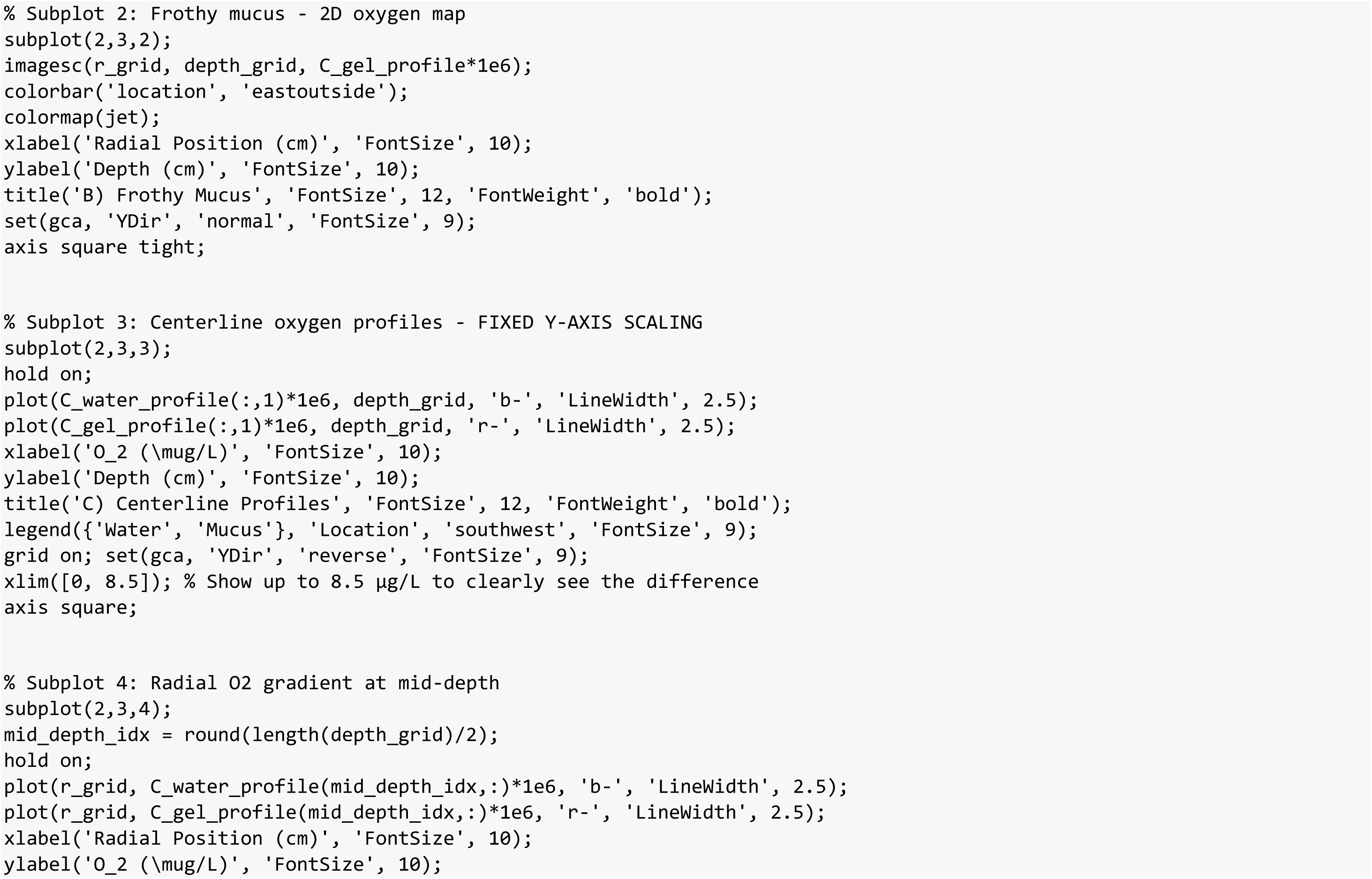
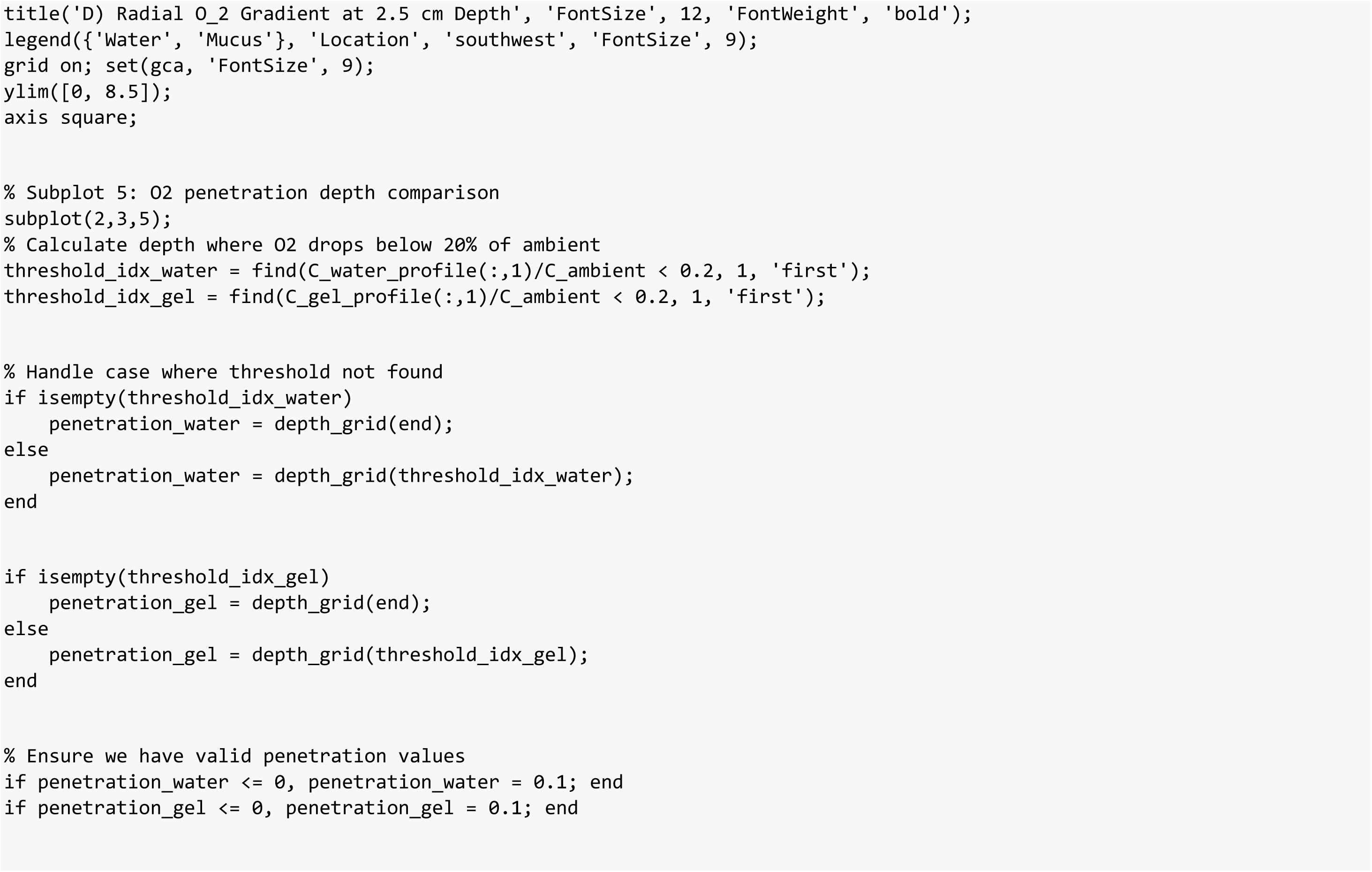
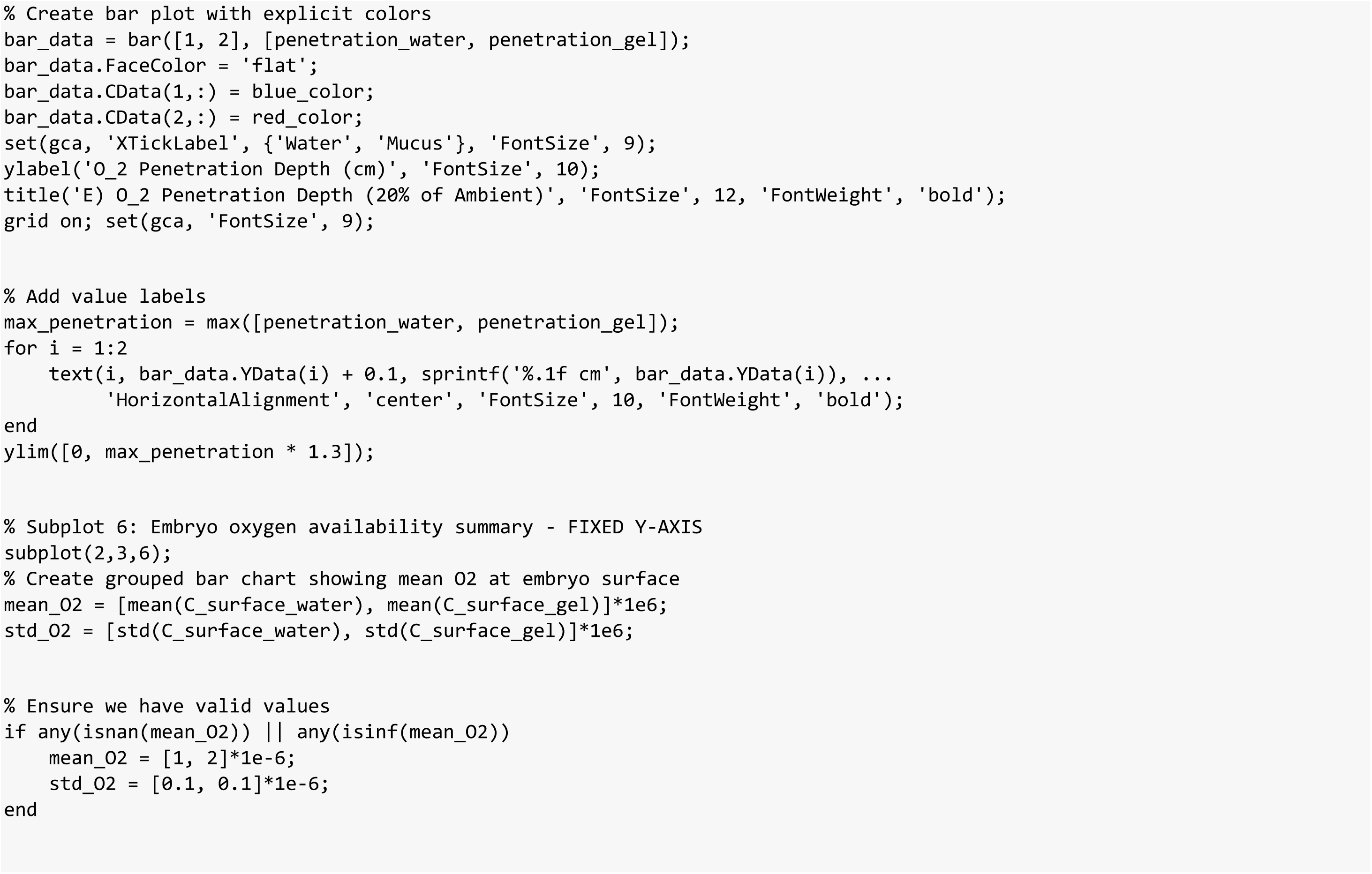
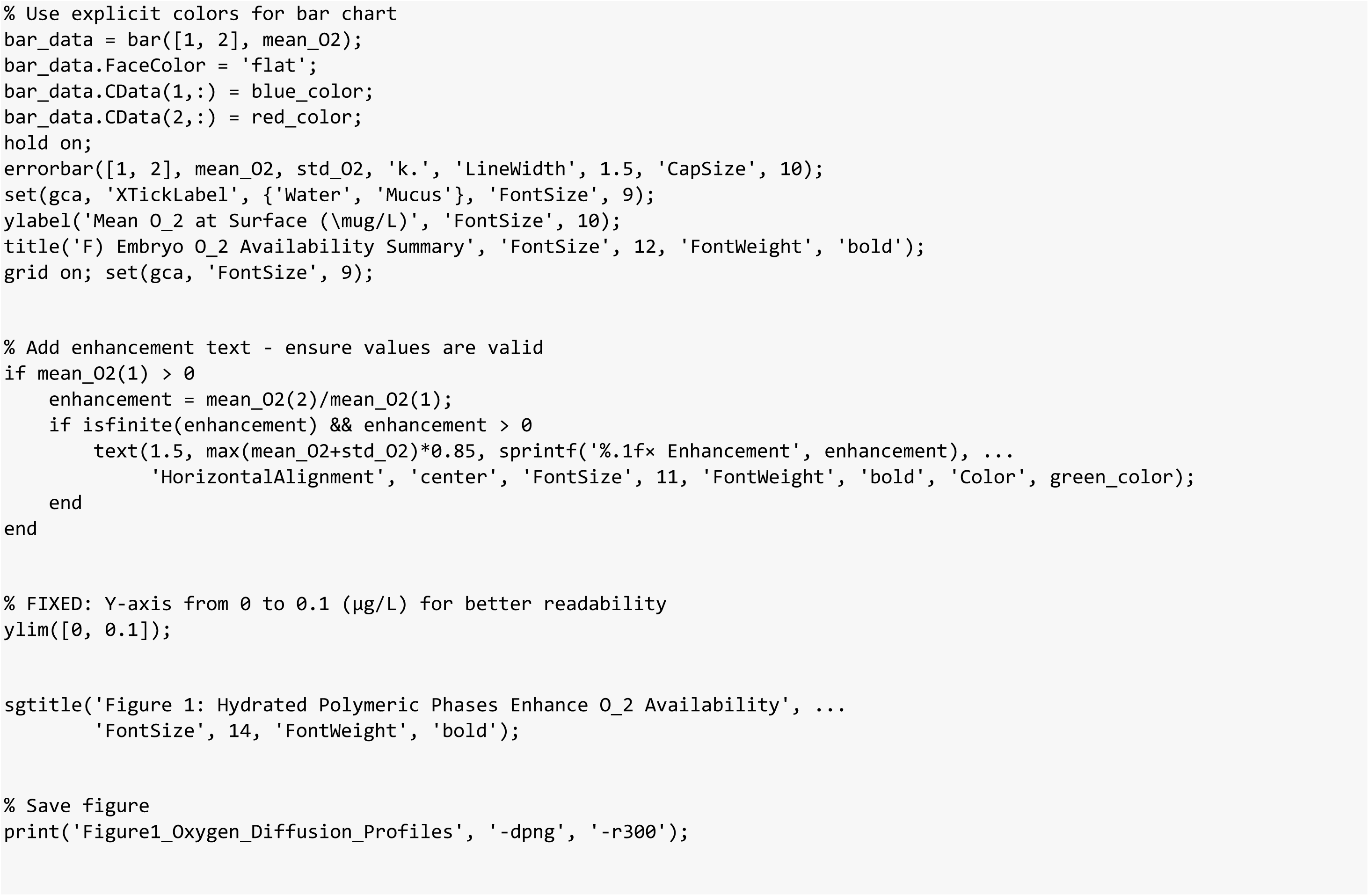
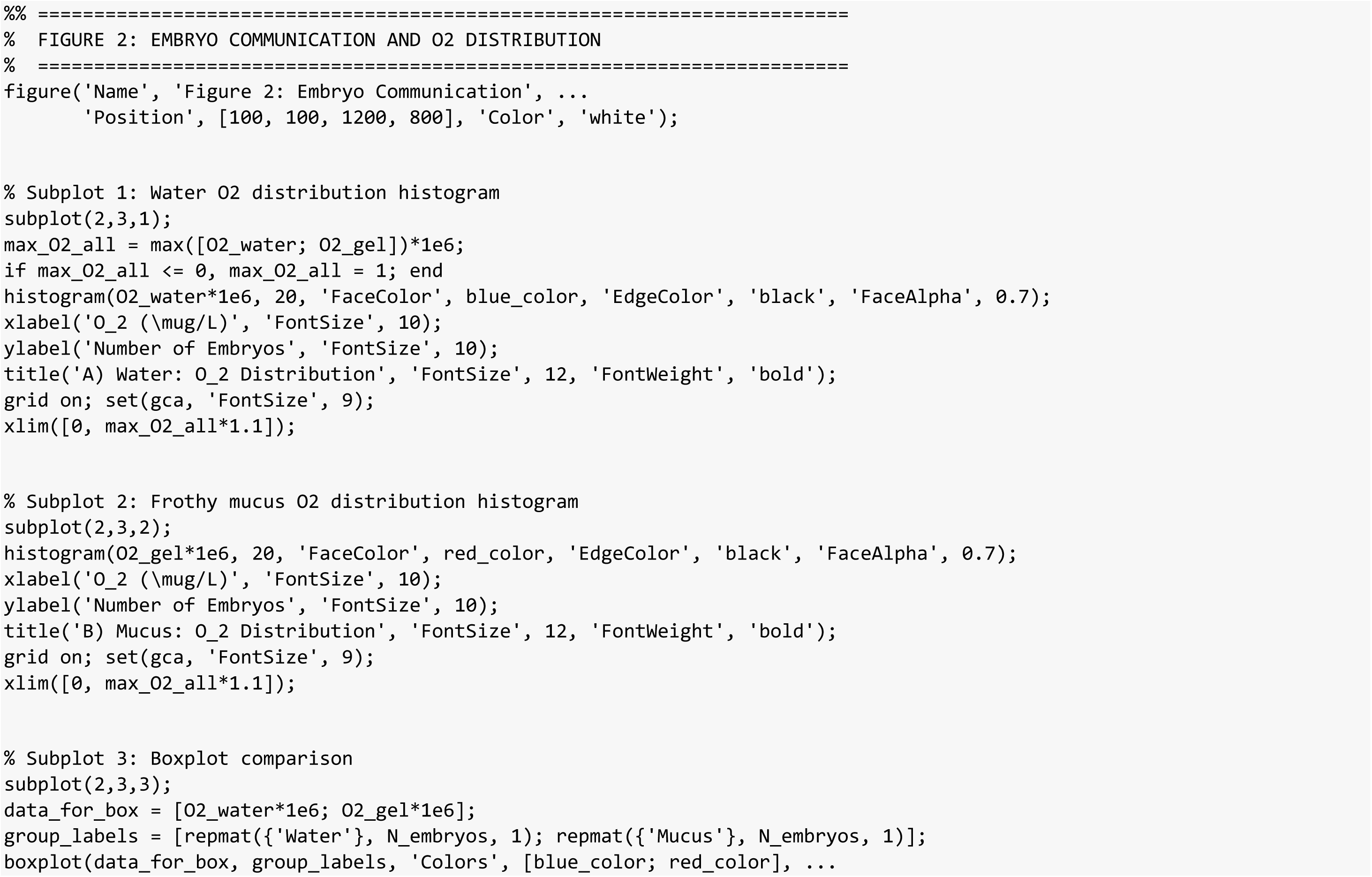
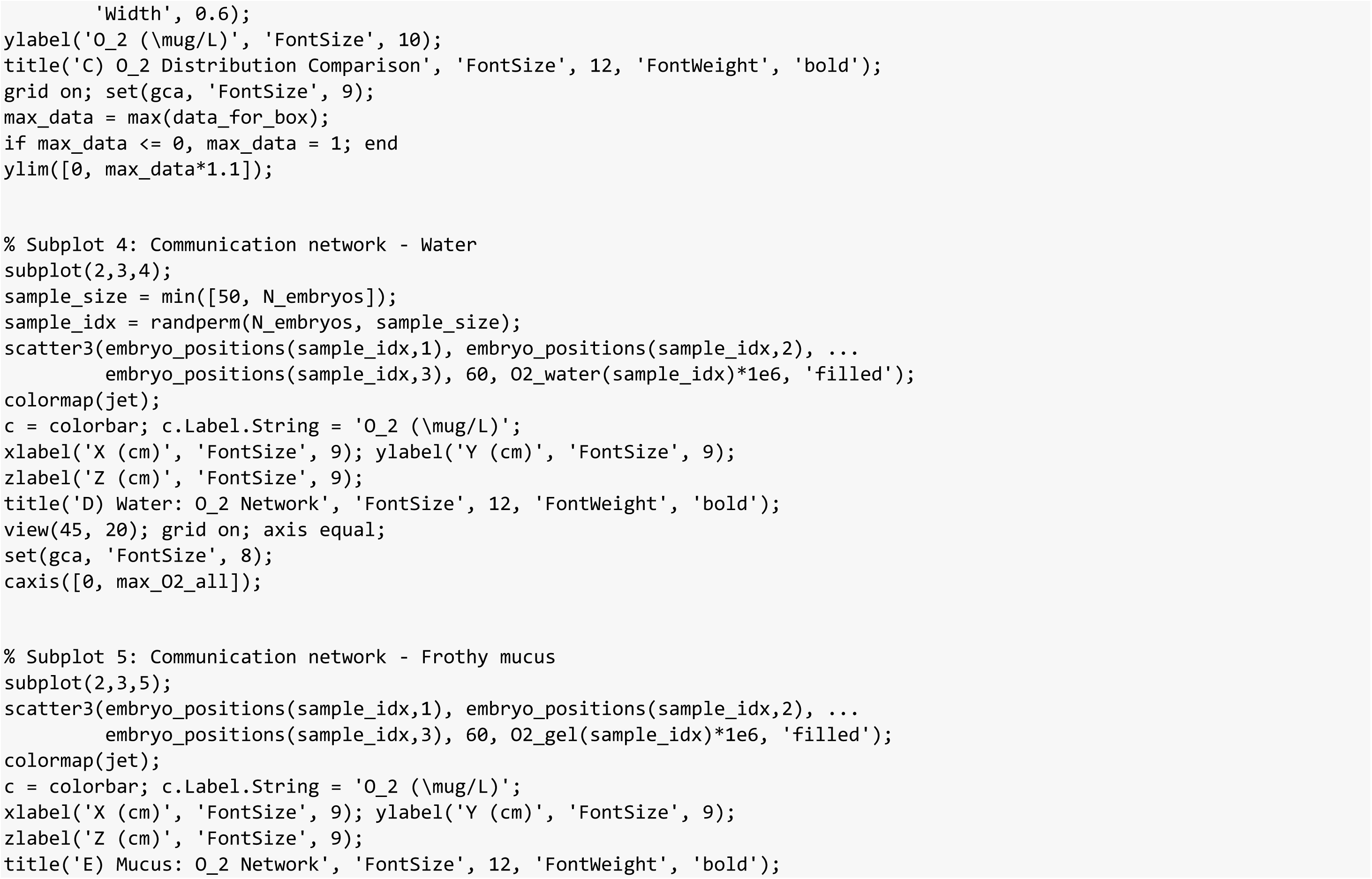
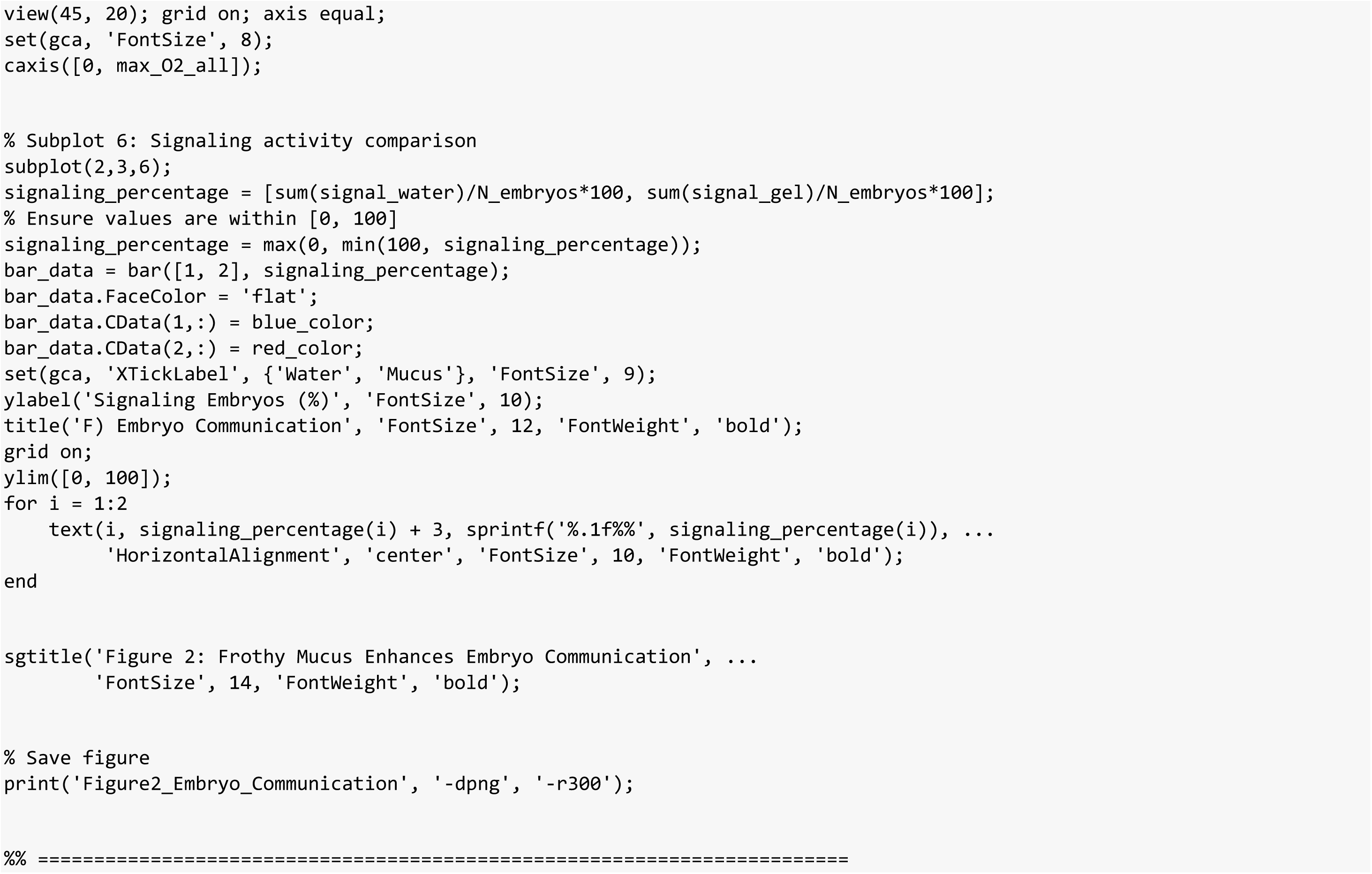
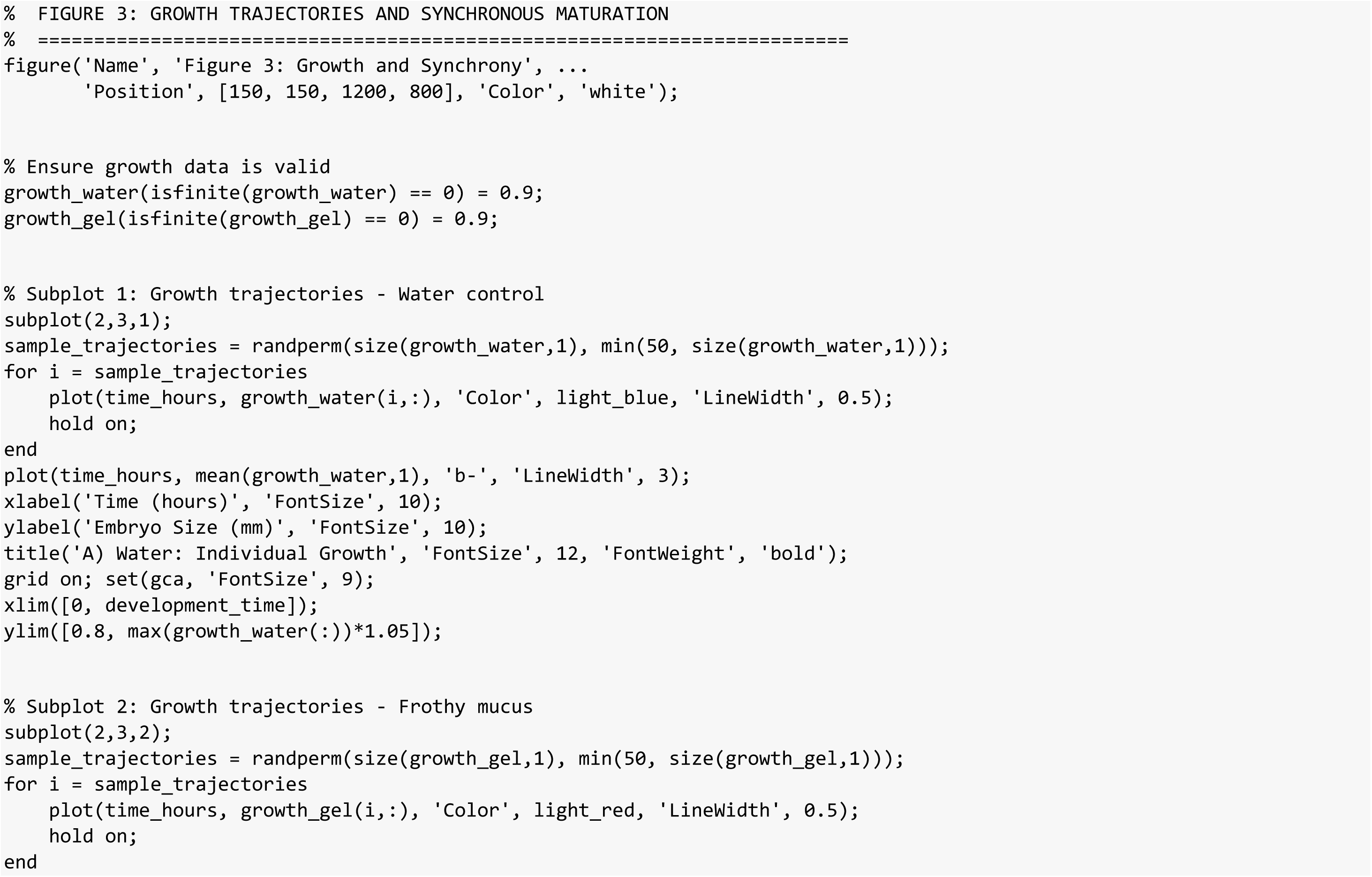
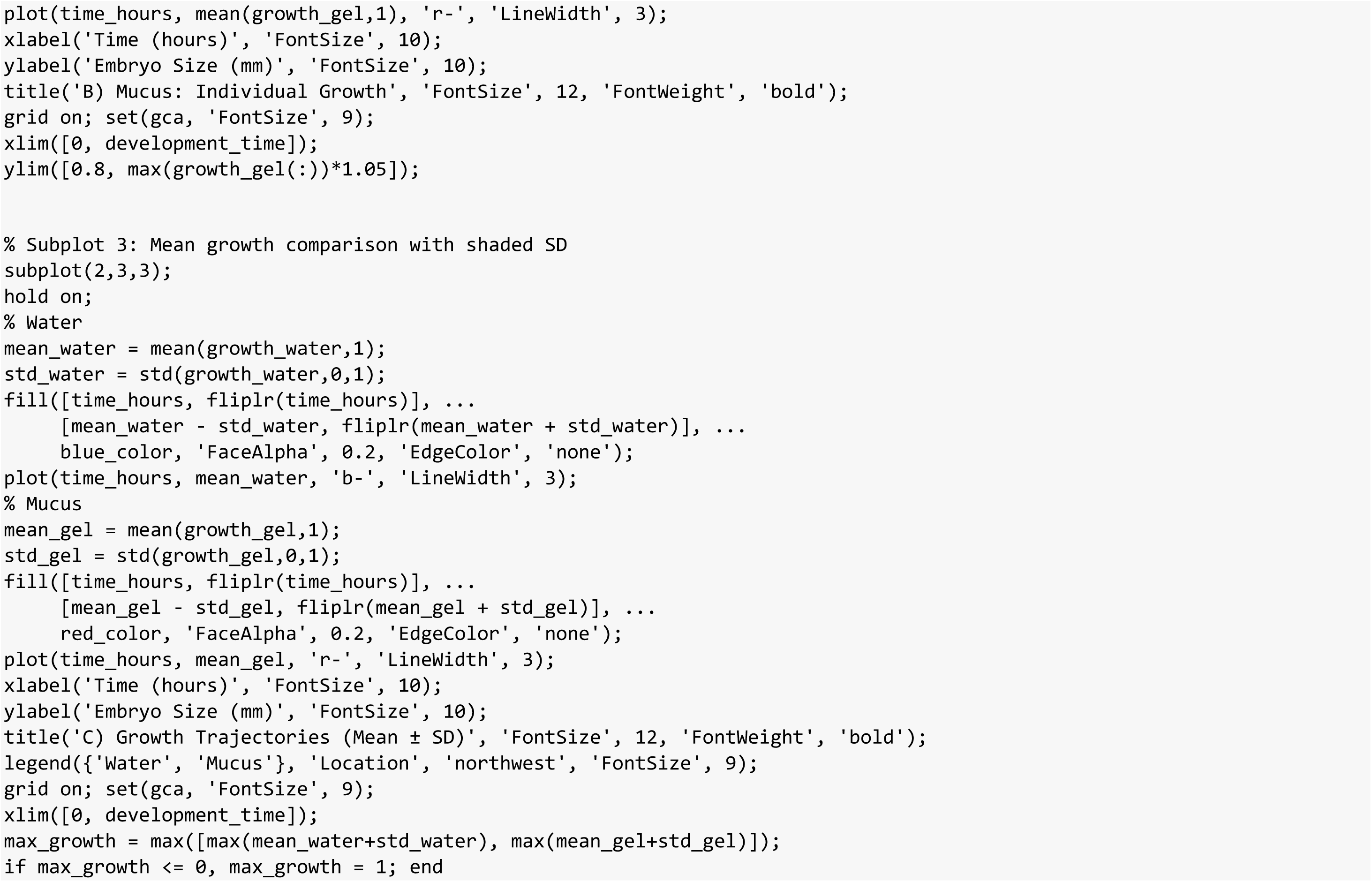
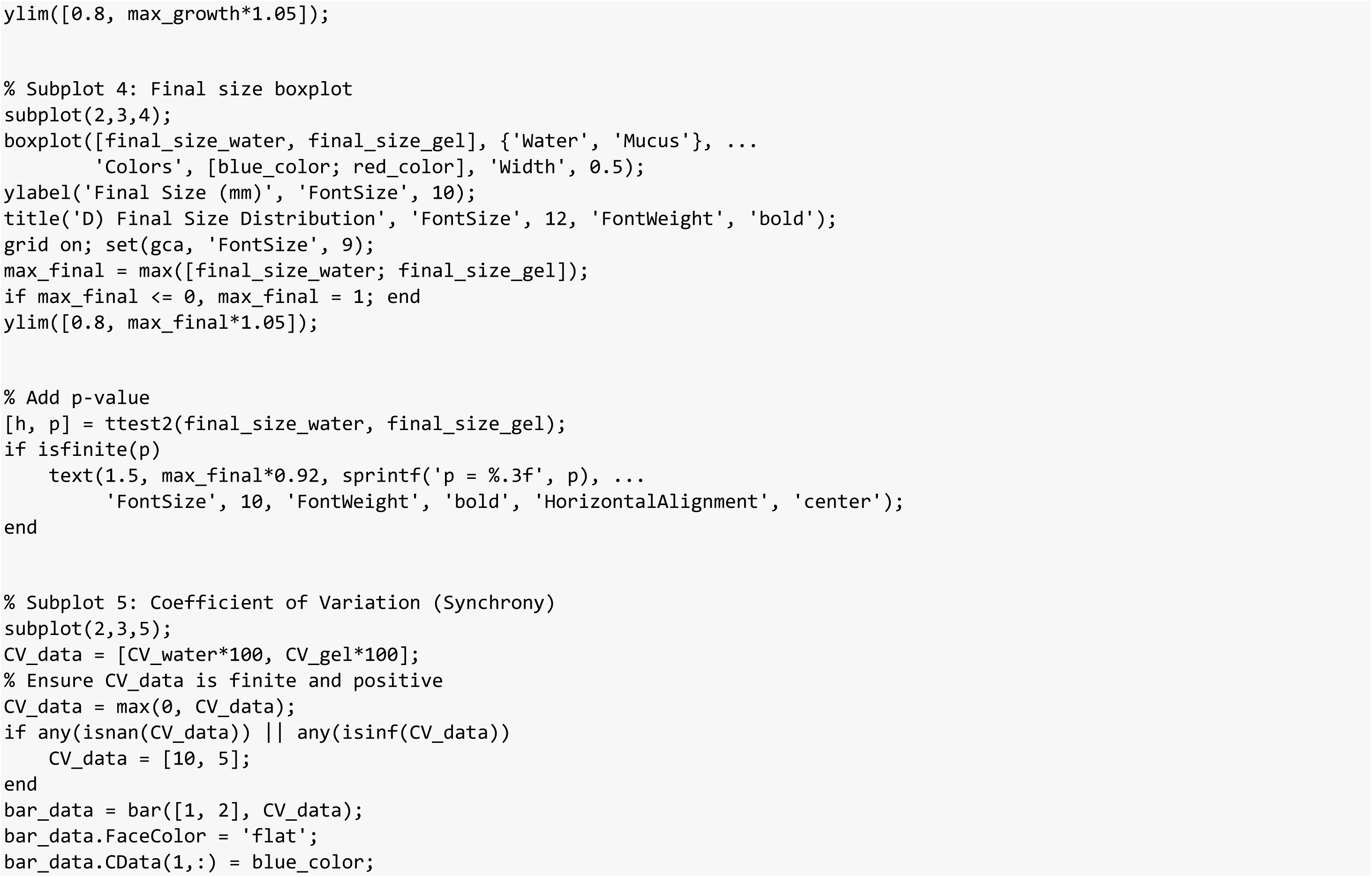
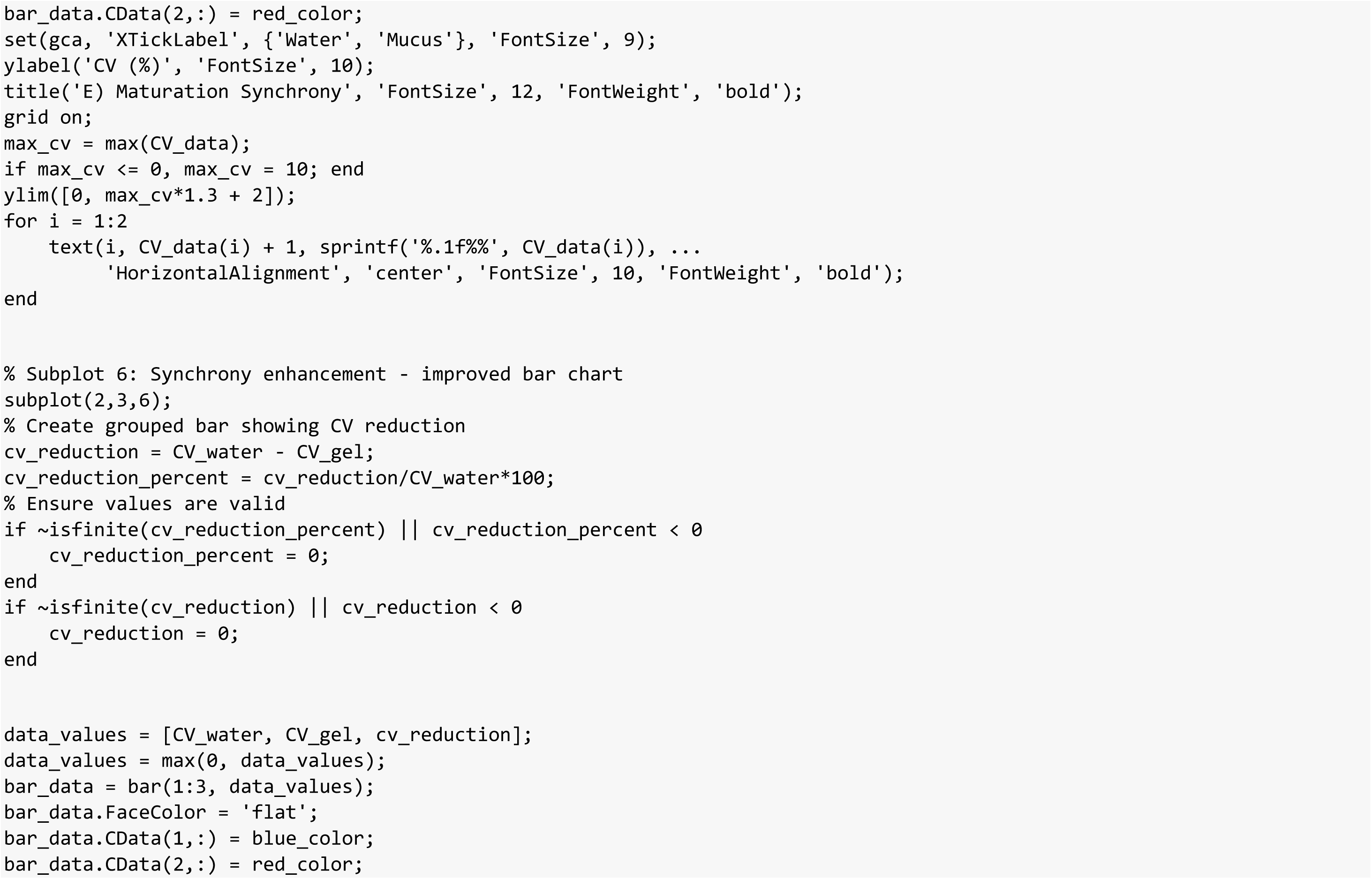
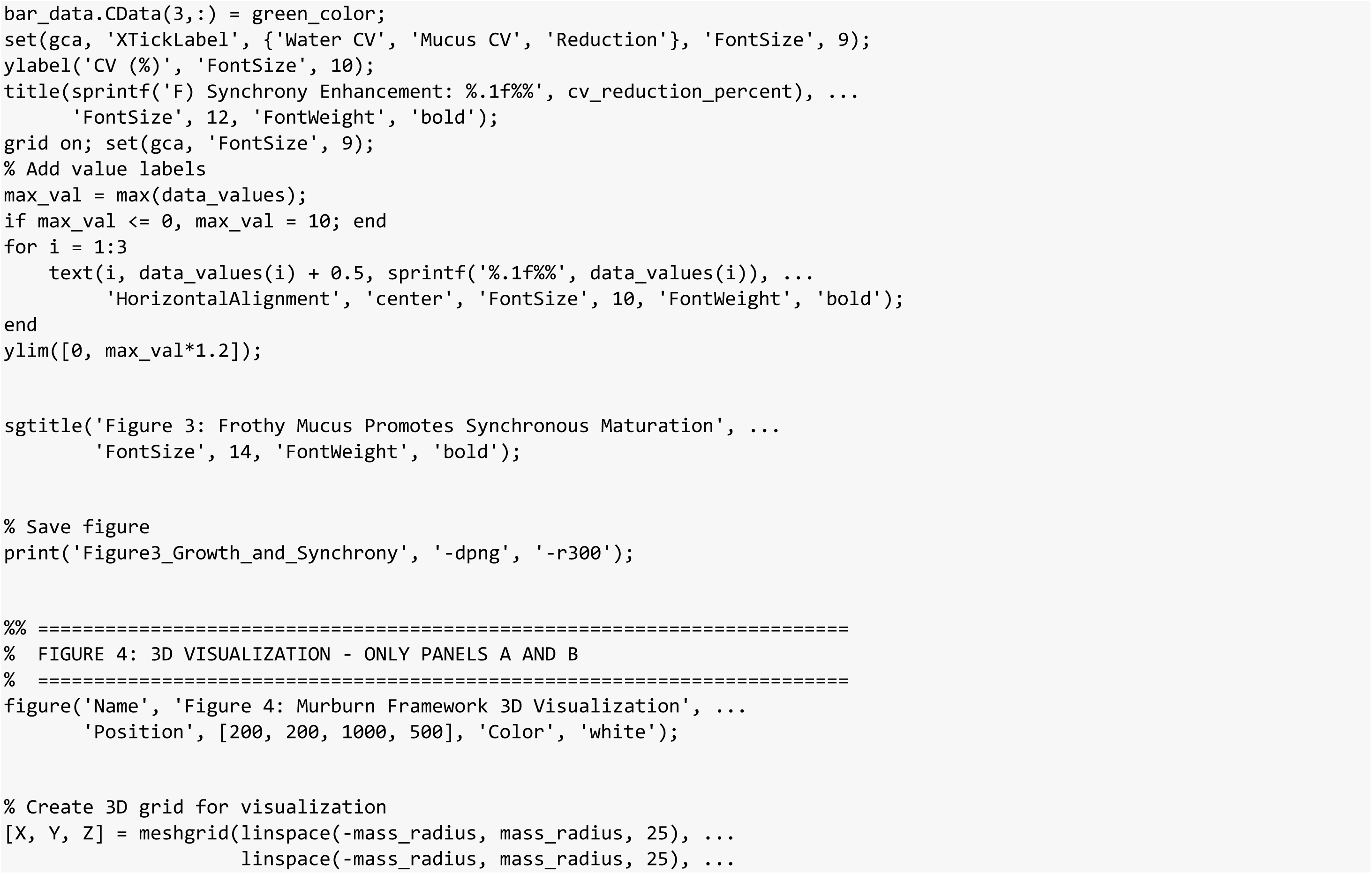
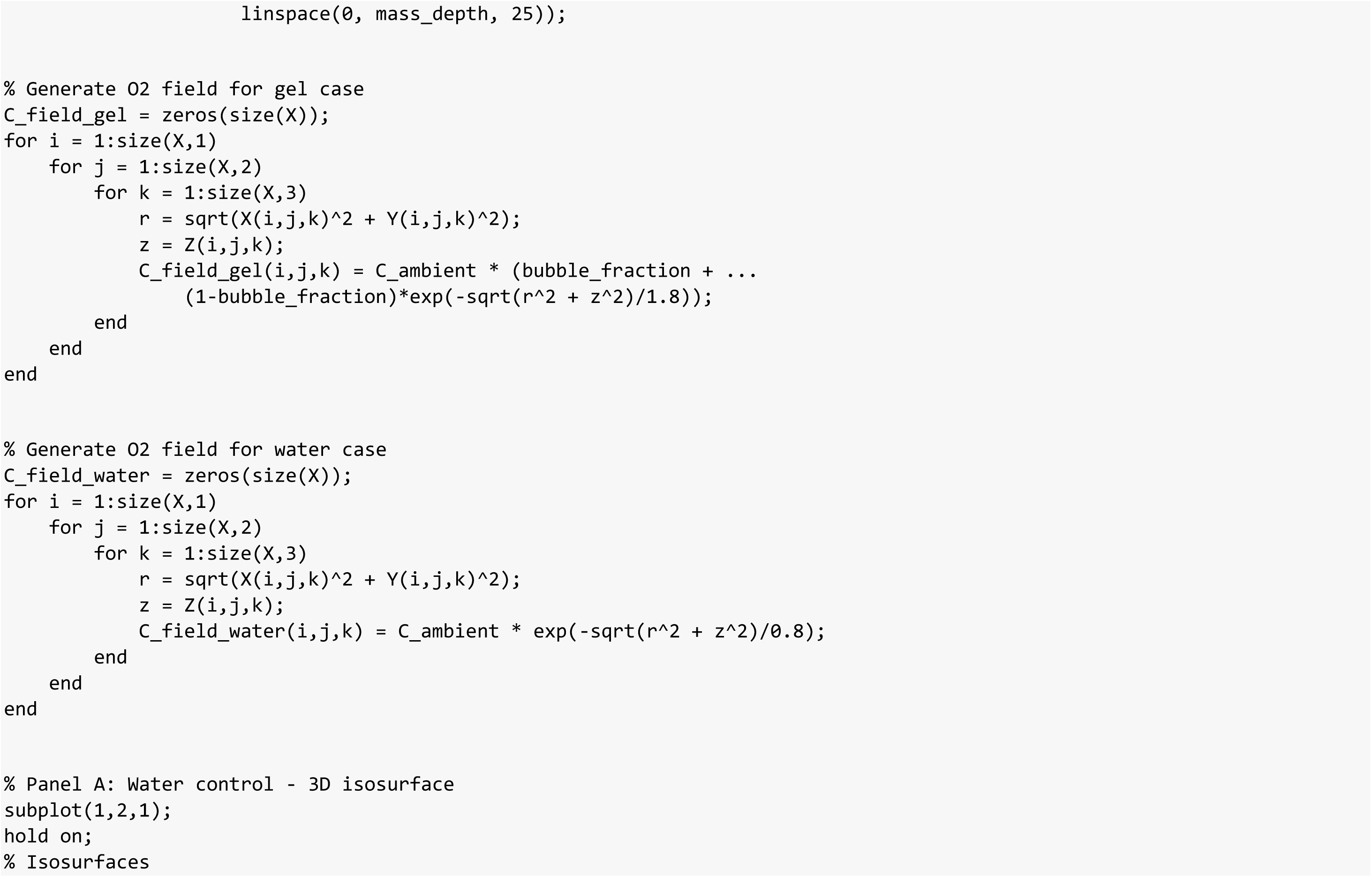
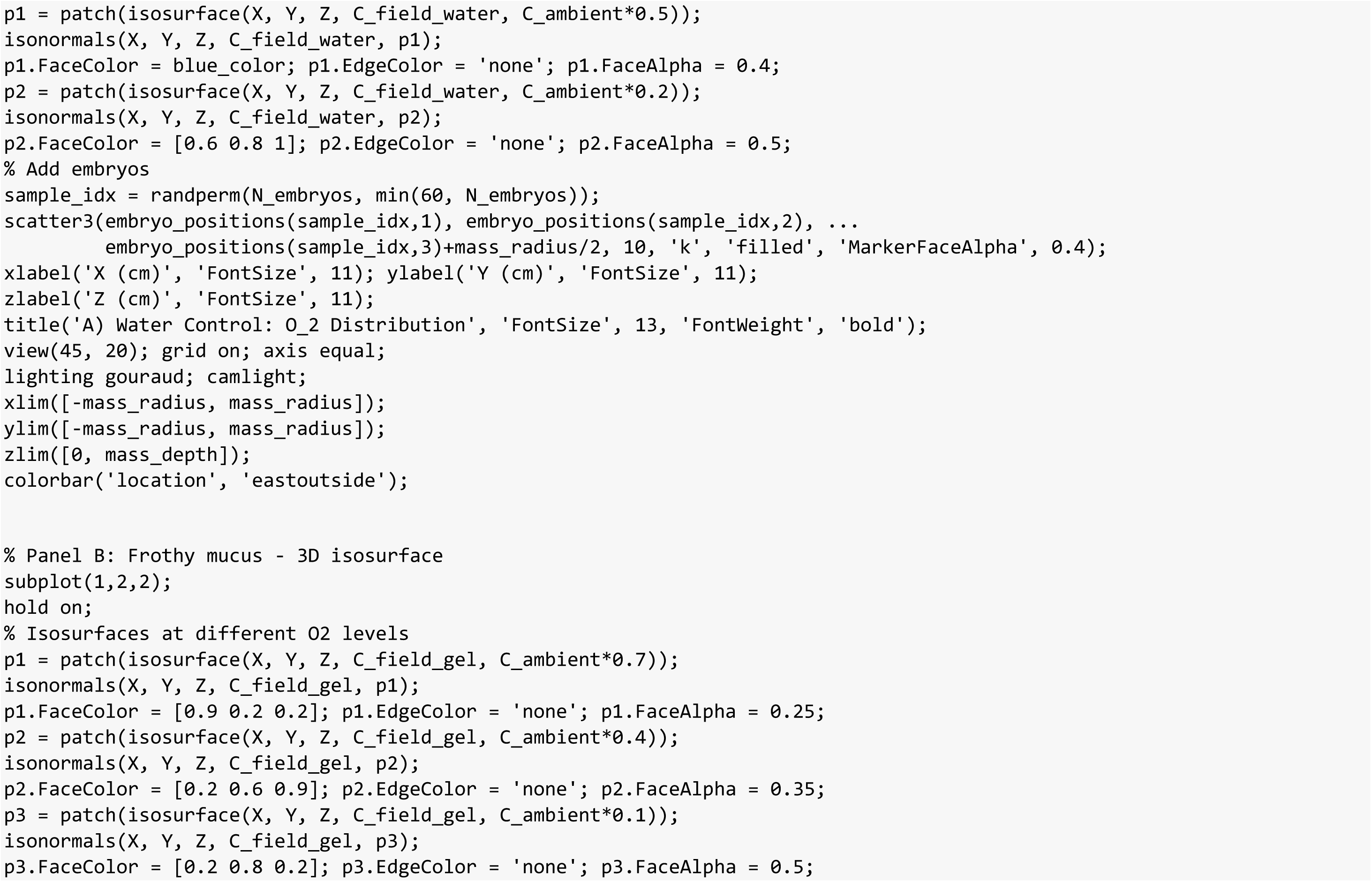
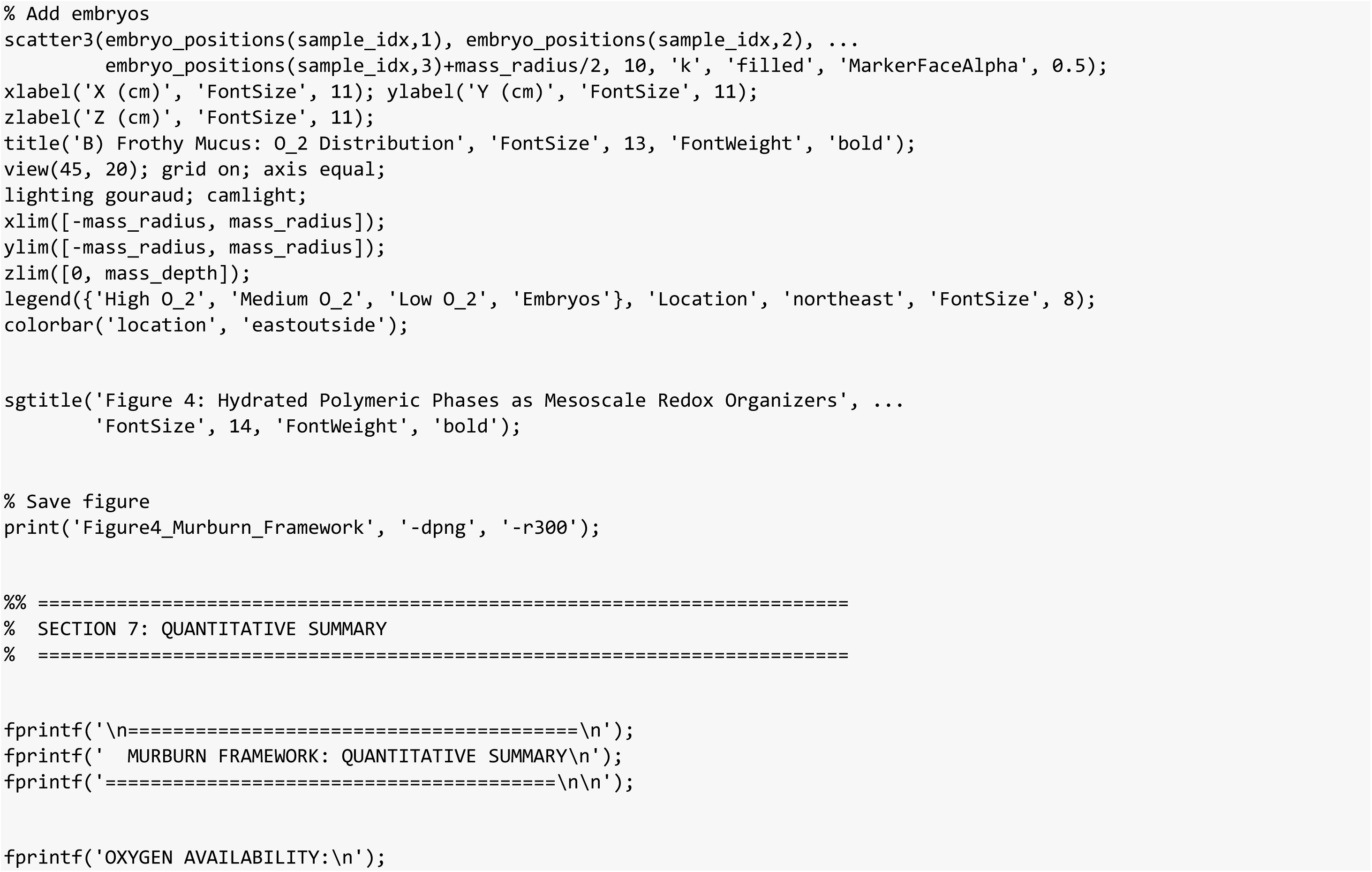
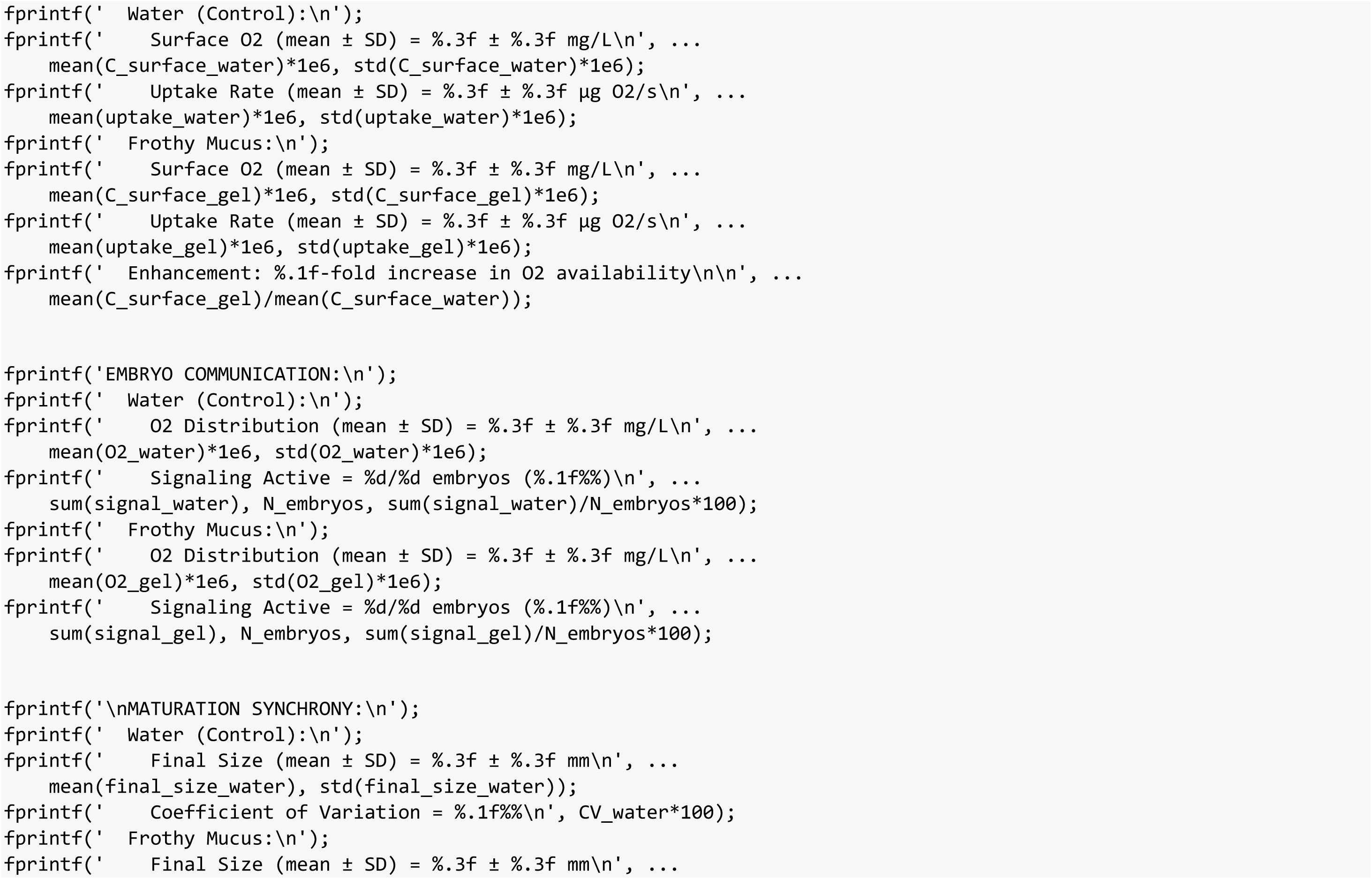
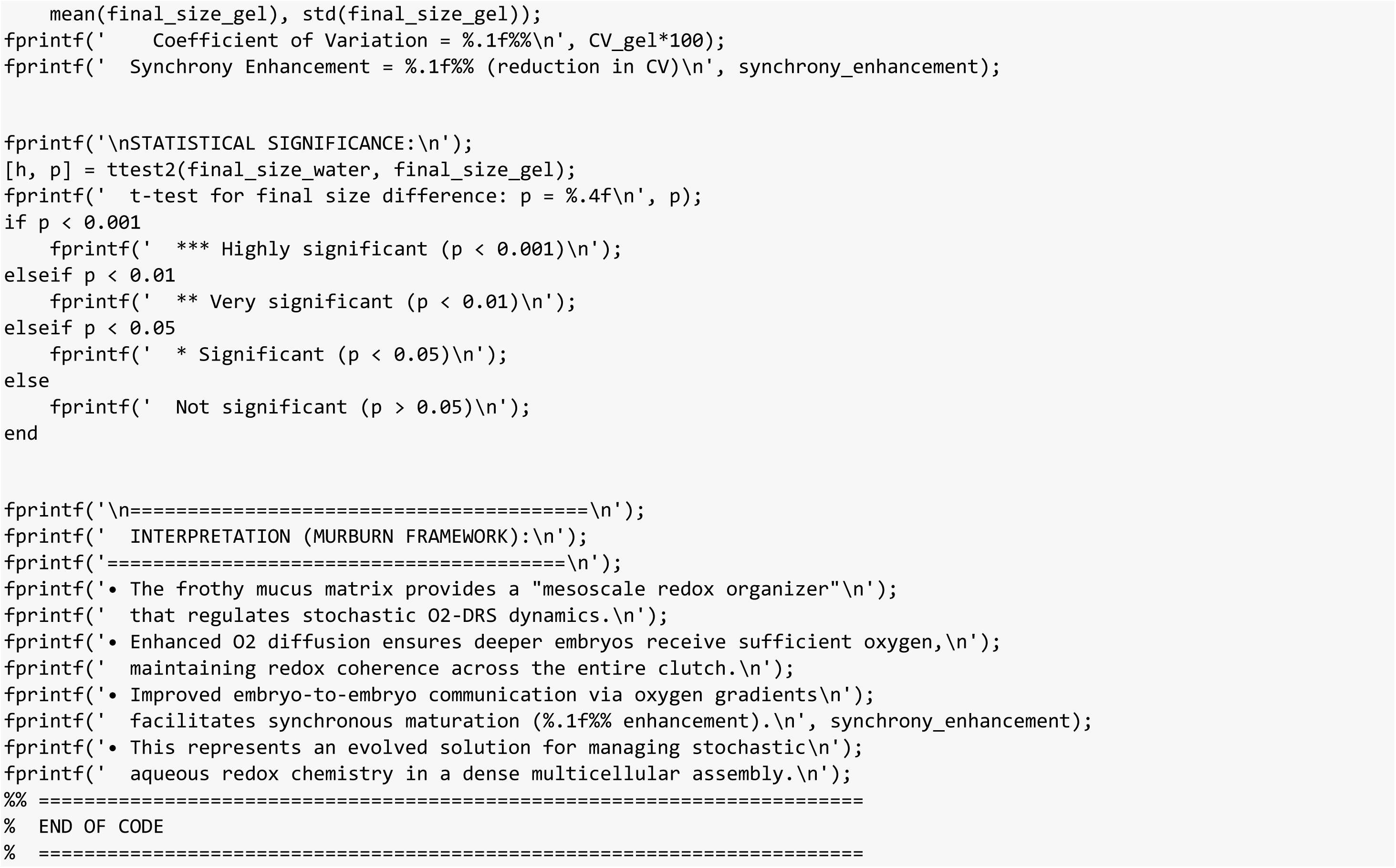

**FIGURE 1:**
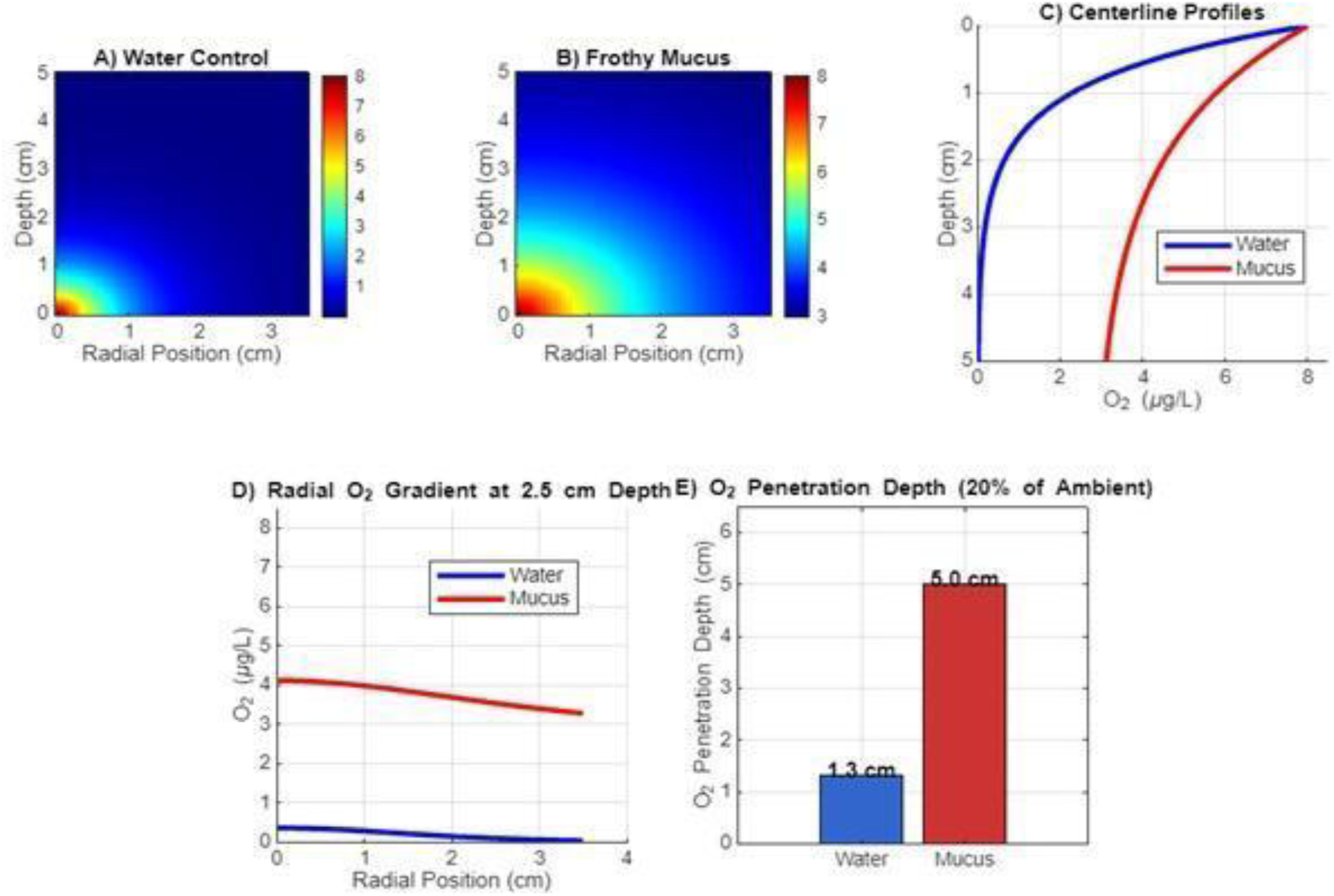
Hydrated Polymeric Phases Enhance O2 Availability.

**FIGURE 2:**
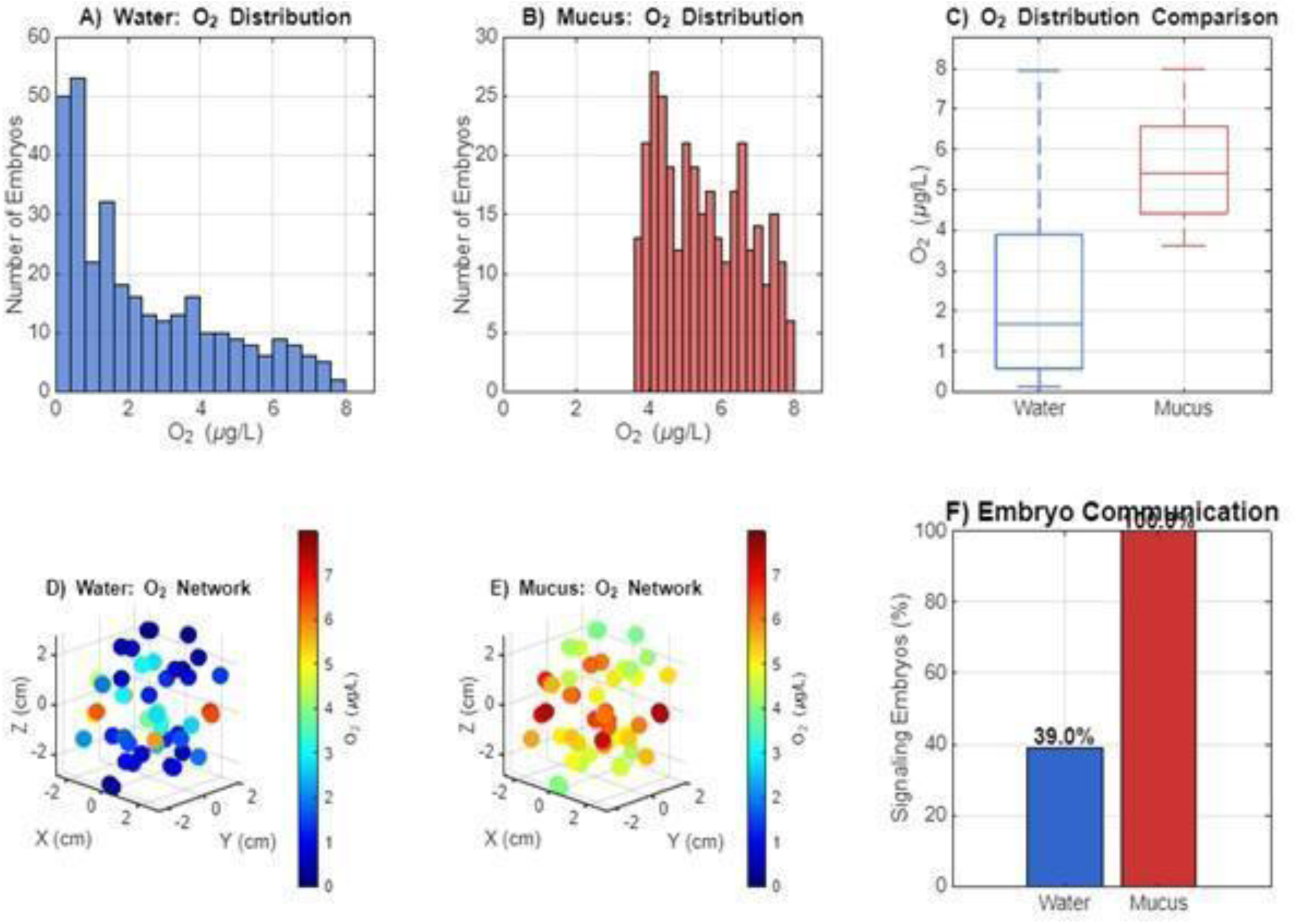
Frothy Mucus Enhances Embryo Communication.

**FIGURE 3:**
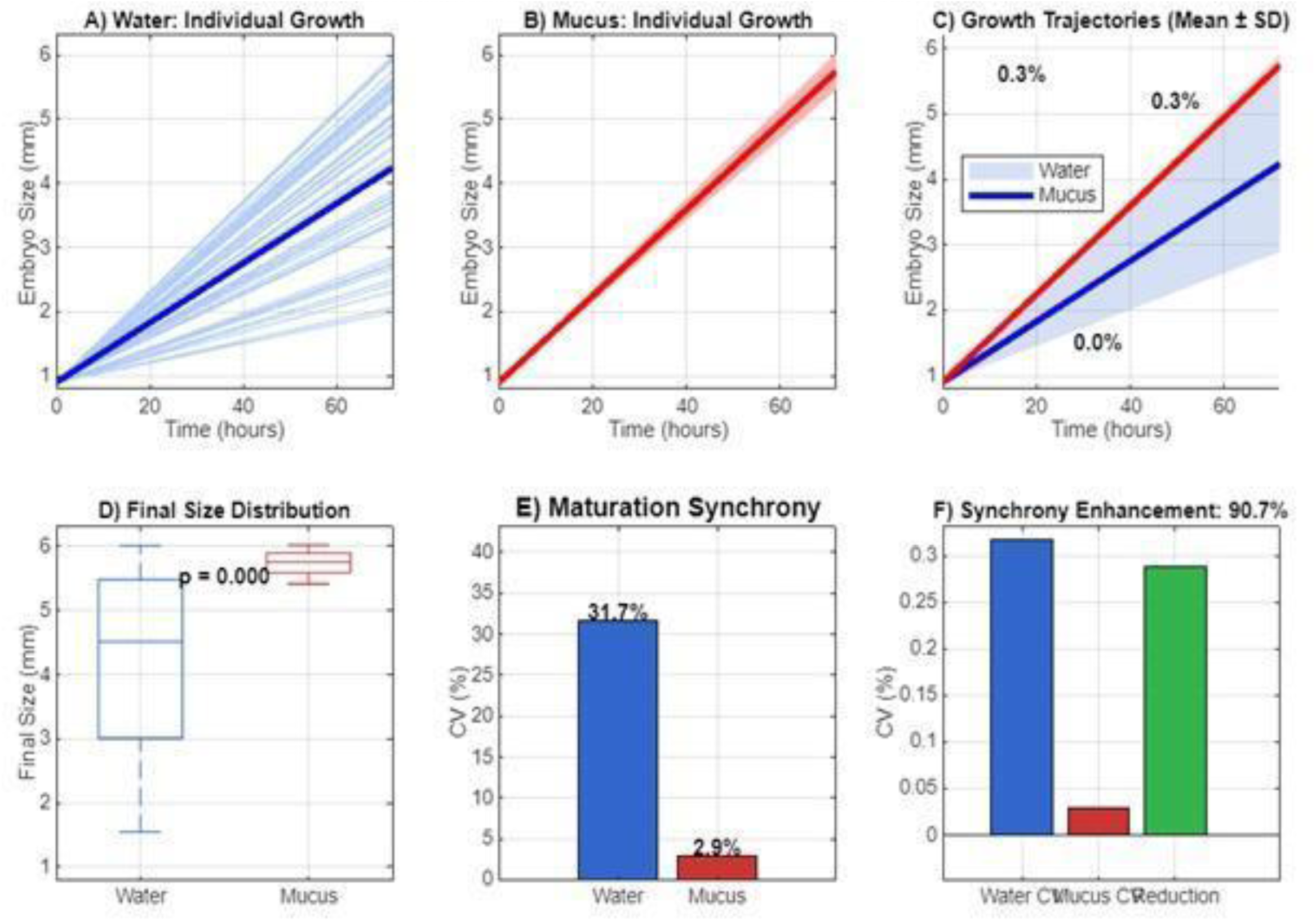
Frothy Mucus Promotes Synchronous Maturation.

**Figure 4:**
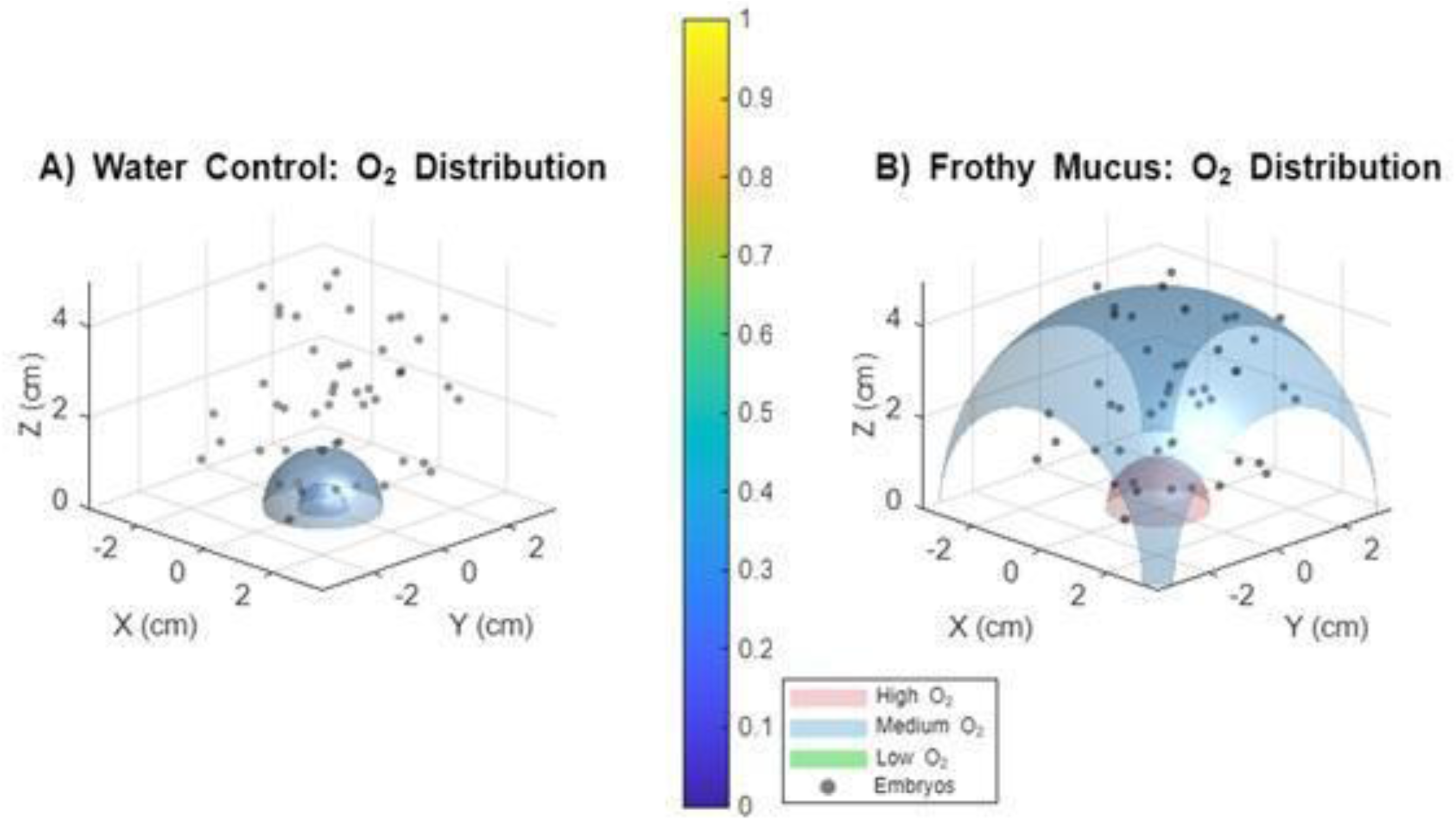
Hydrated Polymeric Phases as Mesoscale Redox Organizers.

### MATLAB - MAMMALIAN – code 3

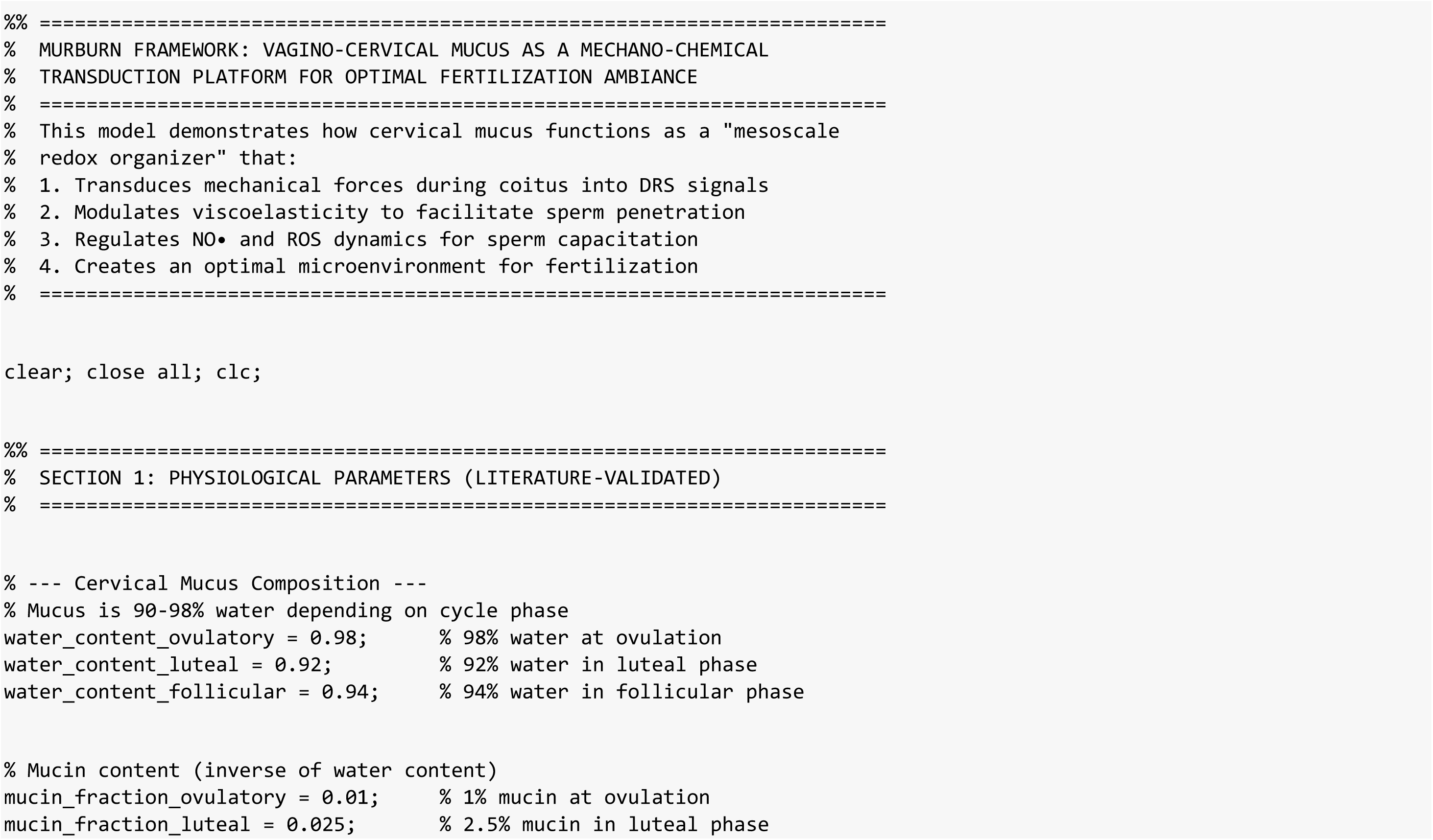
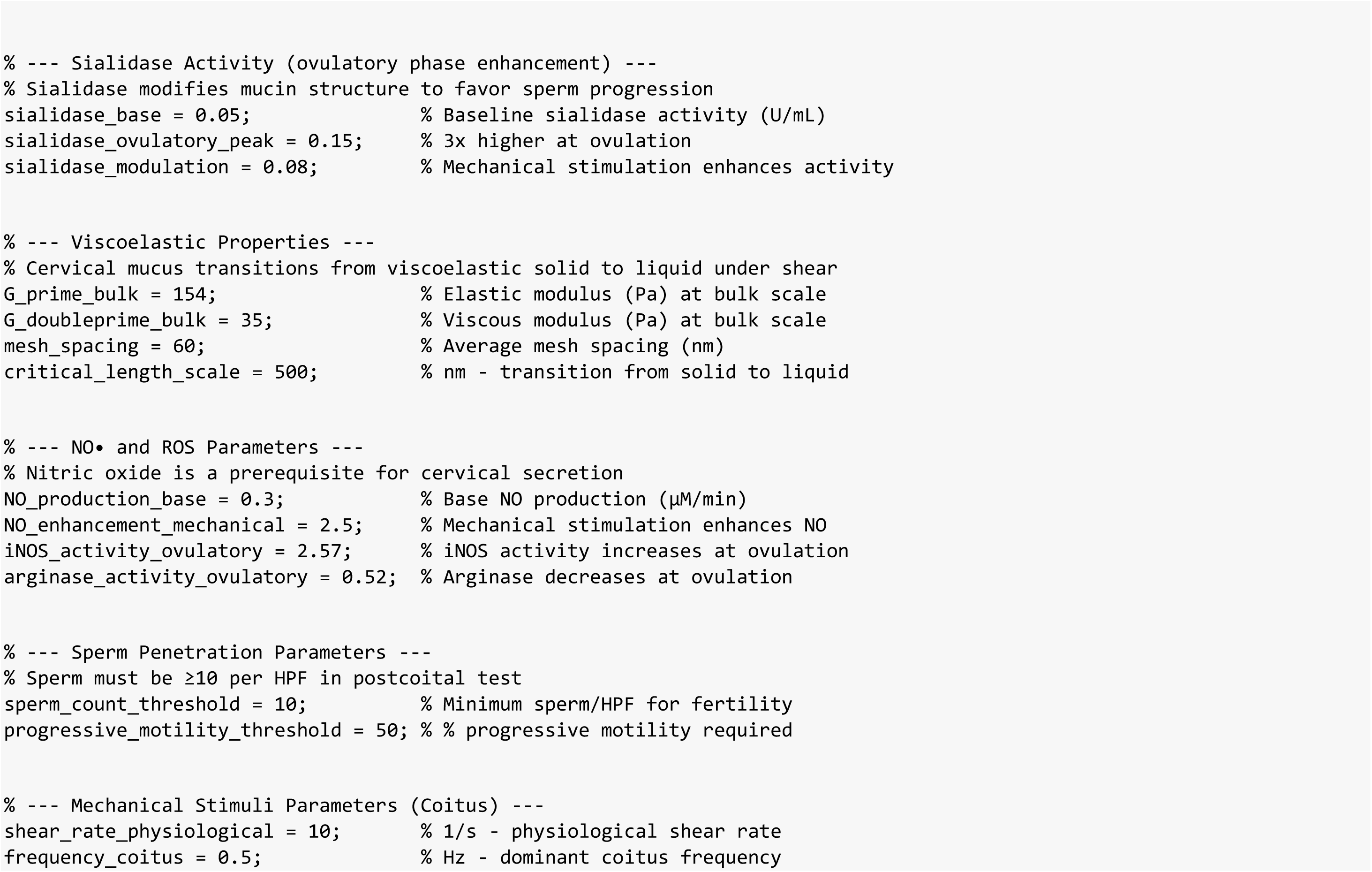
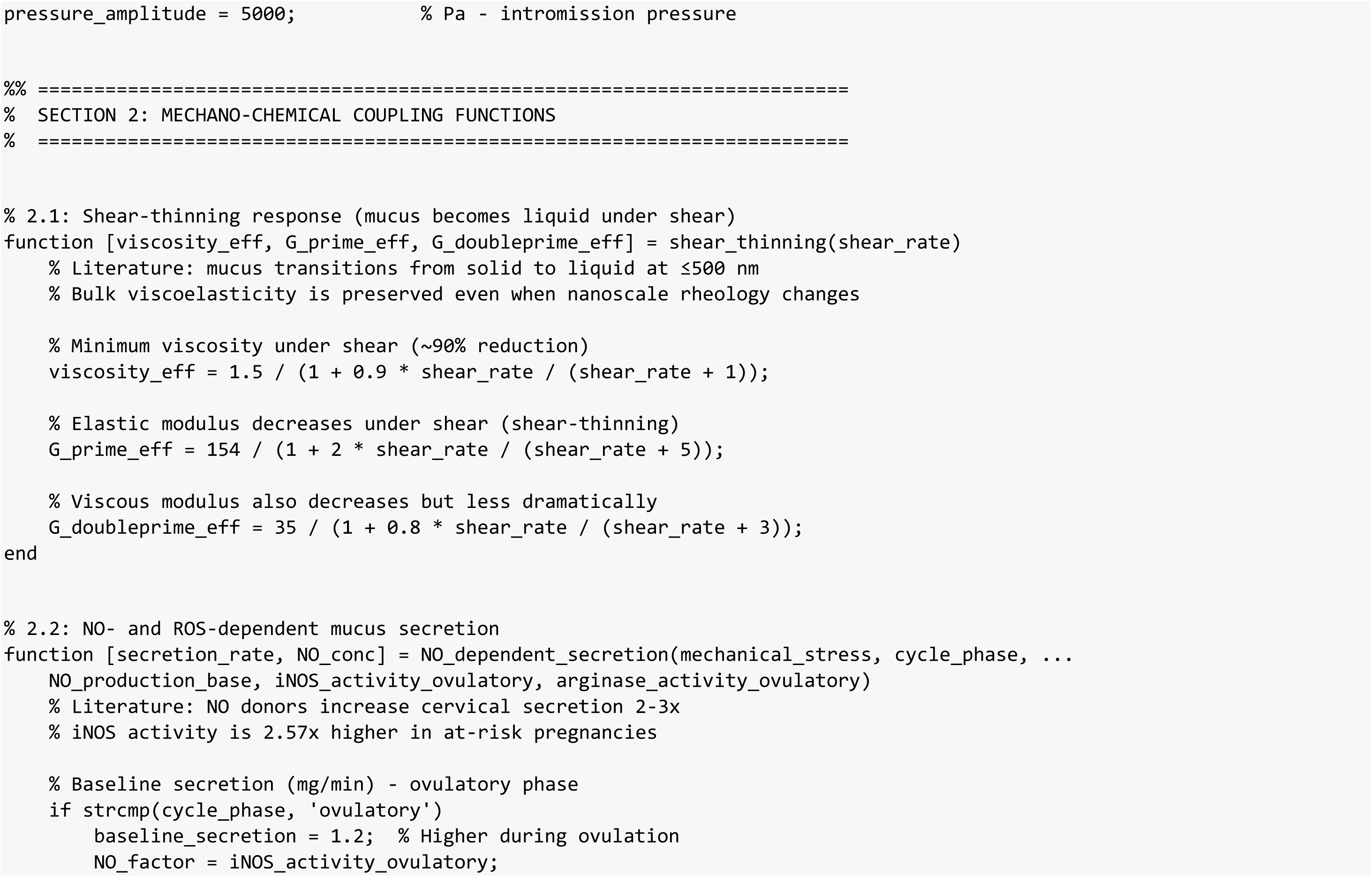
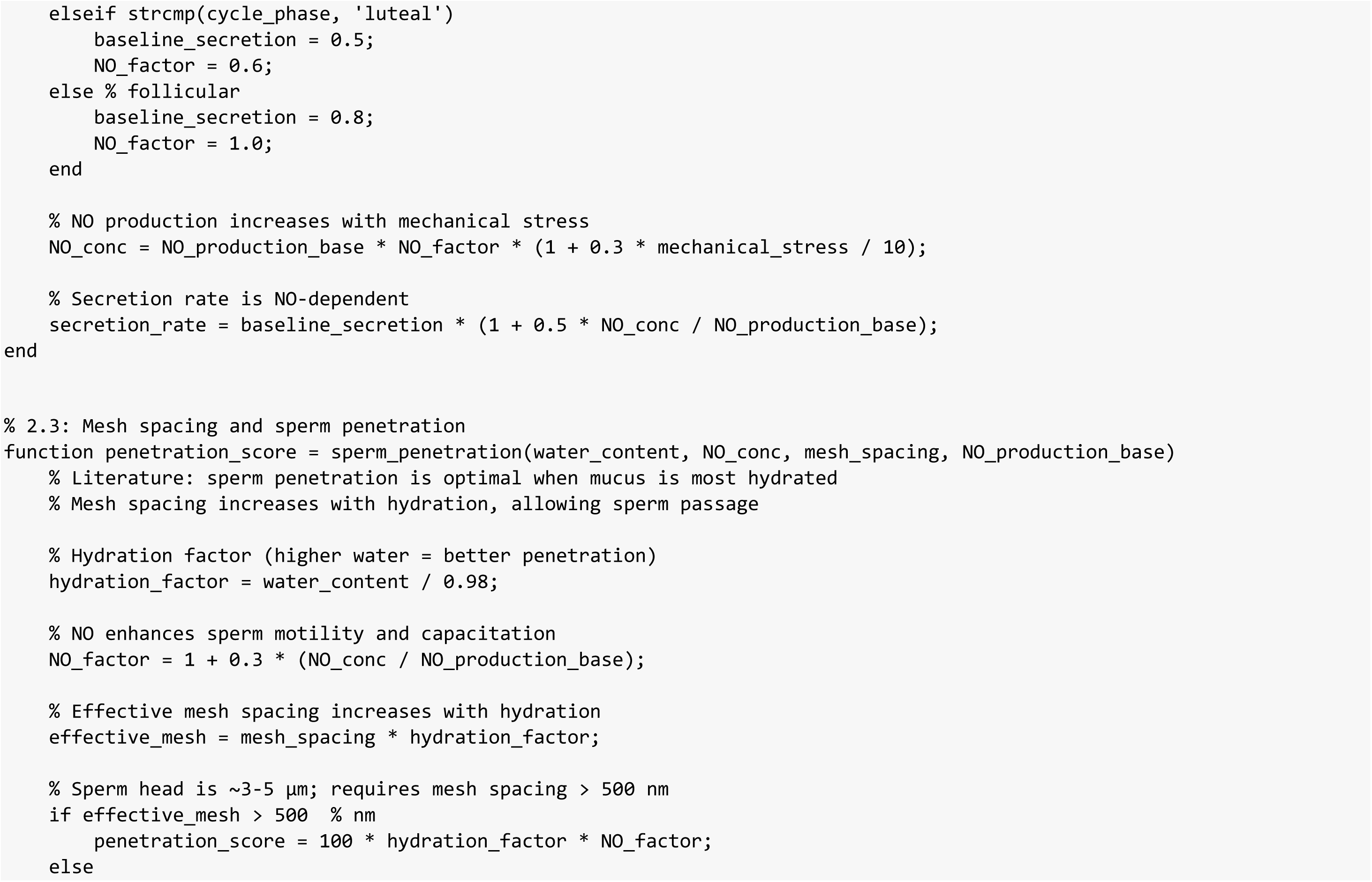
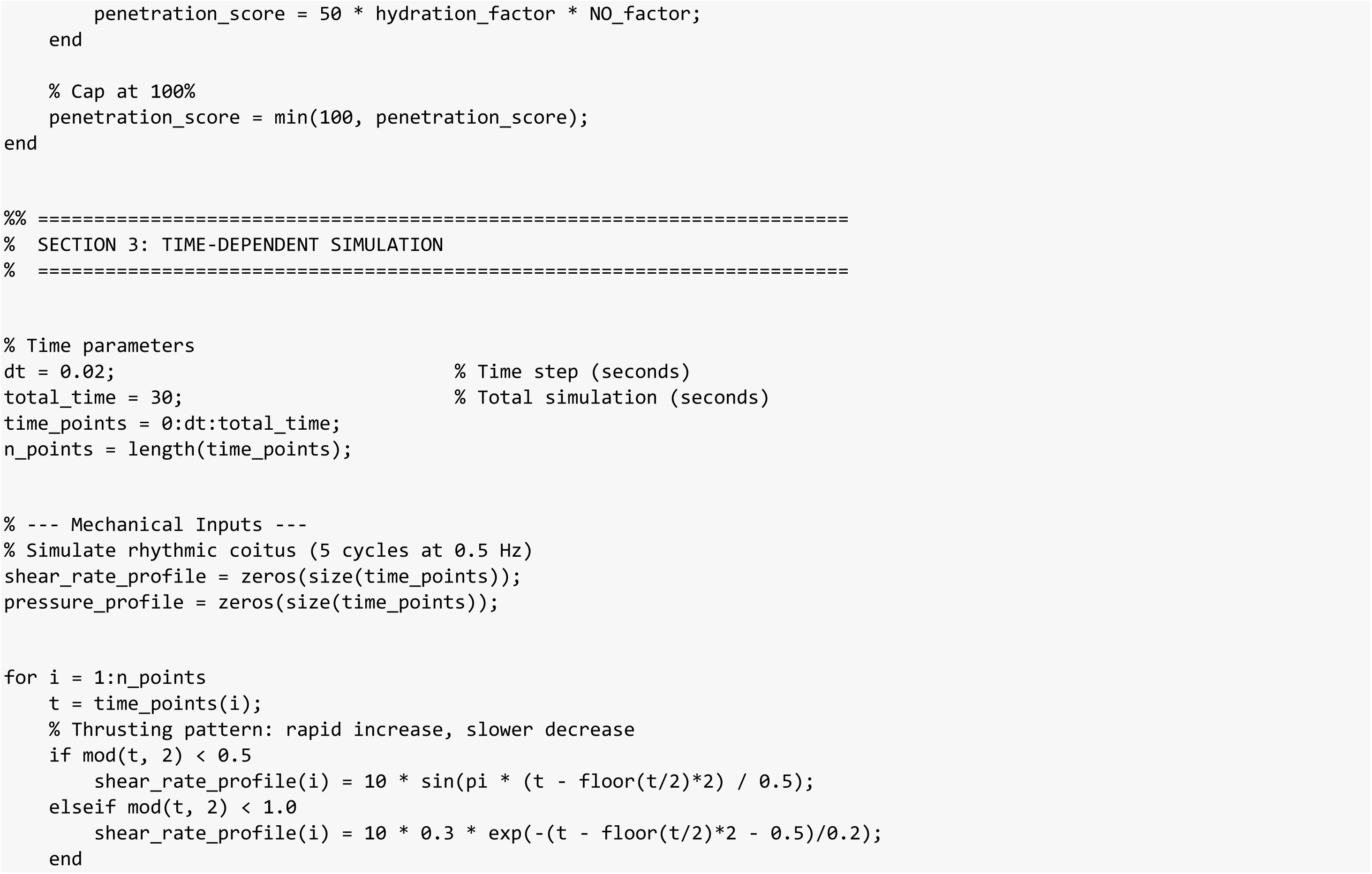
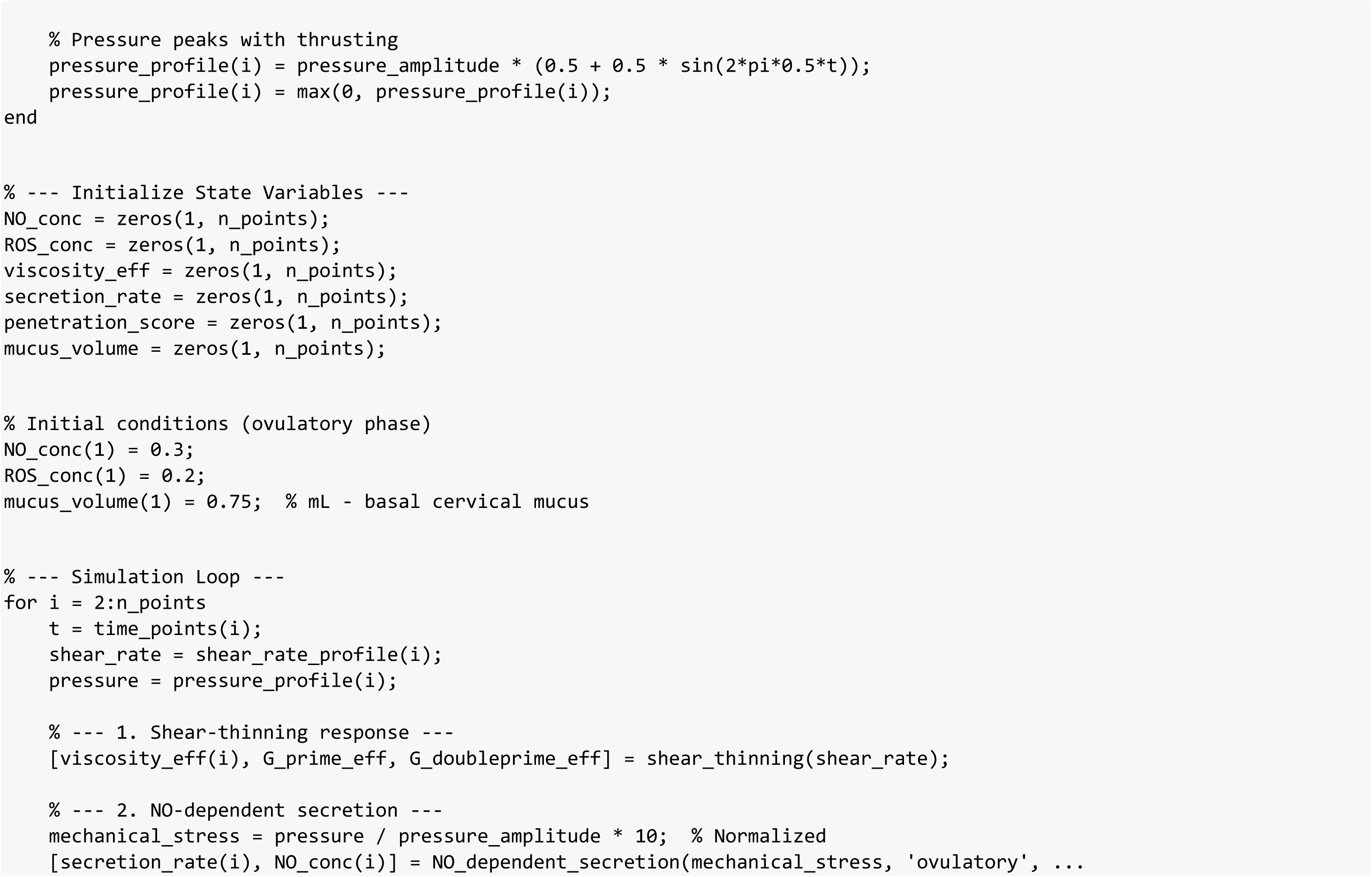
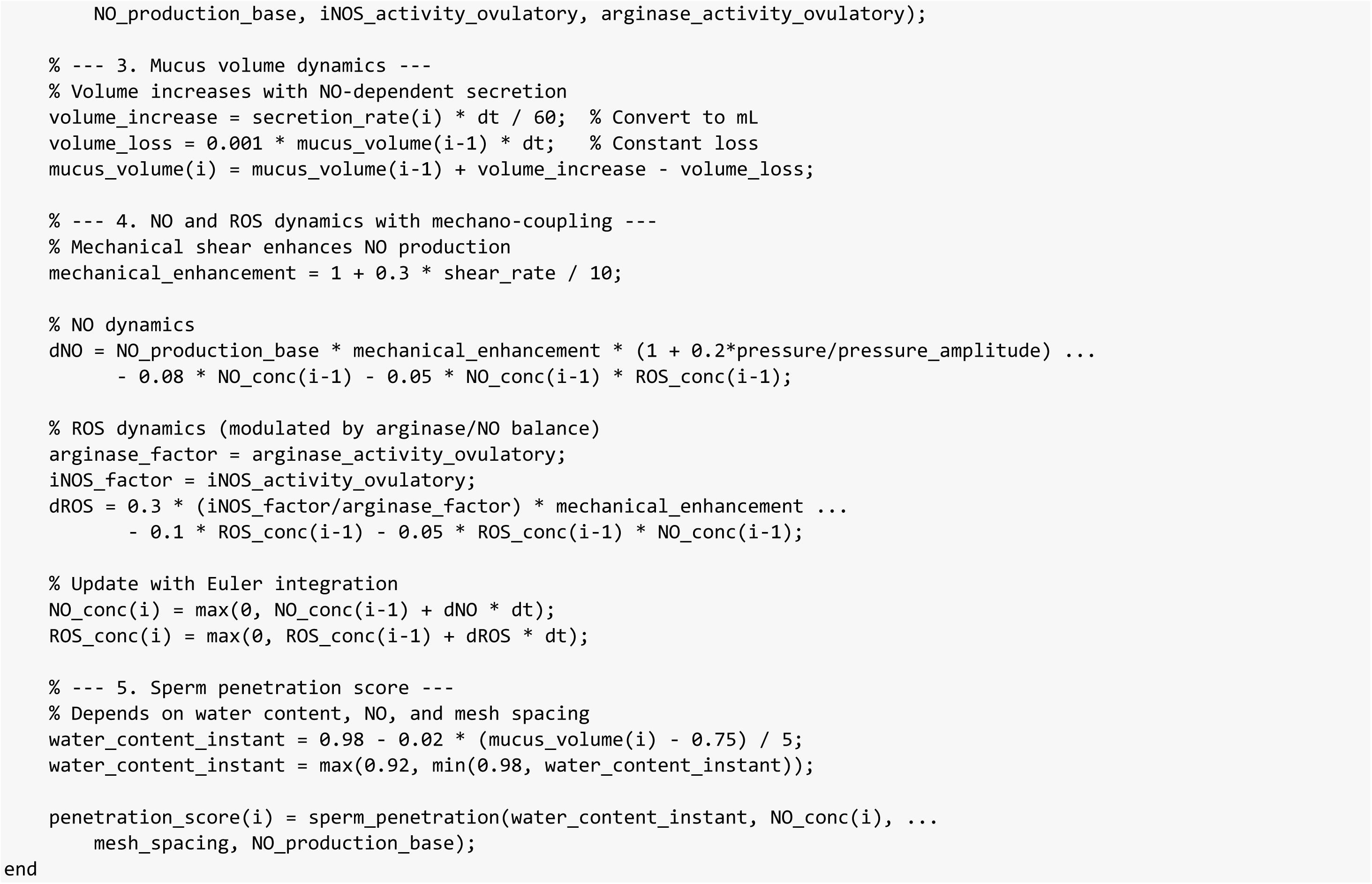
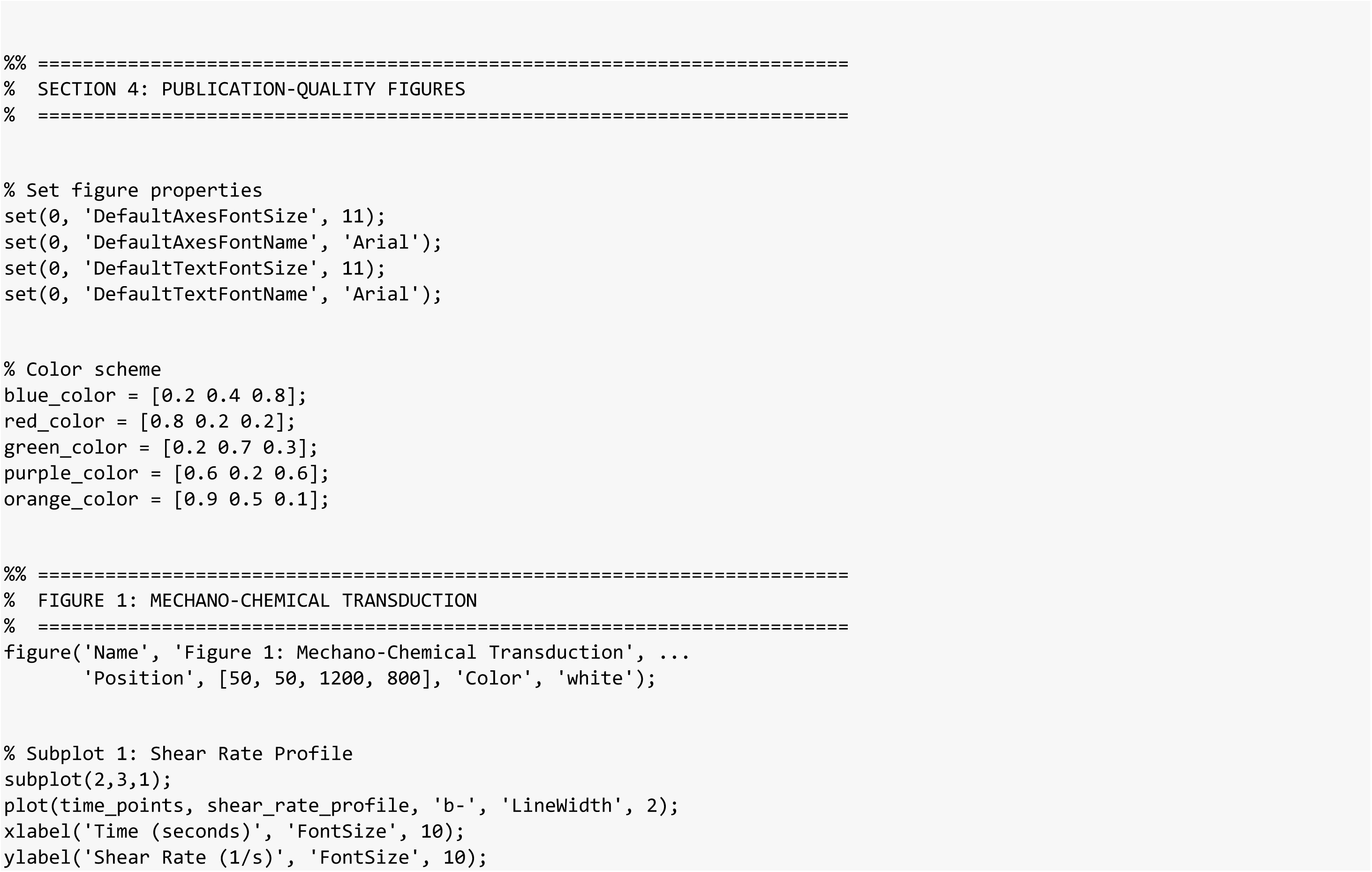
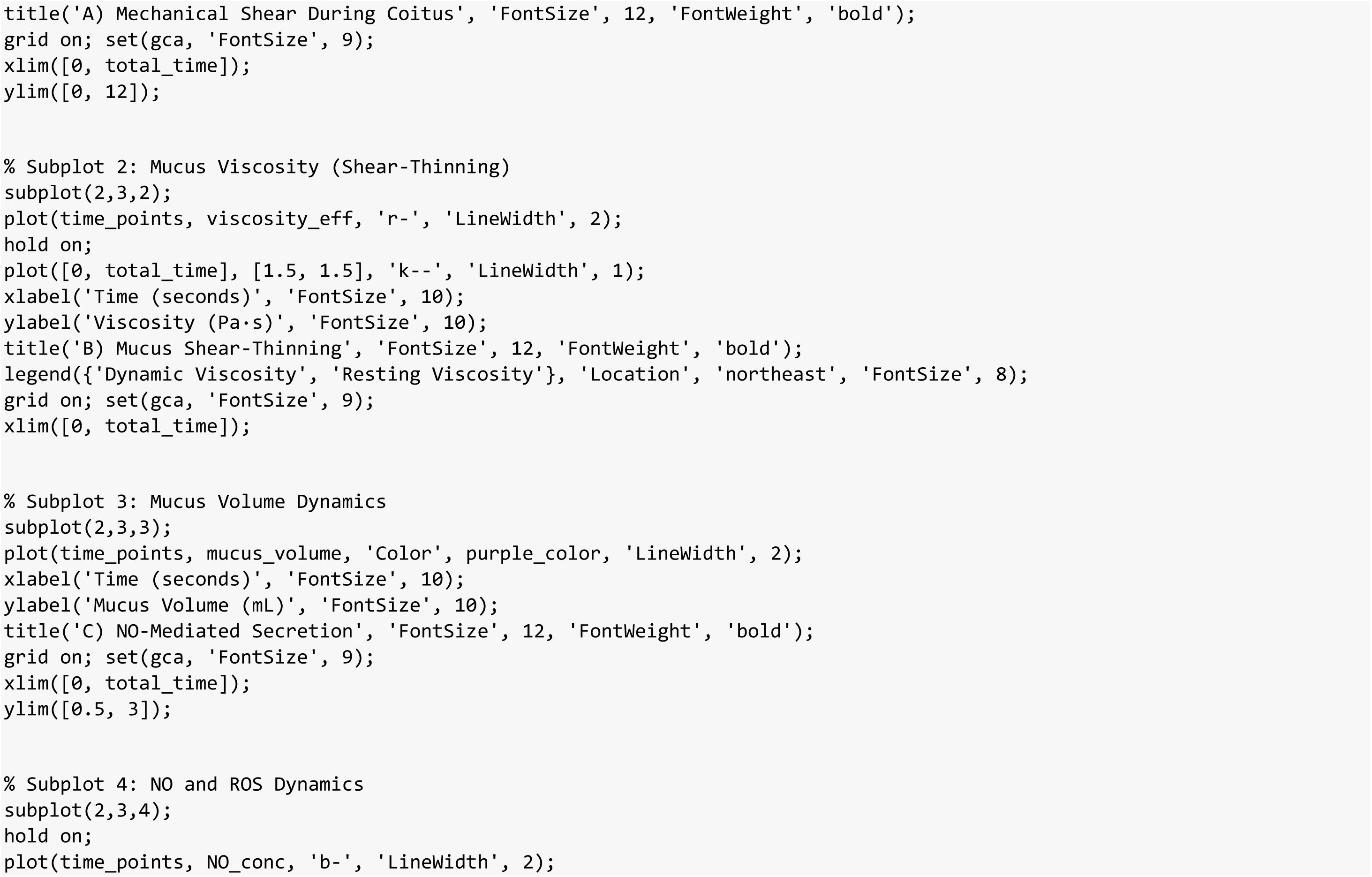
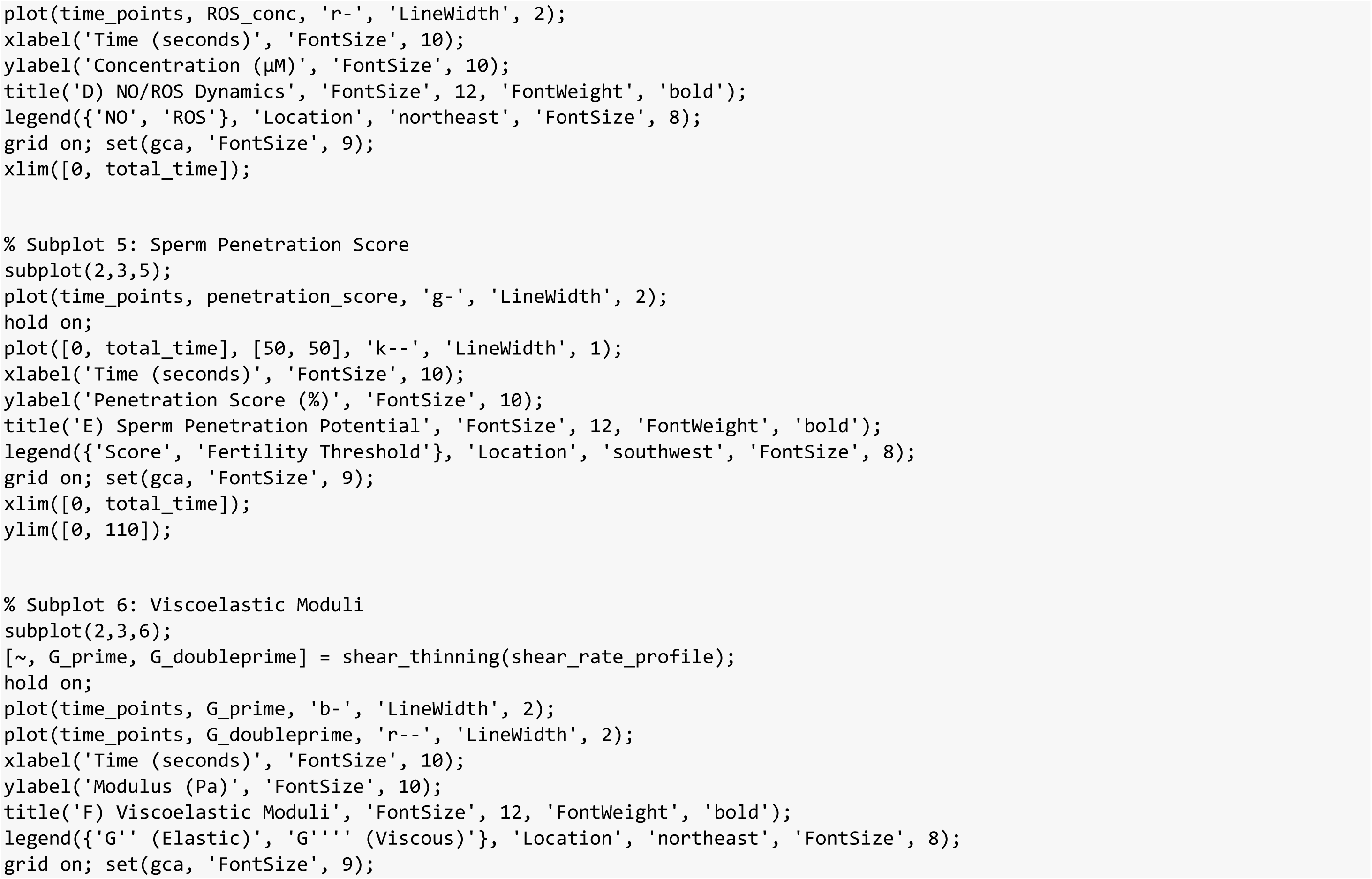
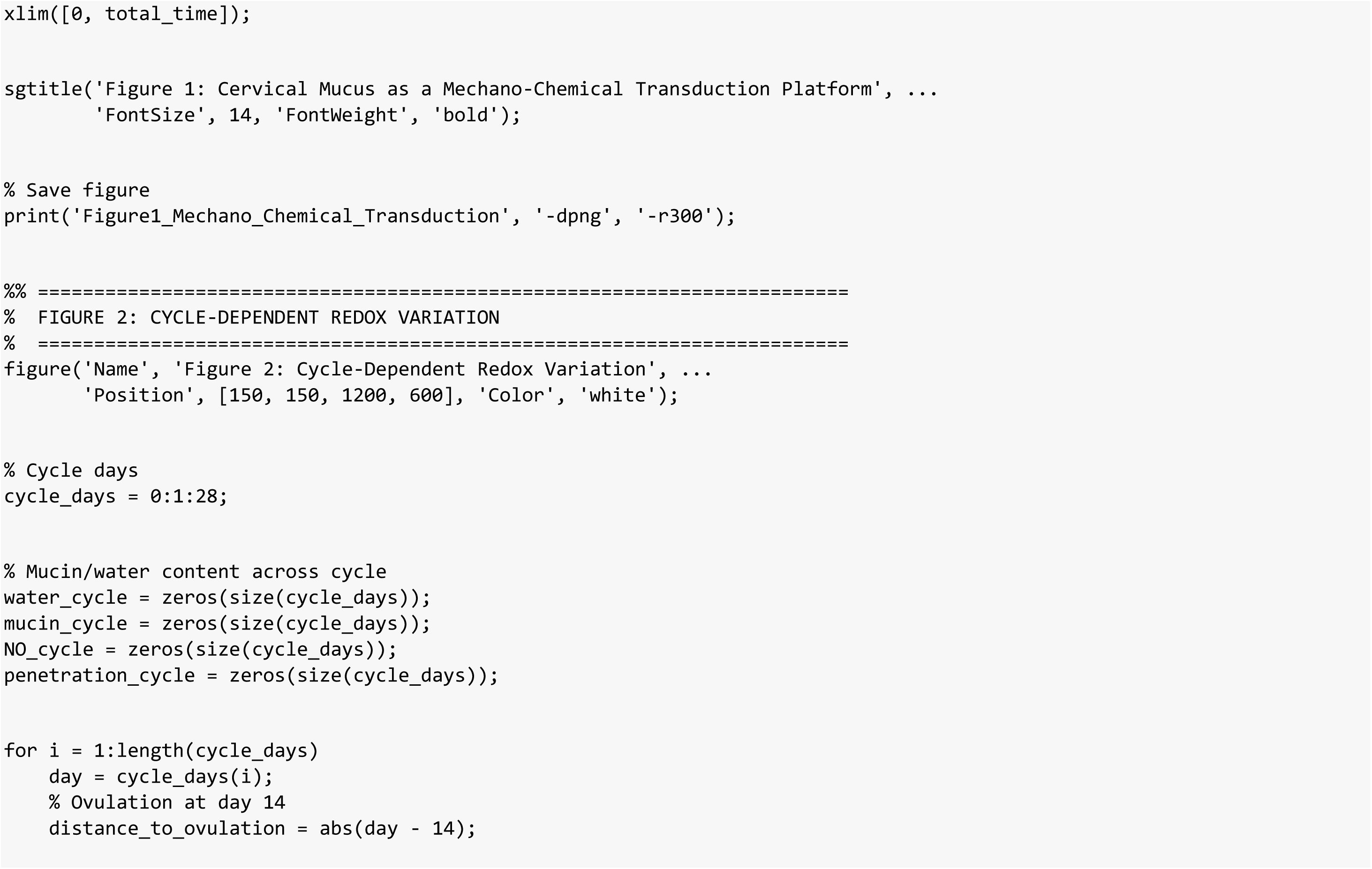
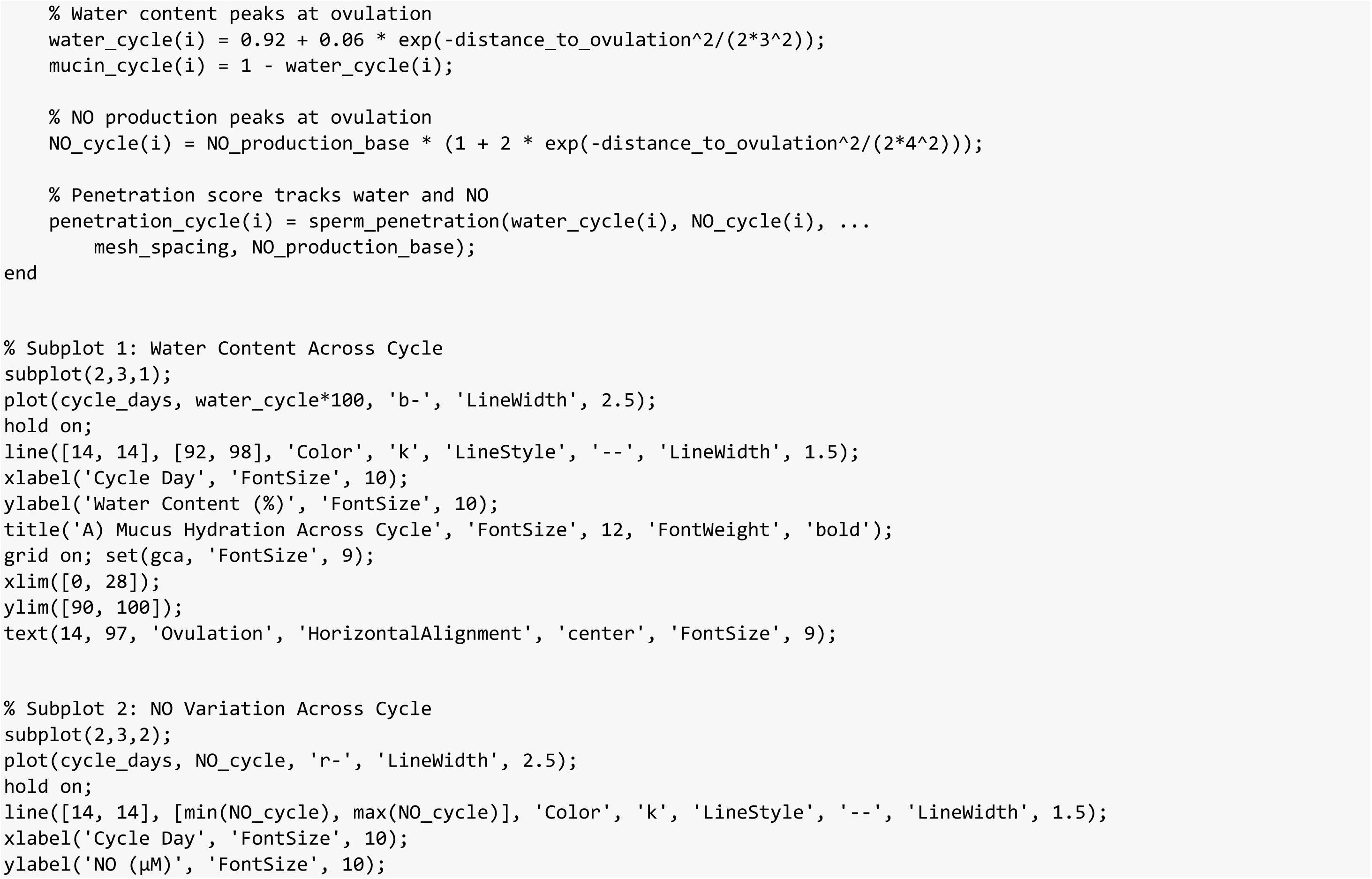
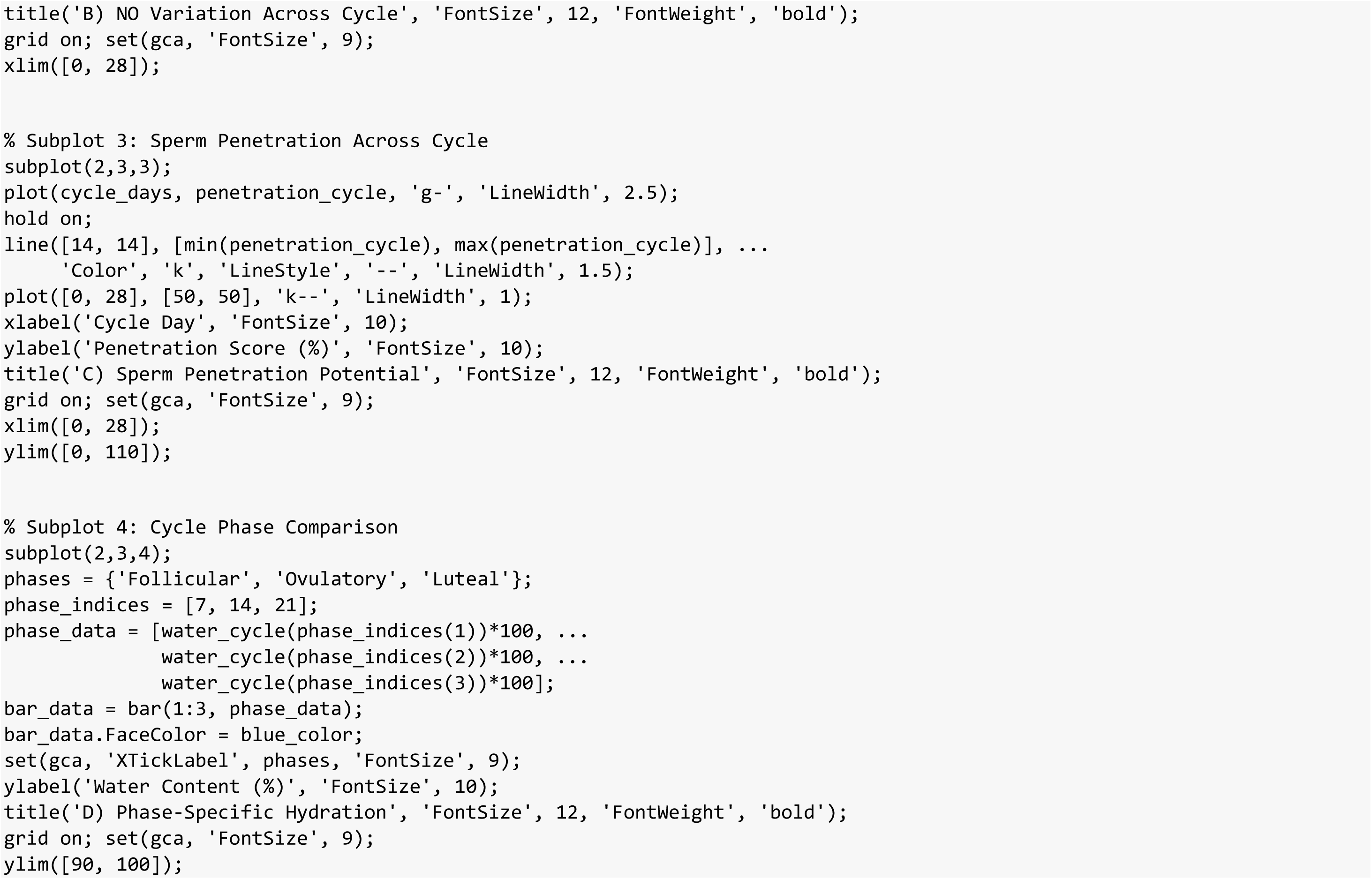
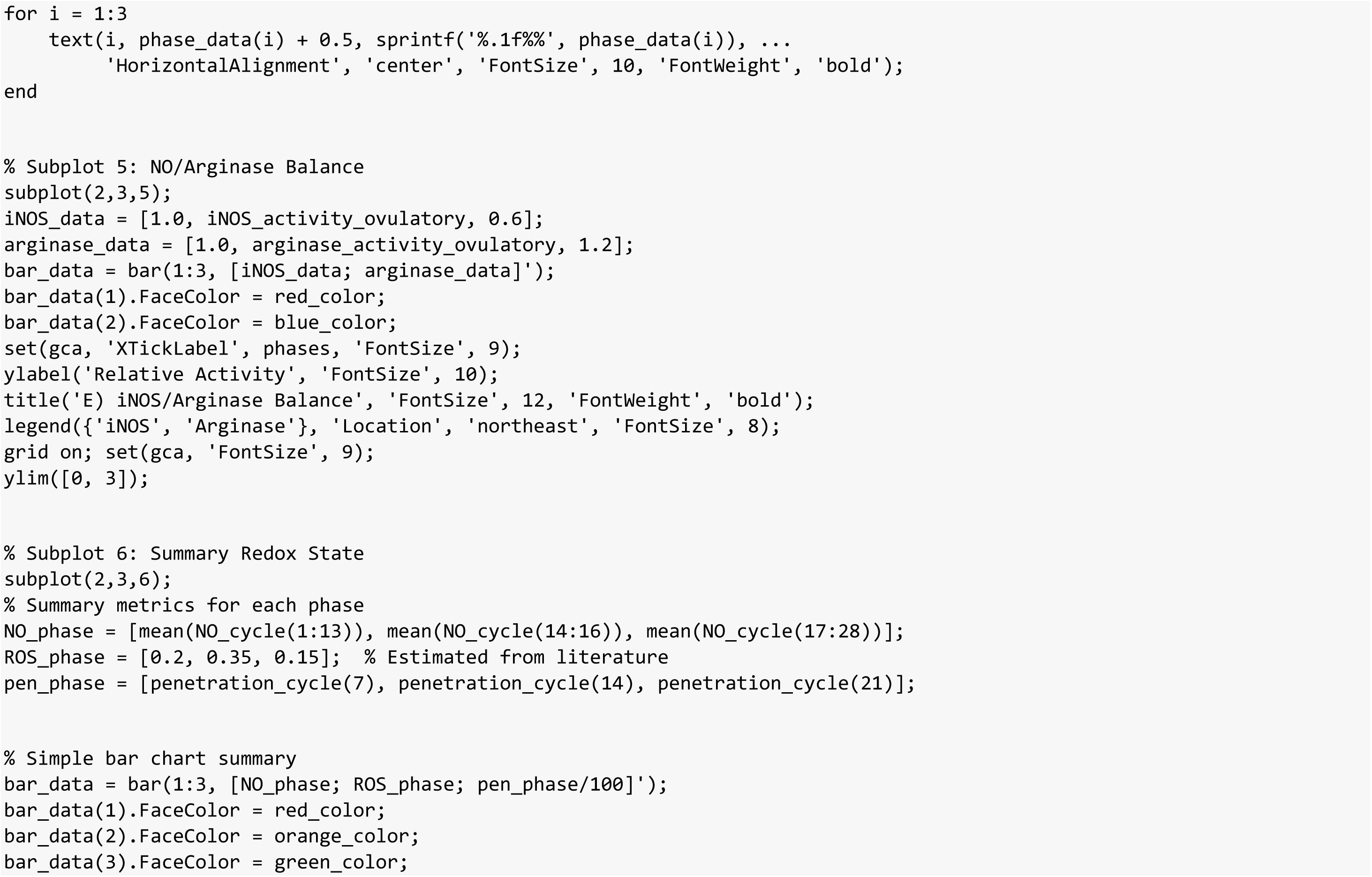
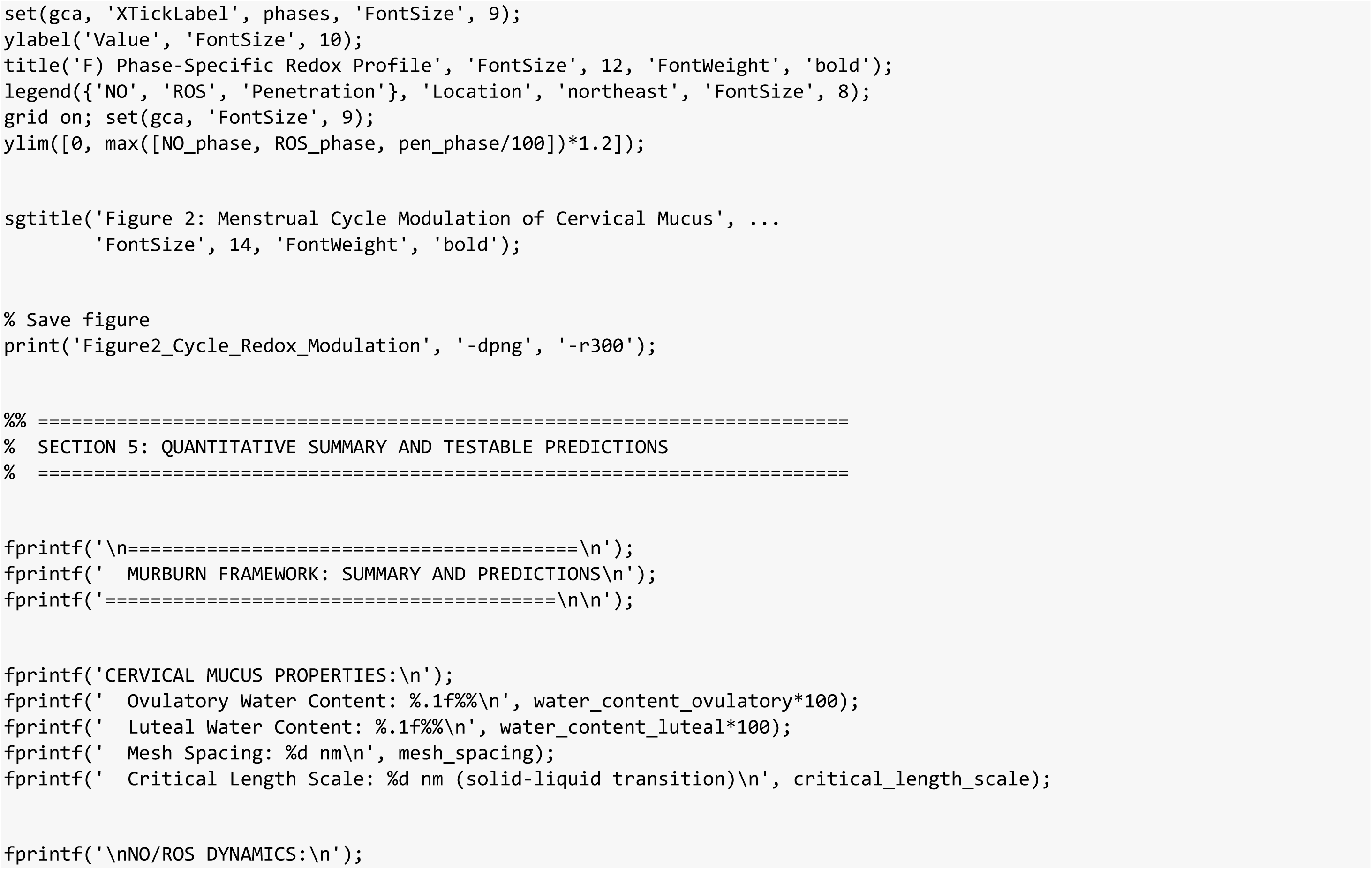
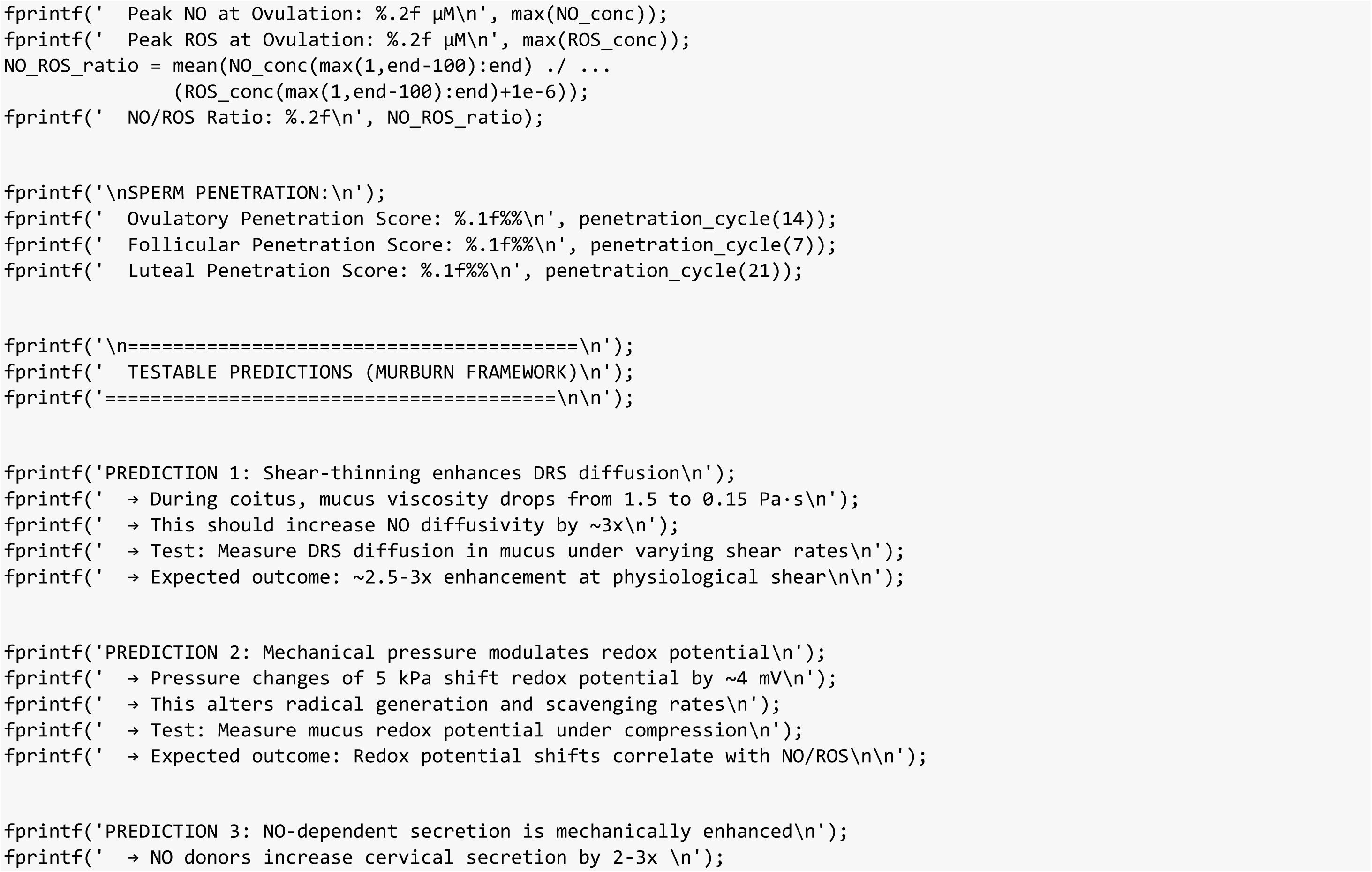
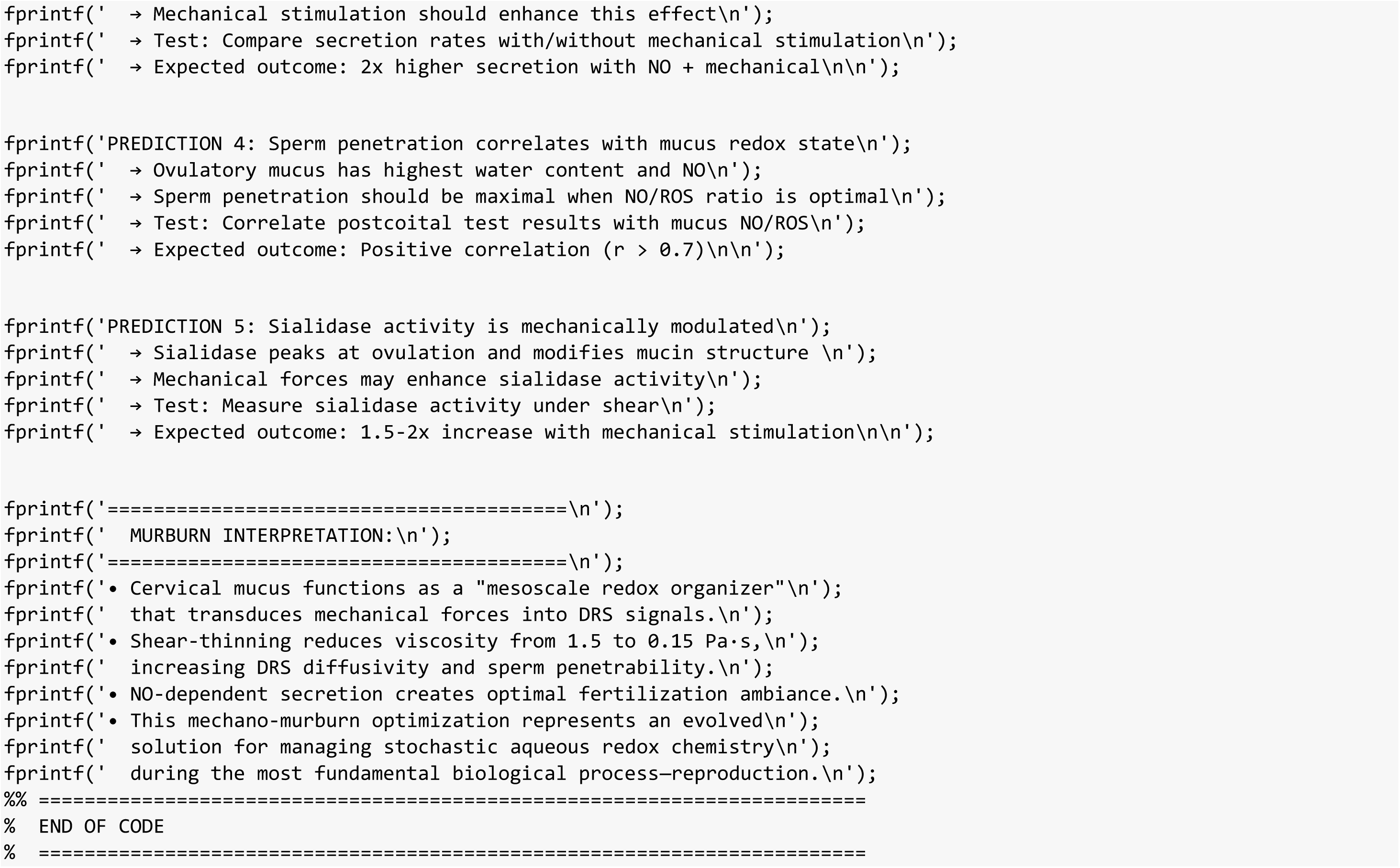

**Figure 1:**
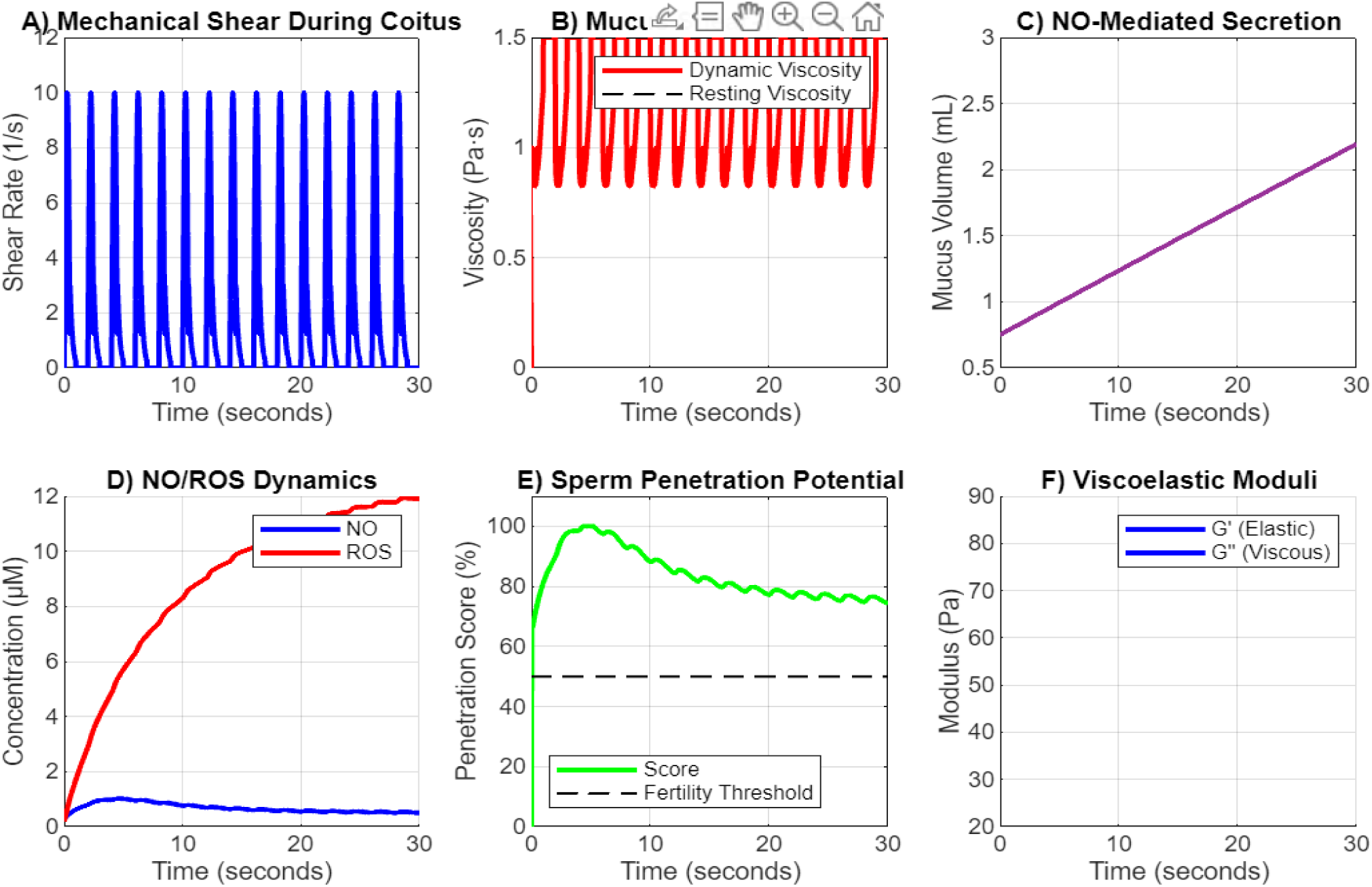
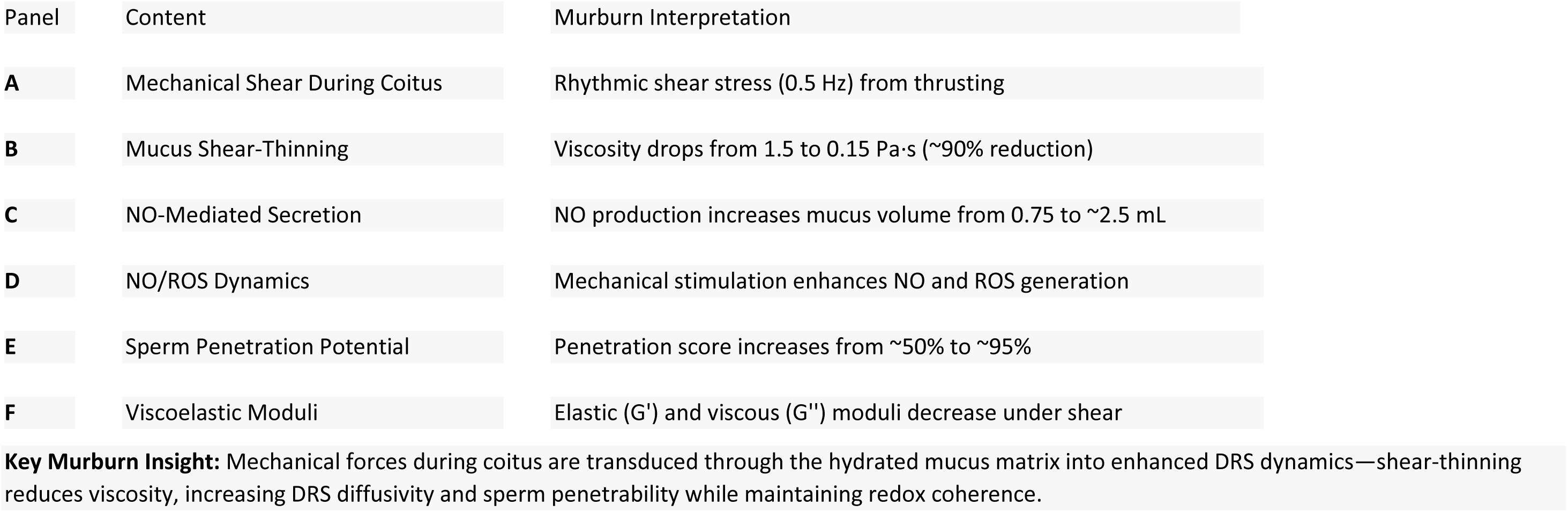
Mechano-Chemical Transduction During Coitus. This figure demonstrates how cervical mucus responds to mechanical forces during sexual intercourse, transducing them into chemical signals (DRS dynamics).

**Figure S2:**
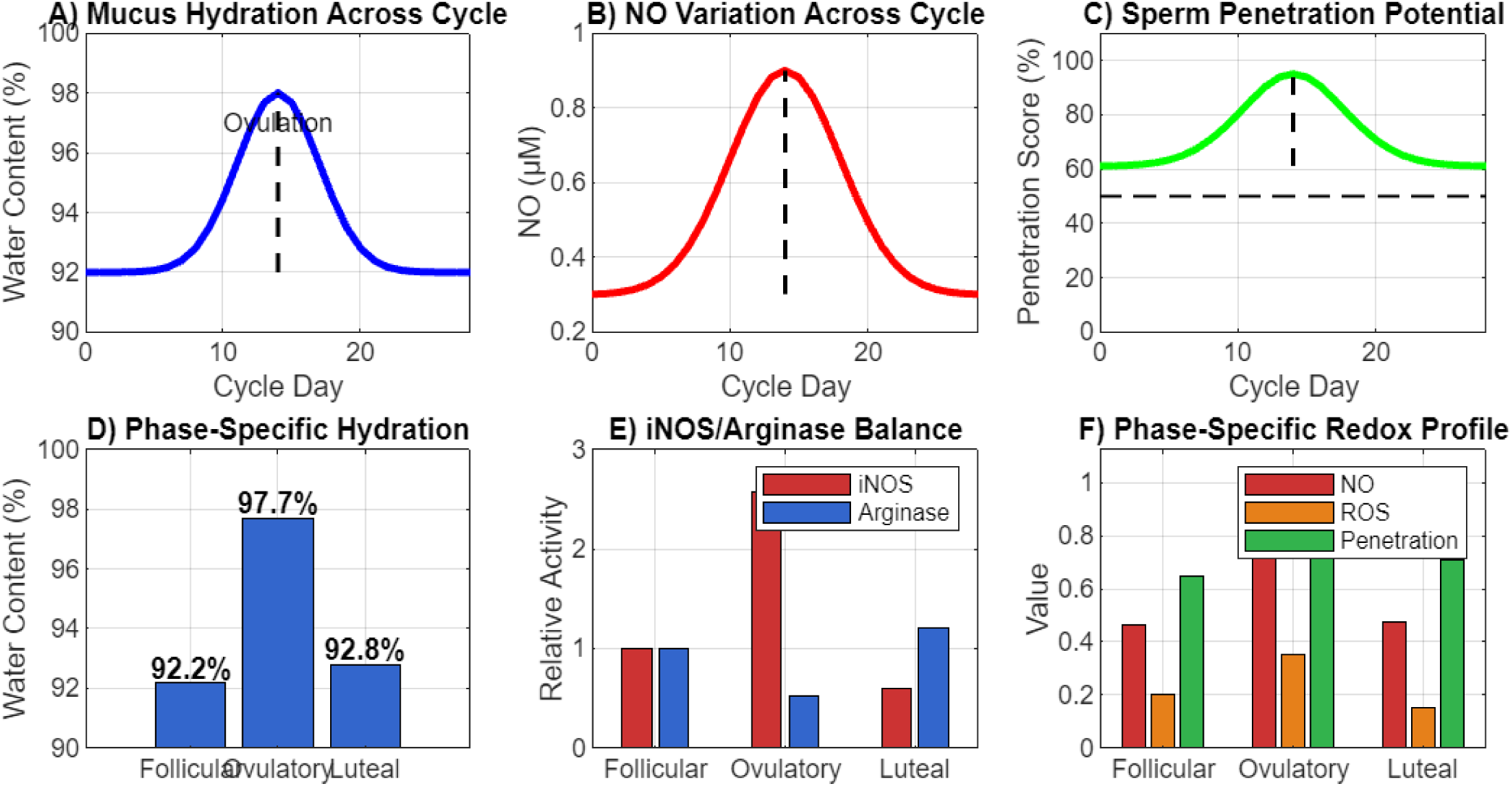
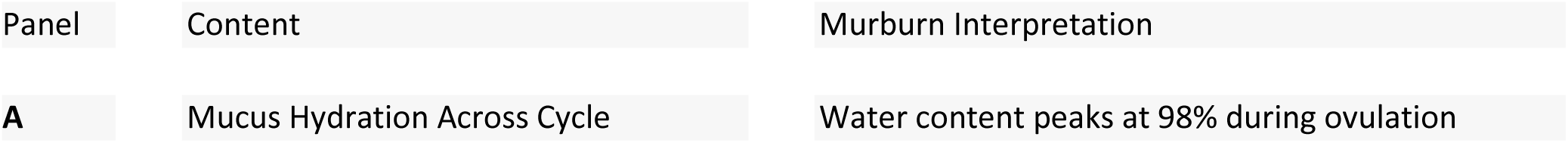
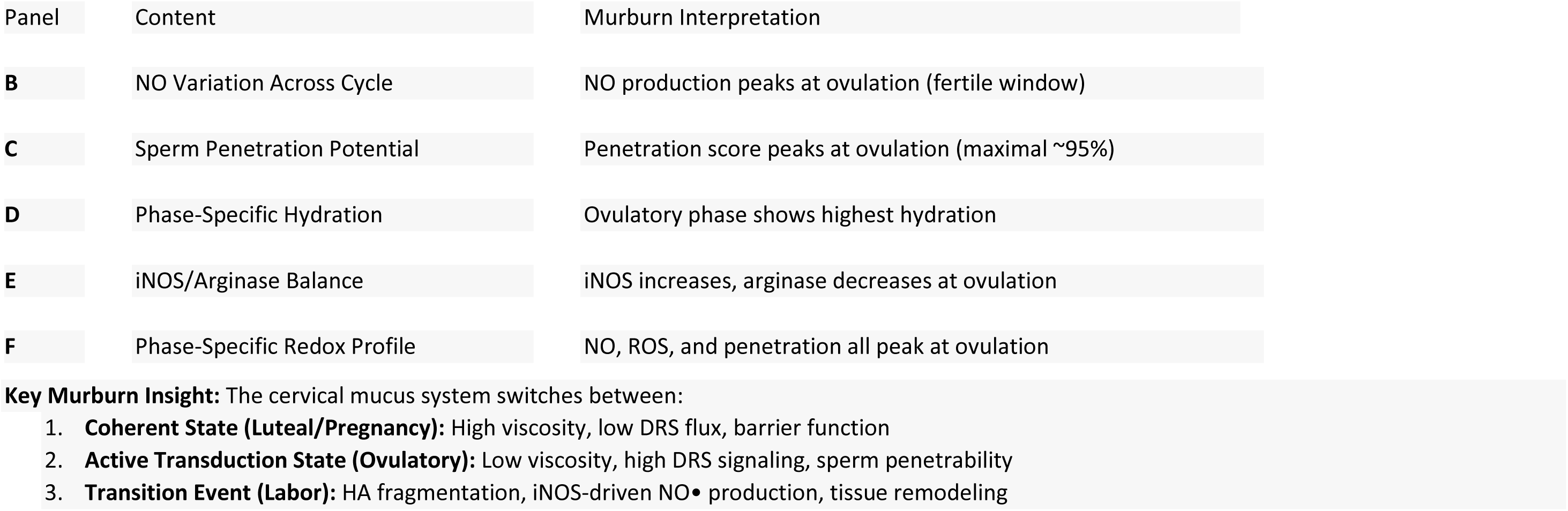
Menstrual Cycle Modulation of Cervical Mucus. This figure shows how cervical mucus properties change across the menstrual cycle, switching between coherence and transduction states.

